# Palynofacies, environments, and climate changes in the Magdalena River Basin

**DOI:** 10.1101/2021.03.07.434303

**Authors:** M. A. Lorente

## Abstract

Four environments (swamp, shallow lake, alluvial flood plain, and lagoon) from the Lower Magdalena River Basin were studied for palynofacies’ quantitative characterization. Each environment has been described based on four criteria: palynomorph assemblage, organic matter concentration, organic matter palynological composition, and organic particle morphology.

**Shallow lakes’** palynological assemblages are dominated by composite and grass pollen. The POM (particulate organic matter) morphology is characterized by a maximum at Φ 5 class (silt), and it has a sphericity histogram with bimodal distribution (peaks at 0.1 and 0.5/0.6). From a composition point of view, POM is mainly opaque amorphous materials. POM concentration is usually lower than 0.1%.

**Swamp** environments palynological assemblages are dominated by grass pollen with a slightly smaller amount of composite pollen. The POM is dominated by finely dispersed amorphous and indeterminate “other” types (organo-mineral gel ?), depending on the oxidation degree. The swamp concentration of organic matter a few centimeters below the water-sediment interface varies between 0.1% and 0.3%. Below that, organic concentration is usually lower than 0.1%.

**Lagoon** assemblages are rich in species and specimens, but assemblages are highly variable. Main components are either finely dispersed amorphous or plant cuticular/epidermal or amorphous homogeneous and heterogeneous or fungal remains. Peat lithology is rich in mangrove pollen, while clay assemblages are dominated by composites, grass, and water plants together with *Botryococcus* algal remains. Lagoon sediments are the richest in POM concentration, with values between 0.13% and 1% (excluding peats). Regarding particle size and shape, in this environment, they show a trend to decrease in grain size from Φ 1 to Φ 2 class (sand) dominated assemblages to Φ 5 to Φ 6 class (silt) dominated assemblages from base to top. Elongated shapes are abundant, with 30% to 50% of particles in the tabloid to elongated tabloid classes.

**Alluvial - fluvial flood basin** samples are often barren in palynomorphs and organic matter. Occasionally present grass pollen and fungal remains. The POM, when present, is mainly of organo-mineral gel type and has a bimodal grain size distribution, with a minor peak at Φ 7 class (v.f.silt) and a major peak at Φ 4 to Φ 2 class (c. silt to f. sand).

Significant changes in quantitative palynofacies occur within the top few meters of the cores, representing the last 1000 yr of sedimentation in the area. These changes are related to shifts in climate, from colder to warmer conditions or from dry to wet periods, most probably linked with E.N.S.O. A short dry and cold period related to the “Little Ice Age” was identified in the Ayapel and Cienaga de El Medio cores.

## INTRODUCTION

This paper is about the characterization of fluvial environments using quantitative palynofacies. The aim was to improve the understanding of the sedimentation and preservation of palynomorphs and palynological organic matter (POM) in recent environments as a tool for paleoenvironmental interpretation. Some previous palynofacies studies in fresh to brackish water, deltaic, and marine environments (Lorente, 1990a; Caratini, 1994; Gastaldo, 1994; Gastaldo et al., 1996; von Waveren & Visscher, 1994; Rull 1995; and Prasad et al. 2007) revealed that the recent assemblages of palynomorphs together with the textural characterization of the palynological organic matter are a useful but complex tool for environmental characterization. Some of the differences observed in recent assemblages show that it is fundamental for the characterization of recent palynofacies to study surface assemblages and the changes that occur to the organic matter within the first few burial meters. Changes are reflected in composition, particle morphology, as well as organic and palynomorph concentration. Some changes in the average preservation of the organic materials within the same environment may be related to minor climate changes that generated drier or wetter conditions during relatively short time intervals.

Preserved palynofacies are the result of a complex set of geological, climatological, and biological conditions. It is possible to identify a group of parameters (POM composition, concentration, particle morphology, and palynomorph assemblages) to characterize palynofacies from the various recent environments.

Studies of palynofacies had been widely used in older sediments with different purposes, from paleoenvironmental interpretation to the evaluation of hydrocarbon’s source rock potential (e.g., Person et al., 2016; Guler et al. 2013; Alaug et al., 2013; Vajda et al. 2013; Makled and Baioumi, 2013; Zooba et al. 2011; Al-Ameri et al., 2009; Ghasemi-Nejad et al., 2009; Martinez et al., 2008; Skupien & Mohamed, 2008; Ruckwied, 2008; Pross et al. 2006; Araujo Carvalho et al., 2006; Al-Saad and Ibrahim, 2005; Bak et al. 2005; Batten and Stead., 2005; Roncaglia, 2004; Vajda, V. 2001; Jaramillo and Oboh-Ikuenobe, 1999; Buchardt and Nielsen, 1991; Lorente, 1986; Tyson 1984, 1987, 1989; 1993; Batten, 1982, 1985).

In this study, samples are either from the surface or shallow cores taken in recent fluvial environments in the Lower Magdalena River Basin. The cores are part of the project ‘Recent palynofacies in the Lower Magdalena River Basin’ at the Hugo de Vries Laboratory, University of Amsterdam, carried out during the nineties. The research results were registered in an internal report (Lorente, unpublished, see attachment 1) but never before published or made publicly available in any other form.

The core and surface samples belong to those taken during the development of the “Proyecto Cuenca Magdalena-Cauca” as part of a cooperation program between the Dutch and the Colombian Governments in the seventies, a multidisciplinary project designed to recover the flooded area of the Lower Magdalena River for agricultural purposes. For this study, a total of 61 sub-samples were taken from those cores and surface samples representative of different river basin environments (Figure 1).

**Figure 1:**
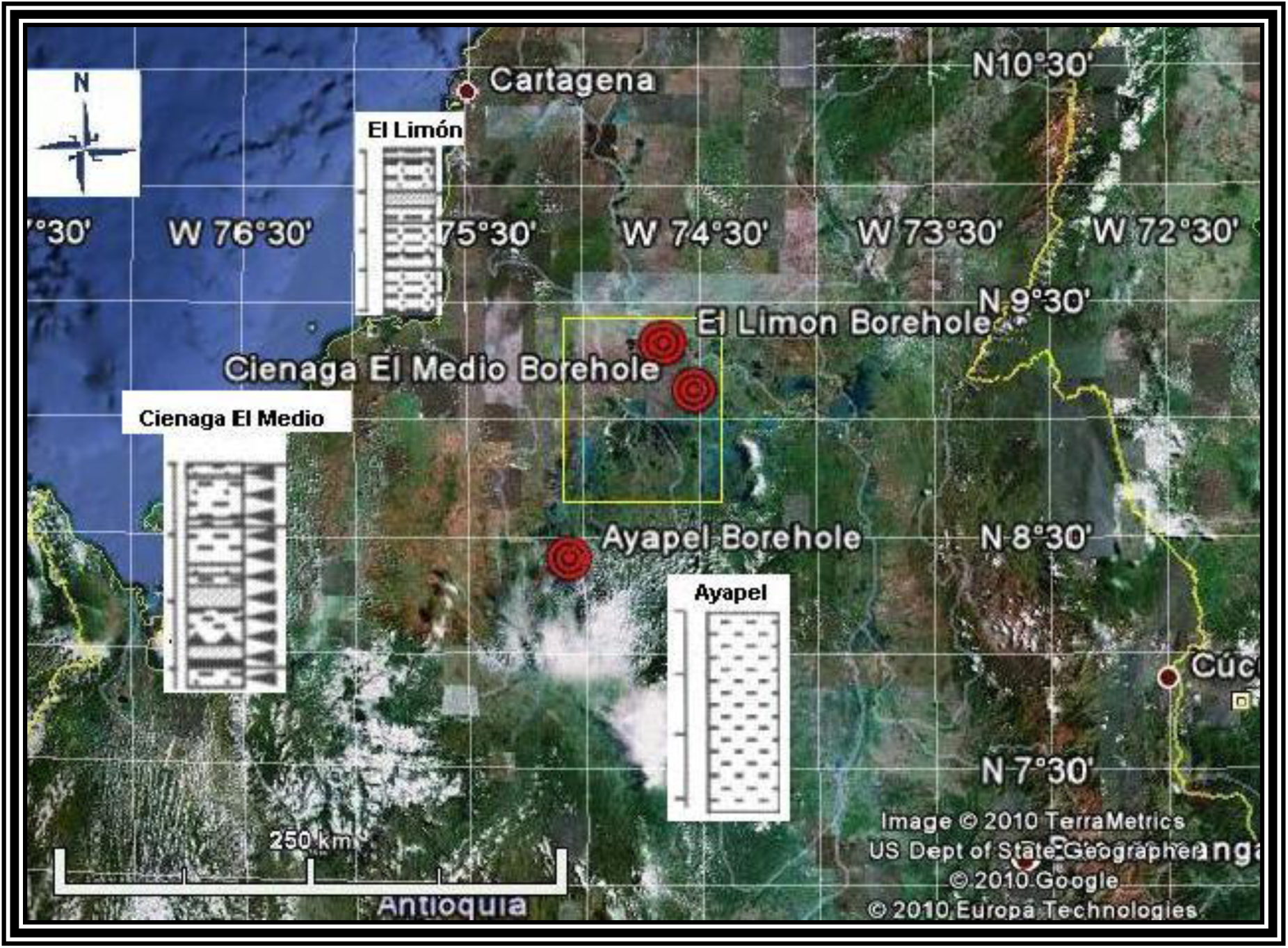
General location of studied samples and sections Lower Magdalena River Basin, Colombia. Red circles show the location of cores, and the yellow square shows the swamp surface sampling area.

### General information

The Magdalena River’s tributary basin is located between 2° and 11° N, and is restricted to equatorial latitude, covering a drainage area of 257.000 km^2^ known as the Magdalena – Cauca Basin. From Barranquilla to Paramo de las Papas, the total river length is 1550 km, with a 6700 m3/sec flow rate. The mean rainfall for the drainage basin is 2050 mm, from a maximum of 5000 mm to a minimum of 800 mm.

According to Restrepo and Kjerfve, 2000, the area between Rio Cauca and Caño San Jorge, where the swamps and shallow lakes cores used in this study were taken, is characterized by moderate rainfall, which averages 1200 mm/y (region C in their paper). Those authors also reported two wet and two dry seasons during the year. December to March and June to September are low rainfall periods, while March to May and October to November are high rainfall periods.

The swampy depression (“depresion cenagosa” or “zona lacustrina’) has an area of 207.700 Km^2^. The statistics show that the ‘flooded area” covers 16% of the total area while the “dry area “covers about 84%.

According to the data collected for the Dutch-Colombo Project (H.I.M.A.T., 1977), the flooded area has the following regime:

6 - 12 months inundated: 19%
3 - 6 months inundated: 30%
1- 3 months inundated: 14%
> 1 month inundated: 21%

The average annual sedimentation rate in the swampy depression during the last 1500 years was 2.9 mm, but there are significant variations related to local conditions (H.I.M.A.T., 1977) like the sedimentation rates in:

Palmitas: 2.7 mm
Sucre: 3.3 mm
Boquillas: 4.0 mm
Mompos: 1.7 mm

### Vegetation in the Lower Magdalena River Basin

The flora in the Magdalena River’s lower course was described by Cuatrecasas (1958) and Wijmstra (1967). The main assemblages described from the river-plain are grassy plains with scattered groups of trees or open vegetation, gallery forest along the riversides, or dense vegetation and vegetation belts around the swamps (“cienagas”).

The open vegetation typically occurs under extreme climatic conditions, e.g., rainfall concentrated in few months every year with long dry periods (Wijmstra, op. cita). It consists of two different types depending on vegetation density:

- The grassy plains, with grasses as *Andropogon bicornis*, *Paspalum millegranum*, *Panicum vulgaris*, etc.; palms and trees as *Mauritia minor*, *Hirtella elongata*, *Bowdichia* sp., *Byrsonima crassifolia*, *Curatella americana*, *Palicourea rigida,* and *Cecropia* sp.
- The “savanna woodland”, with all the plants mentioned above plus *Croton glabellum* and *Ficus elliptica*.
- The gallery forest along the riversides is usually characterized by species of trees as *Inga calliandra*, *Abizzia* sp., *Heliocarpus* sp., *Salix* sp., *Cecropia* sp., or *Celtis* sp. Other trees present in the association belong to the Bombacaceae, Anacardiaceae, Polygalaceae, *Hedyosmum* sp., *Miconia* sp. and various species of Palmae. Gramineae and Cyperaceae are present in the places usually inundated by the river.

The vegetation around the cienagas is composed of the following belts:

- Innerbelt, where the water is deeper, with *Nymphaea* sp., *Limnanthemum* sp., *Trapa* sp. and *Cabomba* sp.
- First vegetation belt landwards is partly continuously inundated and partly temporarily inundated, with *Eichhornia crassipens* and *Pistia stratiotes*. Also present *Salvinia* sp, *Marsillia* sp., *Ludvigia* sp. (Jussiaea), and *Polygonum* sp.
- Second vegetation belt landwards is inundated during most of the year, but gets drier gradually. Grasses and sedges characterize it. Grasses may consist of one species like *Paspalum* sp., or several species like *Eriochloa* sp., *Panicum* sp., *Leptochloa* sp., *Eragrostis* sp., *Echinochloa* sp. and Cyperaceae like *Cyperus* sp., *Ligularis* sp. and *Imperata contracta*.
- Third vegetation belt landwards, consists of a bush- and tree-vegetation.
- Fourth vegetation belt landwards, is tree-dominated where the groundwater level is at its lowest. Present in the association are *Cecropia* sp., *Ficus* sp., *Celtis* sp., *Psychotria* sp., *Zygia* sp., *Inga* sp. and other Mimosaceae. Plants like *Salix humboldtium*, *Alnus* sp. and *Myrica* sp. only occur in the upper course of the Magdalena River, in the Andes, where they grow between 2000 m and 3500 m.

## MATERIALS AND METHODS

The original sampled materials were cores and surface samples stored in good conditions at INGEOMINAS core storage in Bogotá, Colombia.

All project sampling and sample processing were devised systematically and as quantitative as possible to ensure quality semi-quantitative to quantitative results.

For each sample, always 10 gr. of rock were processed following the standard palynological method (HF, HCl, and ZnBr2), subjected to no oxidation or sieving unless otherwise indicated in the text, analyzed for palynomorphs, palynological organic matter composition, and textural characterization. For measurements of the POM particles, the method followed was similar to the one described in Lorente (1986, 1990a and 1990b) for quantitative studies of palynofacies. The organic matter classification used follows the guidelines of the “Amsterdam Palynological Organic Matter Classification” according to the simplification of categories agreed on the 3 ^rd^ Workshop on Palynological Organic Matter Classification, Cocodrie, Louisiana, 1993 (Figura 2).

**Figure 2:**
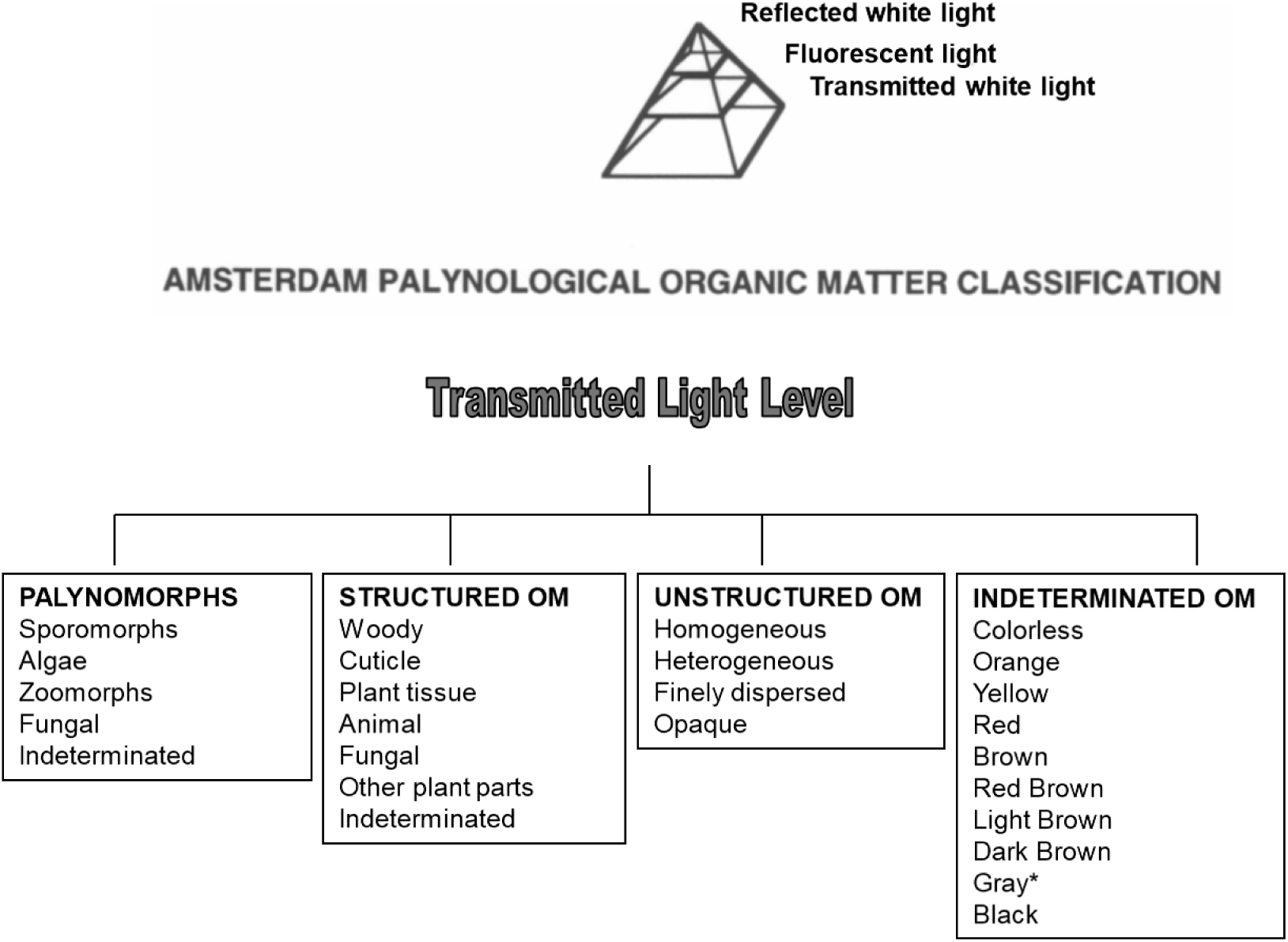
Amsterdam Palynological Organic Matter Classification, a three-level system for morphological classification of POM. It was originally approved at the 1st Workshop on Palynological Organic Matter Classification in Amsterdam, 1991. In this work, we adopted the simplification agreed on in the 3erd Workshop held in Cocodrie., Louisiana in 1993 (Gastaldo et al. 1996) for the transmitted light level. * Also referred to as organo-mineral gel type in this paper.

The morphological characterization of the organic particles’ was done with the O.M.A.S. system developed at the Hugo de Vries Laboratory. The hardware comprised a microcomputer, a video camera, a microscope with transmitted white light, and white and fluorescent incident light. The software for image analysis was commercial. The calculations were performed with a series of excel macros developed at the Laboratory.

The measurements taken for each particle were:

- AREA: calculated as the sum of pixels in an object.
- PERIMETER: length of the curve following the center of the particle contour pixels
- P2A: square contour length divided by 4 Pl times the area of the object. The minimum value is 1.0 for a circle; any other figure has a higher value.

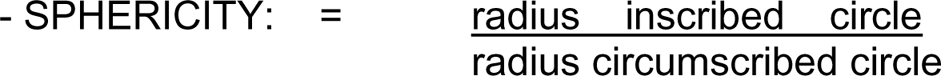

Maximum value equals 1 for a circle; any other figure has a lower value.

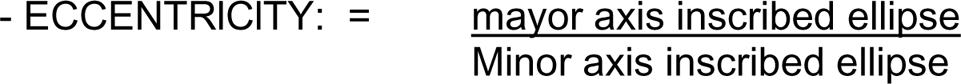

The value for a circle is 1. Any other figure has a value higher than 1.

The following parameters were calculated with the above-described measurements: area, size, sphericity, and relative elongation using the methods described in Lorente 1986, 1990 a & b. The results were plotted in the form of histograms.

Regarding the grain size distribution, two different types of histograms were plotted:

- Sum of areas: grain size distribution in which the total area covered for each size class was related to the total area covered by particles in the palynological slide.
- Number of particles: For each size class, the number of particles was counted and related with the total number of particles measured in the palynological slide.

The palynomorph assemblage was quantified in terms of the absolute amount of specimens per gram of rock. The pollen and spores were identified up to the level of species whenever possible.

## ANALYSIS AND RESULTS

### Fresh Water Lakes and Swamps

A total of twenty-two samples from two cores and five surface samples were studied from these environments. The cores were taken at the Ayapel Lake and the Cienaga El Medio swamp, both permanent freshwater bodies.

The surface samples were taken from 5 swamps: Pimiento, Coyongal, Los Limones, Punta de Blanco, and El Medio, all of them located between 8° 40’N - 9° 10’ N and 74°95’ W - 74°25’W (Figure 1).

#### Ayapel Lake

##### General information

The Ayapel Lake is located very close to the right bank of San Jorge River, between San Jorge River and Cauca River in the vicinity of the localities of “Ayapel” and “EI Cedro” (Figure 1). Numerous water courses and creeks drain the area, among them Caño Muñoz, Caño Cedro, Caño Aguas Claras, Quebrada Escobillas and Ouebrada Ouebradona.

The lake is in the Palmitas area, which has an average sedimentation rate of 2.7 mm/year, according to H.I.M.A.T. (1977). It has an irregular shape with the main axis of about 25 Km length and a total area of water coverage of 250 km^2^.

##### Lithology and sample description

A core of about 3 m length was taken at 75°03’ W - 8°24’N (Figure 1). The lithology is homogeneous, with some intervals richer in plant and root remains. From the core, a total of 11 samples were taken and studied (Figure 3a).

**Figure 3a:**
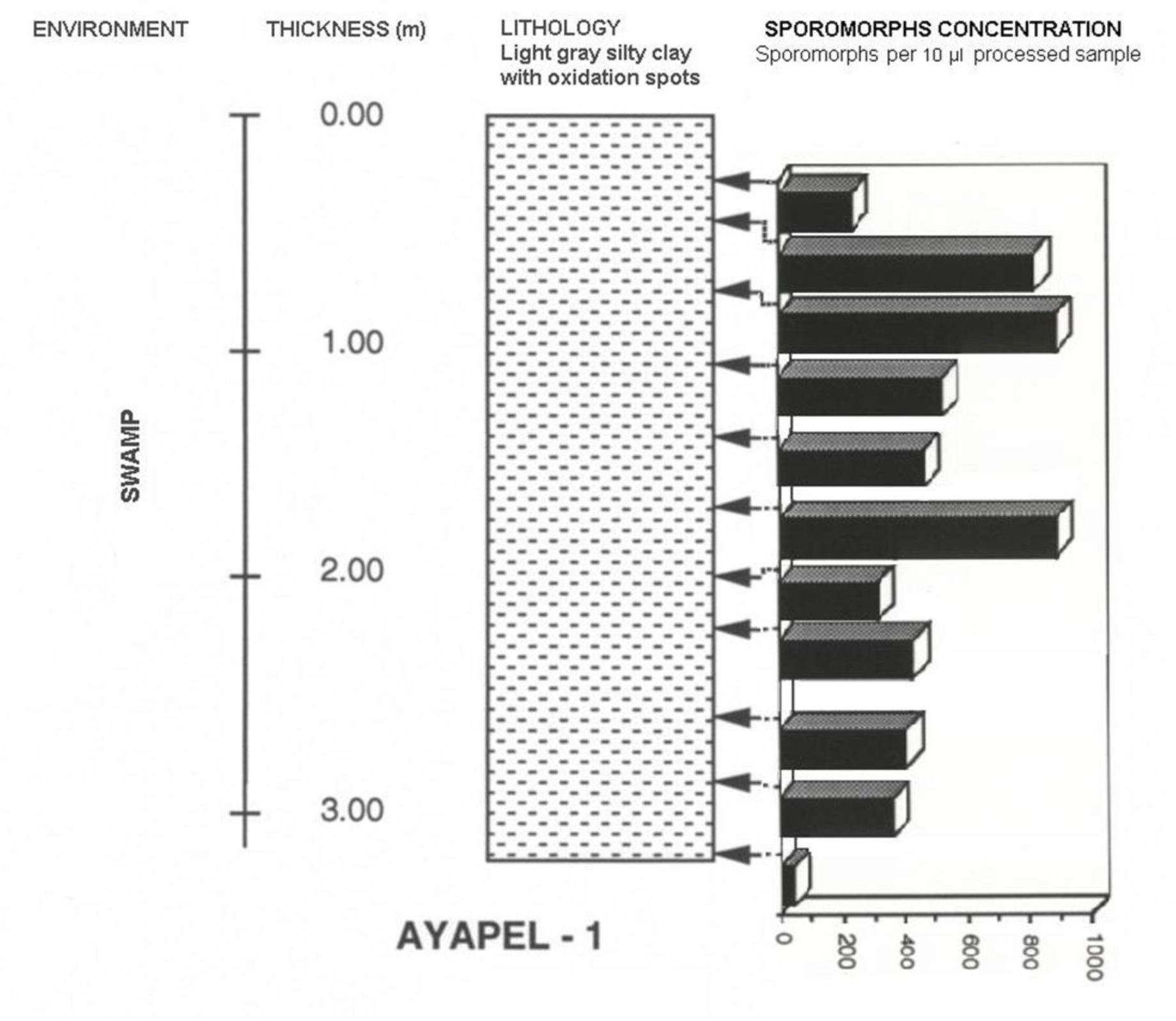
Ayapel Core lithology, sample position, and sporomorph concentration. Concentration: Number of Sporomorphs per 10 μl of the palynologically processed sample.

##### Palynology

Ayapel Lake core samples show variations in the sporomorph concentration (Figure 3a). The flora is richer in specimens from 0.45 m to 1.65 m depth than from 1.95 m to 3.20 m depth. The relatively low value at 0.25 m is maybe due to the diluting effect of abundant organic matter.

The flora of the Ayapel Lake core from base to top can be described as follows (see Figure 3b):

- 2.25 m to 0.25 m, the flora is dominated by composite and grass pollen.

**Figure 3b:**
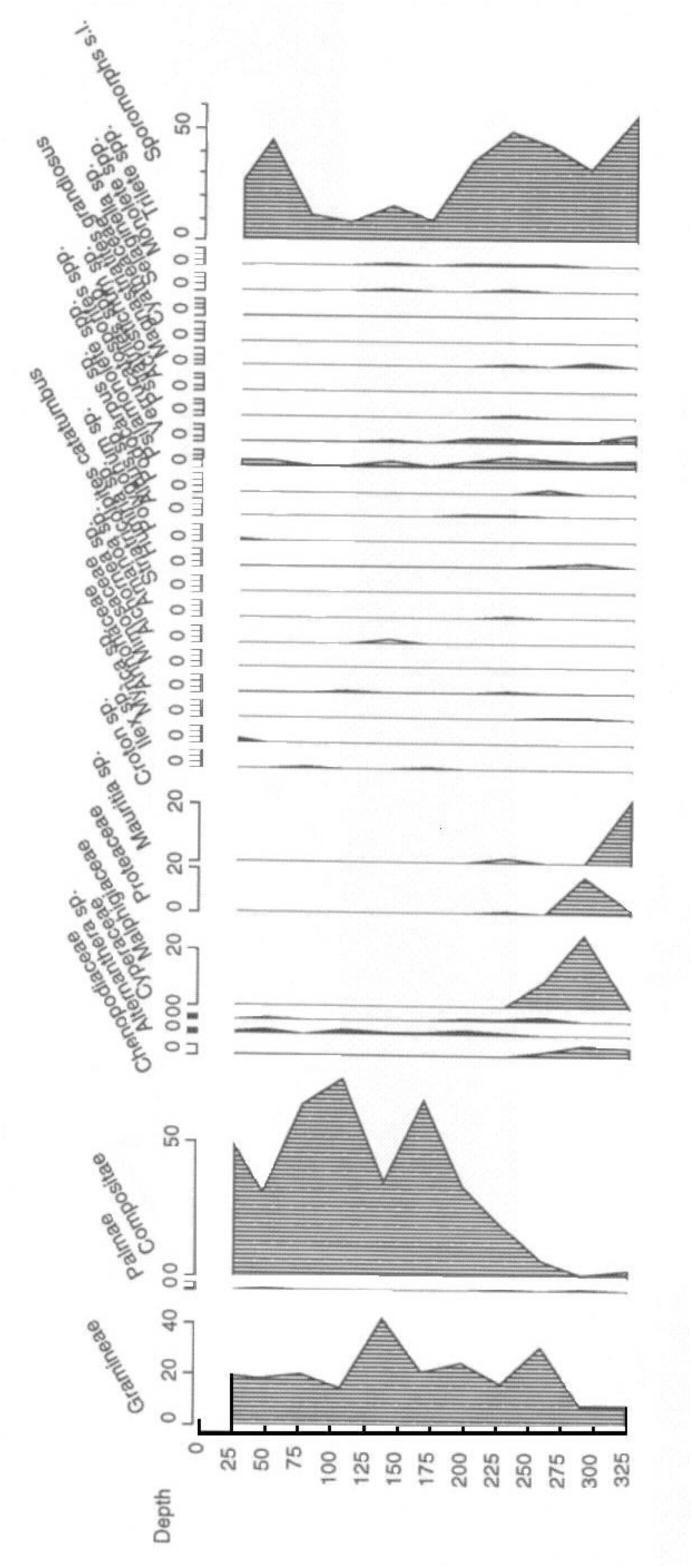
Ayapel Section sporomorph’s diagram.

Pollen from floating/water plants is present throughout the interval but in minor concentrations, with a general increase of sporomorphs abundance from 1.65 m upwards.

In general, sporomorphs tend to be more abundant than fungal remains. From 1.65 m upwards, it increases the number of fungal remains and freshwater algae (including *Botryococcus* sp.).

- 2.85 m - 2.55m, grass pollen is dominant over composite pollen, while pollen from freshwater/floating plants and palms is present in significant amounts.
- 3.20 m, the core deepest sample is poor in sporomorphs.

##### POM facies

The organic matter concentration is, in general, low with values below 0.1% (Figure 3a).

The observed composition may be summarized as follows:

- At 0.25 m.depth, the assemblage is characterized by very abundant light-colored plant tissues as epidermal/cuticular material, and other plant remains. Present but in lower concentrations are opaque amorphous POM, palynomorphs, animal, fungal, and woody remains. The finely dispersed organic matter is very abundant.
- At 0.45 m depth, the assemblage is characterized by abundant palynomacerals, including amorphous heterogeneous and amorphous finely dispersed POM with minor amounts of amorphous opaque POM, light-colored plant tissues, and palynomorphs.
- At 0.75m depth, the assemblage comprises very abundant amorphous homogeneous POM, amorphous opaque POM, and fungal remains.
- From 3,20 m to 1,05 m (3.20 m is very poor) depth, the assemblages are dominated by amorphous opaque POM. Palynomorphs, woody, other plant tissues, fungal remains, and other amorphous POM are present but in very low concentrations.

The shape of particles is variable, although grain size (particle number), sphericity, and elongation histograms are always skewed to the left, except for the most superficial sample (Figure 3c).

**Figure 3c:**
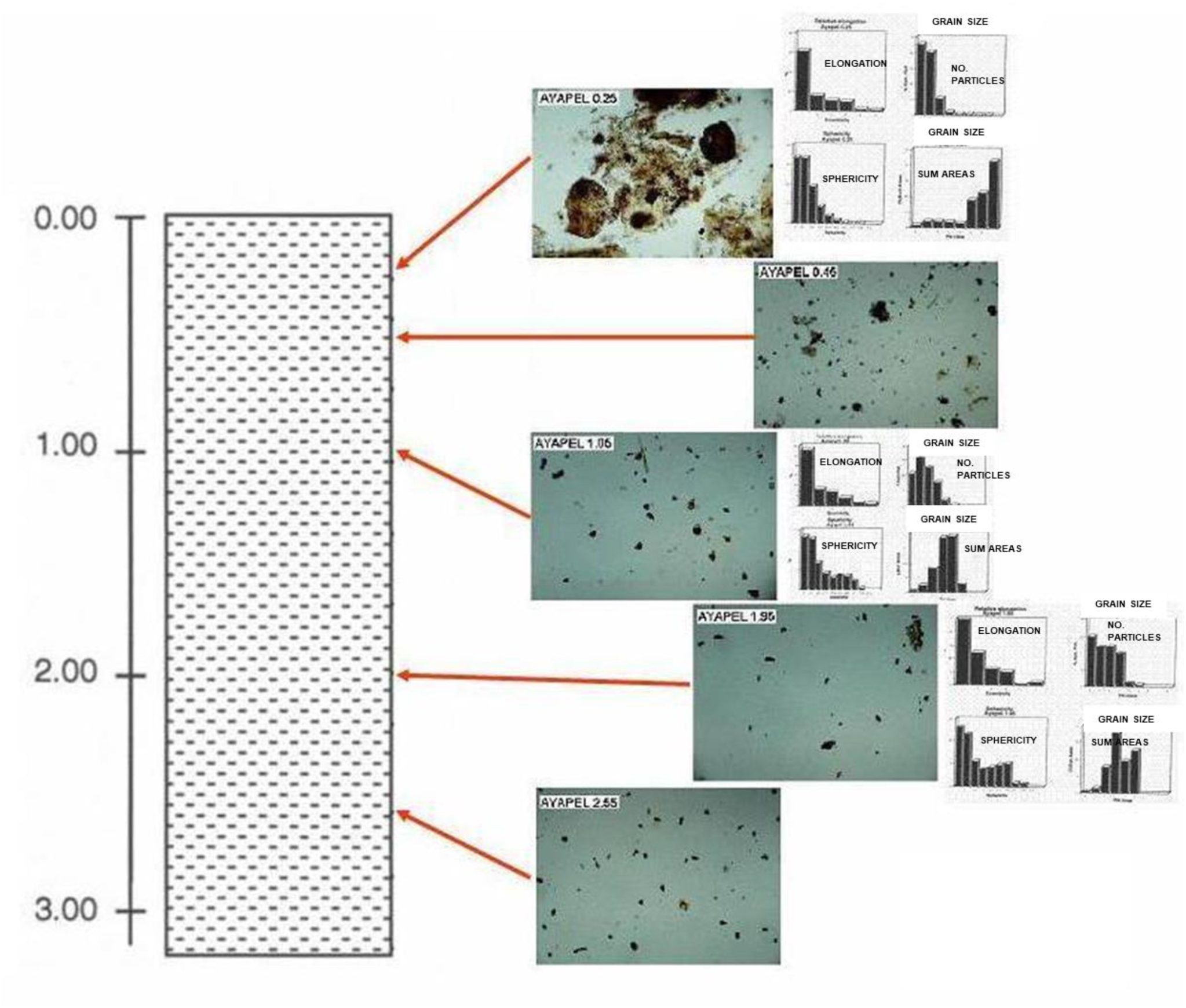
Ayapel section POM quantitative morphological characterization.

#### “Cienaga El Medio”

##### General information

The “Cienaga de El Medio’ is located to the north of Caño Violo, between “Las BoquiIlas’ and “Candelaria’ (borehole coordinates: 74°30’W - 9°08’N, (Figure 1), in the swampy depression, strongly influenced by the Magdalena River flooding. Water stays for long periods covering the area with an average depth of 3 m during the high stand season.

The sedimentation rate for the Boquillas area is according to H.I.M.A.T. (1977) on average 2.92 mm/year, with a maximum value of 4.0 mm/year for the period 1941 - 1975.

##### Lithology and sample description

A core of about 3 m in length represents the sediments of the Cienaga de El Medio swamp. The lithology is heterogeneous (Figure 4a). A total of 11 samples were taken and studied.

**Figure 4a:**
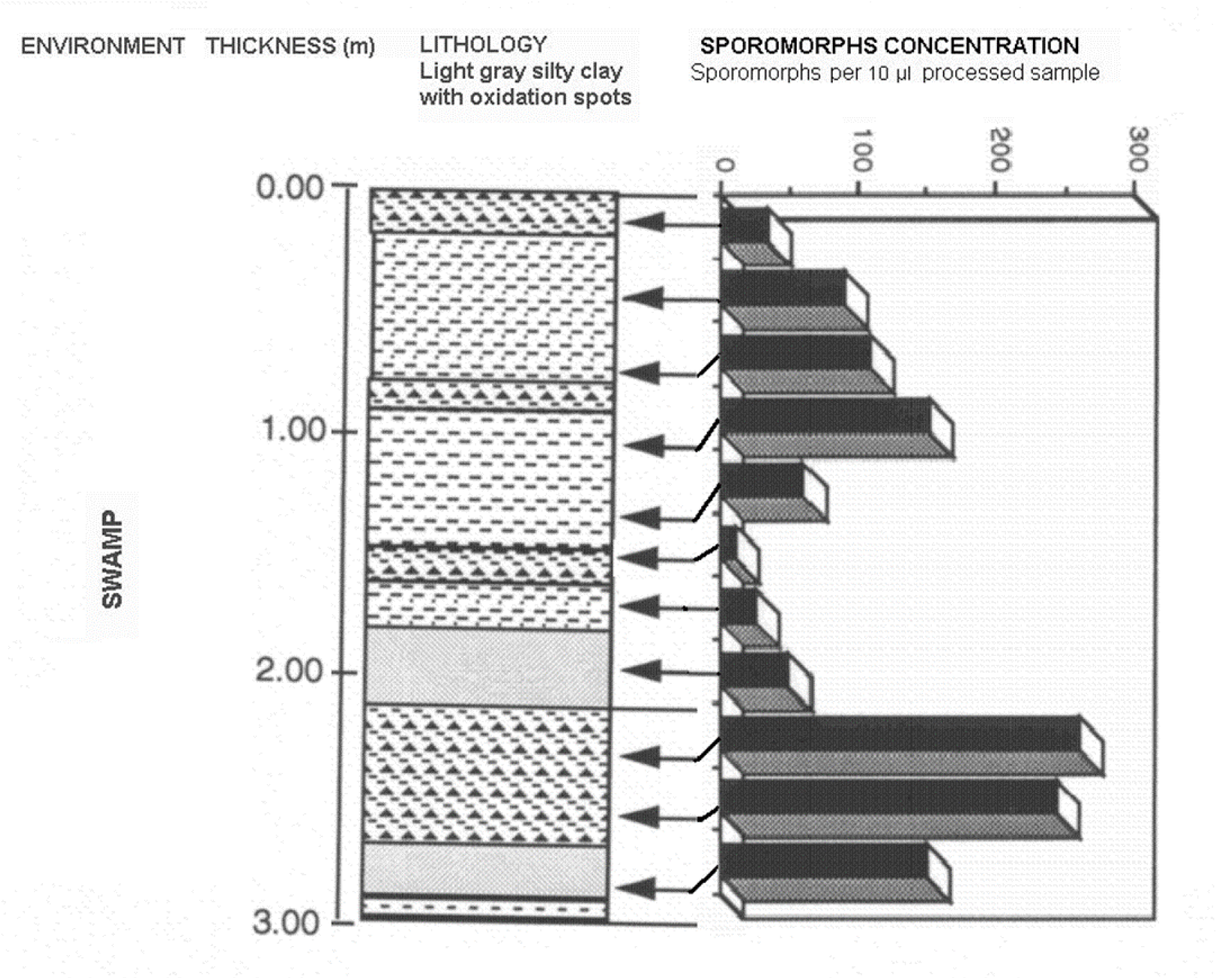
Cienaga El Medio Core Lithology, sample position, and sporomorph concentration. Concentration: Number of Sporomorphs per 10 μl of the palynologically processed sample.

Based on the sedimentation rate (media) estimated as 2,92 mm/year by the H.I.M.A.T. Project, the sampled section of this core represents the last 976 years of sedimentation in Cienaga de El Medio.

##### Palynology

The pollen and spore assemblages are, in general, poor in species and specimens. The richest interval is found between 2.85 m and 2.25 m depth and is dominated by grass pollen and composites (Figure 4b).

**Figure 4b:**
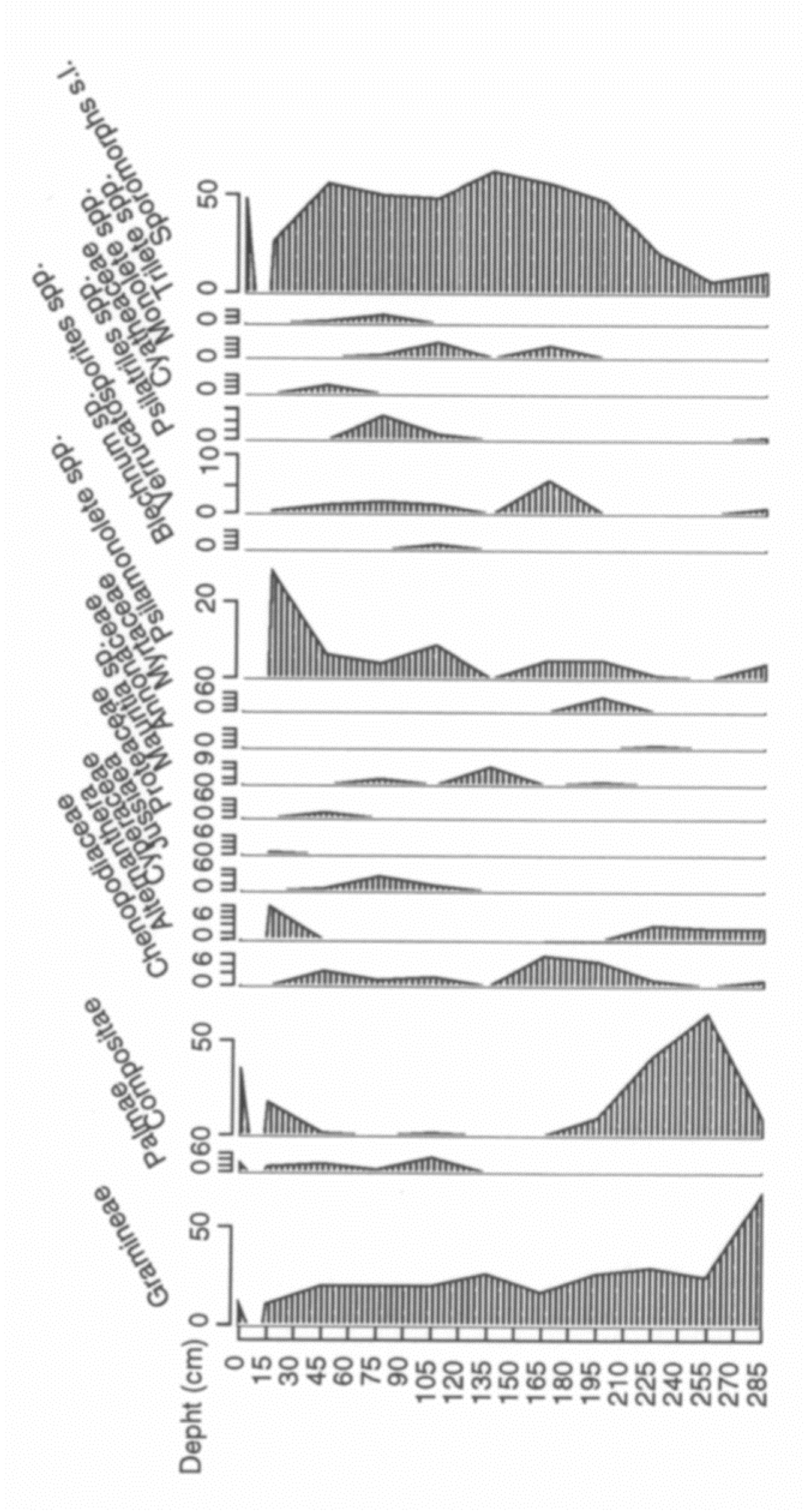
Cienaga de El Medio core sporomorph’s diagram.

Upwards, from 1.95 m to 0.45 m depth, the amount of composite pollen specimens strongly decreases. In contrast, Gramineae pollen together with fem spores are the main components of the assemblage. Chenopodiaceae, Cyperaceae, and *Mauritia* sp. show relative increases.

From 0.15 m depth to the surface, pollen grains of composites become dominant. Fungal remains are as abundant as the sporomorphs, although relative abundance varies from one sample to another. The presence of freshwater algae, including *Botryococcus* sp., is constant throughout the interval. Spores of bryophytes, conodonts, and insect remains are restricted only to this section’s upper part.

##### POM facies

Organic matter concentration in Cienaga de EI Medio is quite low, always below 0.1%, except for at sample 0.15 m depth that reaches 0.14 %.

Regarding POM composition, the observed changes followed the levels of relative oxidation in the sedimentation environment closely, as reflected on the changes of color and abundance of oxidation spots in the Iithology.

The surface sample has an abundance of light-colored plant tissues and some fungal and opaque POM materials.

From 0.75 m to 0.15 m depth, the organic matter is entirely dominated by amorphous finely dispersed POM and in a lower proportion by amorphous opaque POM and ‘organo·mineral gels’. The main difference between these set of samples and the set between 1.95 m and 1.05 m depth is the higher concentration of finely dispersed amorphous POM

From 1.95 m to 1.05 m depth, the level of oxidation of the material increases, based on the rise in the amount of amorphous opaque POM and “organo-mineral gels” POM. In most samples, these types and the finely dispersed amorphous POM account for more than 85% of all the POM material preserved. From 2.85 m to 2.25 m depth, the material is mainly composed of very fine debris classified as amorphous finely dispersed POM. The rest of the material is mainly composed of “organo-mineral” gels, amorphous opaque POM, woody POM, other plant remains, and palynomorphs. However, the most abundant type is the finely dispersed amorphous POM that has to be removed by sieving to describe the other POM types present better.

The morphology of the POM is very similar to the one described for the Ayapel core. The particles’ shape is variable with grain size (particle number), sphericity, and elongation histograms always skewed to the left (Figure 4 c).

**Figure 4c:**
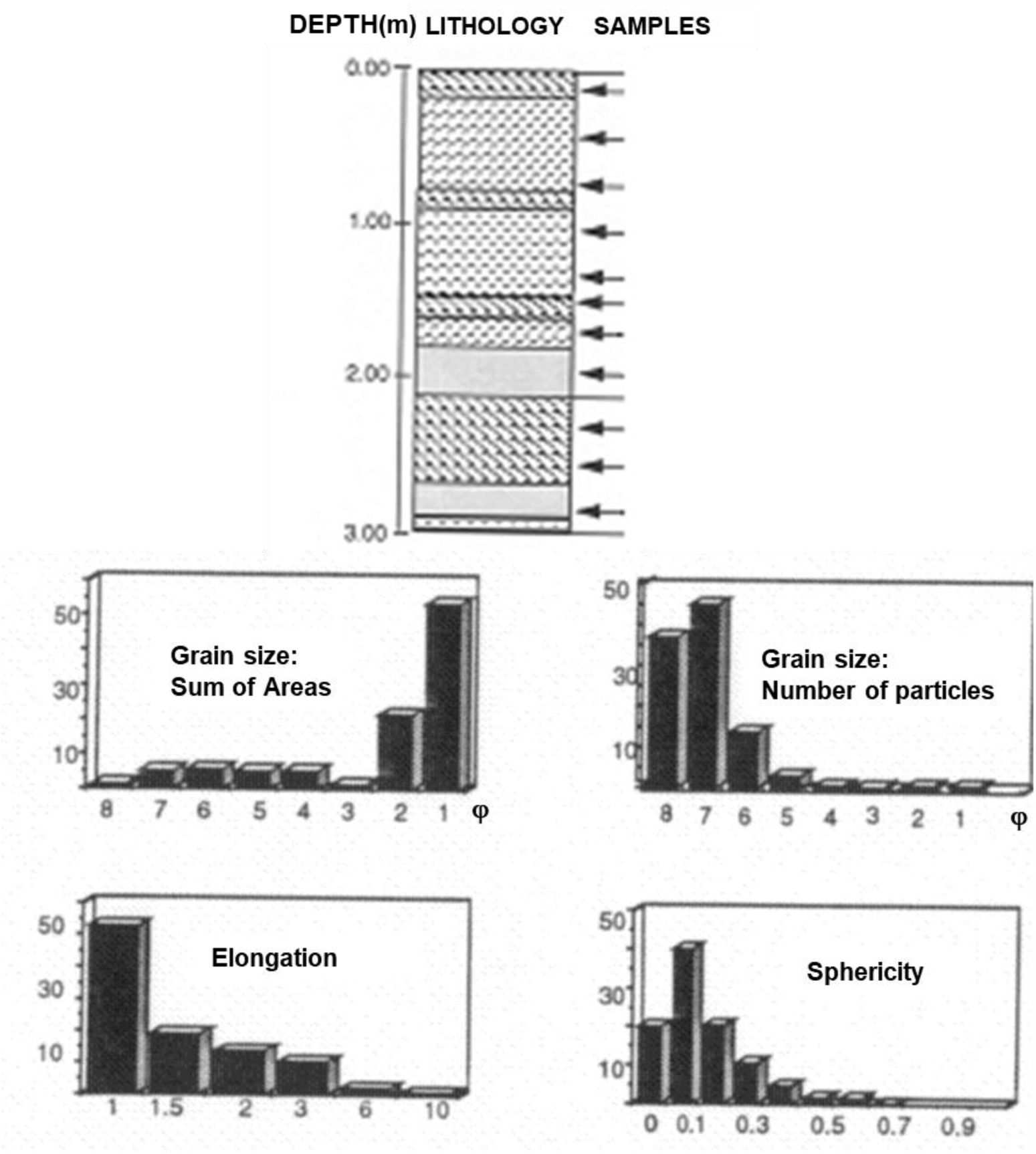
Cienaga El Medio core section POM quantitative morphological characterization.

In the section, three different intervals are recognized based on the characteristics of the grain size distributions:

I. - 2.85 m to 2.25 m depth
II. - 1.95 m to 0.15 m depth and
III. - Surface
I. - The 2.85 m to 2.25 m depth interval is characterized by small particles (clay size range, Φ 8) that, although numerically are close to 50%, cover less than 5% of all the particles area. Between 50% and 65% of particles are equidimensional, with less than 5% in the strongly elongate to filiform shapes
II.- The 1.95 m to 0.15 m depth interval is generally characterized by the numerical dominance of 42% to 50% of a very fine silt size range, Φ 7 particles. Class Φ 7 covers not less than 15% of the particle’s area and commonly more than 20% of it. Showing a relative increase in particle sizes compared with the previous interval, and this is possibly due to the agglutinating effect of the “organo-mineral gel” particles. Most of the samples have 52% to 60% of the particles in the equidimensional range; the relative elongation histogram shows a step-like distribution.
III.- The surface sample is numerically dominated by particles in the silt size range (Φ 7 class). Particles comprised between Φ 8 and Φ 3 classes, clay to very fine sand; cover less than 50% of the particles area with an average of less than 7% per category, whereas particles with sizes larger than Φ 2 class cover more than 50% of the particles area. The relative elongation has a step-like distribution similar to interval 1.95 m to 0.15 m depth.

#### Other Swamps

##### General information

Five samples were taken from the bottom surface of five different swamps from the Lower Magdalena Basin. The sampled swamps were: Cienaga El Medio, Cienaga Pimiento, Cienaga Coyongal, Los Limones and Punta de Blanco. All situated between 8° 40’N - 9° 10’ N and 74°95’ W - 74°25’W (Figure 1).

##### Lithological description

The surface samples lithology is:

CM (Cienaga de El Medio): Silty clay, medium gray color with oxidation spots and roots.

CP (Cienaga Pimiento): Silty clay, medium gray color with oxidation spots. CY (Cienaga Coyongal): Silt with clay, light brown color with plant remains. Los Limones: Silty clay, medium gray color with oxidation spots and root remains

PB (Punta de Blanco): Clayey silt, light brown color with root remains.

##### Palynology

Sporomorph concentration varies from less than 50 grains to more than 250 grains per 10 μl of the palynological processed sample.

In general, the sporomorph assemblage is dominated by grass and composite pollen grains and psilate monolete spores. Also present are *Althernatera* sp., Cyperaceae, *Croton* sp., and *Ludvigia* sp. (Jussiaea) pollen grains.

Fungal remains are the most abundant type of palynomorph. In Los Limones and Punta Blanco, fungal remains account for 2.7% and 5%, while sporomorphs do not exceed 0.3% of the total organic residue after excluding the finely disperse amorphous matter.

Always present in the assemblages are insect remains and freshwater algae (also *Botryococcus* sp.). Occasionally conodonts were found in the assemblages.

##### POM facies

Regarding the POM concentration, the swamp surface assemblages are characterized by a comparatively high organic matter concentration, between 0.1% and 0.3%.

The abundance of light-colored plant tissues characterizes the POM composition, but also cuticular materials, fungal remains, opaque amorphous POM, and animal remains, together with very abundant amorphous finely dispersed POM

Particles show in the grain size histogram (number of particles) that the clay size range materials account for less than 47%. According to the grain size sum of areas histogram, that type of material only represents 2% to 3% of the total particle area. Materials over Φ 2 class, from fine sand upwards, cover more than 60% of the total particle area.

The relative elongation histogram shows that 45% to 60% of the particles fall in the equidimensional category, but usually, not less than 20% have elongate forms.

### Alluvial Flood Plain

#### El Limon

##### General information

El Limon core was taken in an alluvial flood plain, close to the EI Limon town (74°39’N 9°17’W, Figure 1) near the intersection between Brazo de Loba and Brazo de Mompos.

##### Lithology and sample description

The lithology of the 7 m core is heterogeneous (Figure 5 a). A total of 10 samples were taken and studied.

**Figure 5a:**
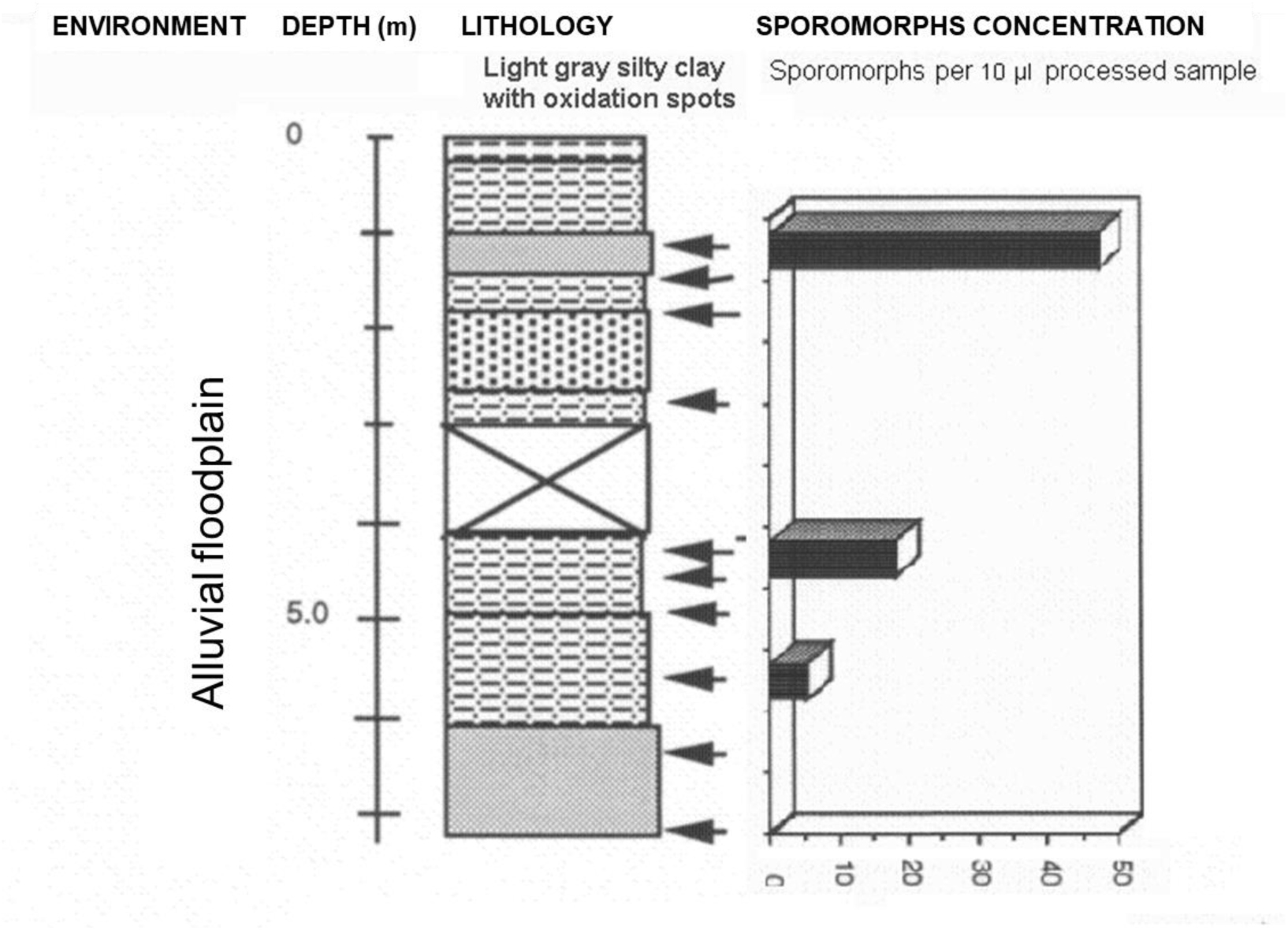
El Limon Core Lithology, sample position, and sporomorph concentration. Concentration: Number of Sporomorphs per 10 μl of the palynologically processed sample.

Core material from 0 m to 1.04 m depth was not available. For that reason, the sampling started at 1.04 m below the surface.

No specific information was available from the H.I.M.A.T. Project about the sedimentation rate of this area. Still, taking a 2.00 mm/yr general sedimentation rate for the Mompos depression, this core may represent about 1400 years of sedimentation on the alluvial plain.

##### Palynology

Regarding sporomorph content, 70% of the samples are very poor to barren (Figure 5b). Only tree samples (1.05 m, 4.37 m, and 5.55 m) had sporomorph assemblages, with specimen concentration from 5 to less than 50 grains per 10 μl of the palynological processed sample.

**Figure 5b:**
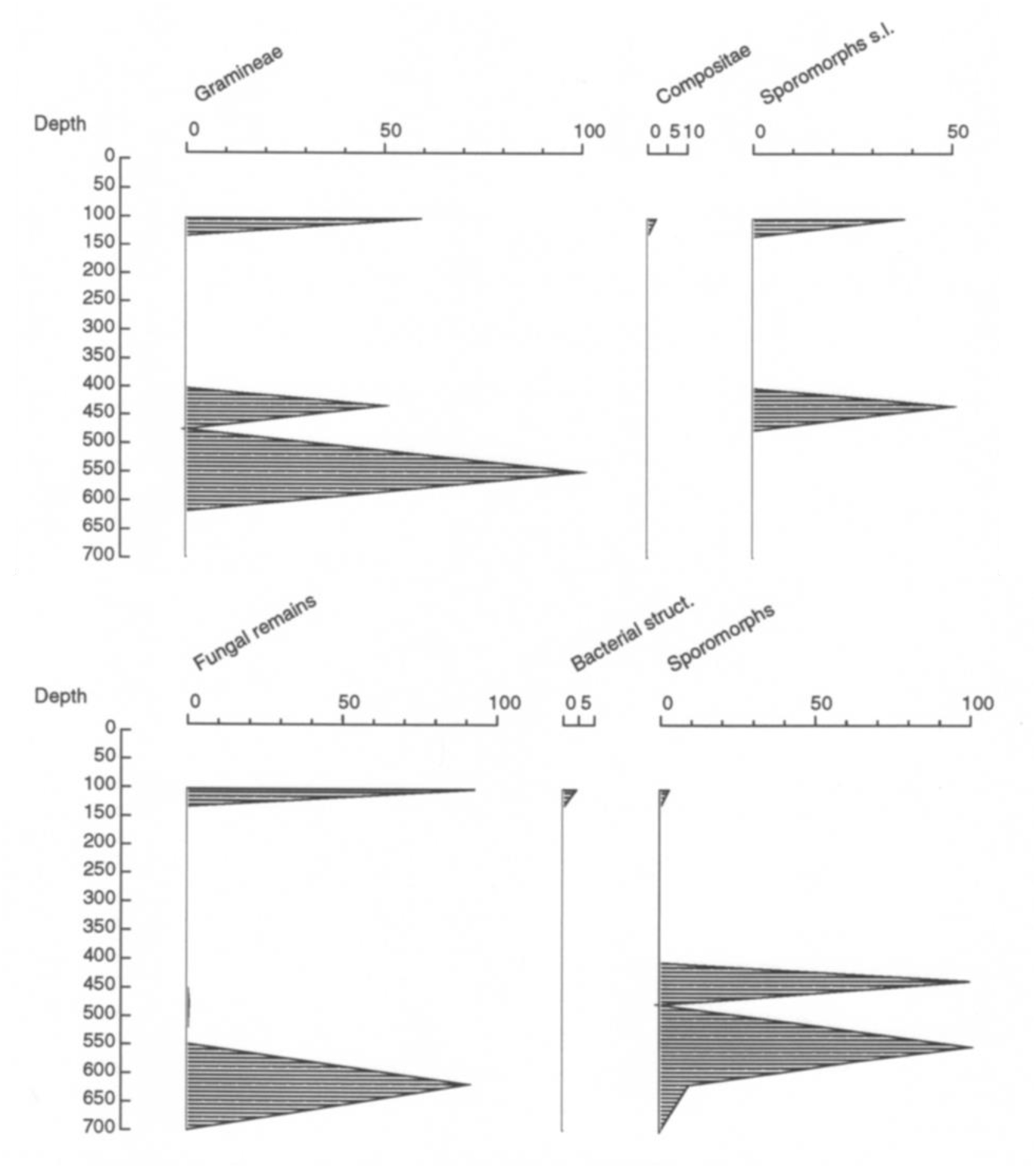
El Limon core palynomorphs diagrams.

**Figure 5c:**
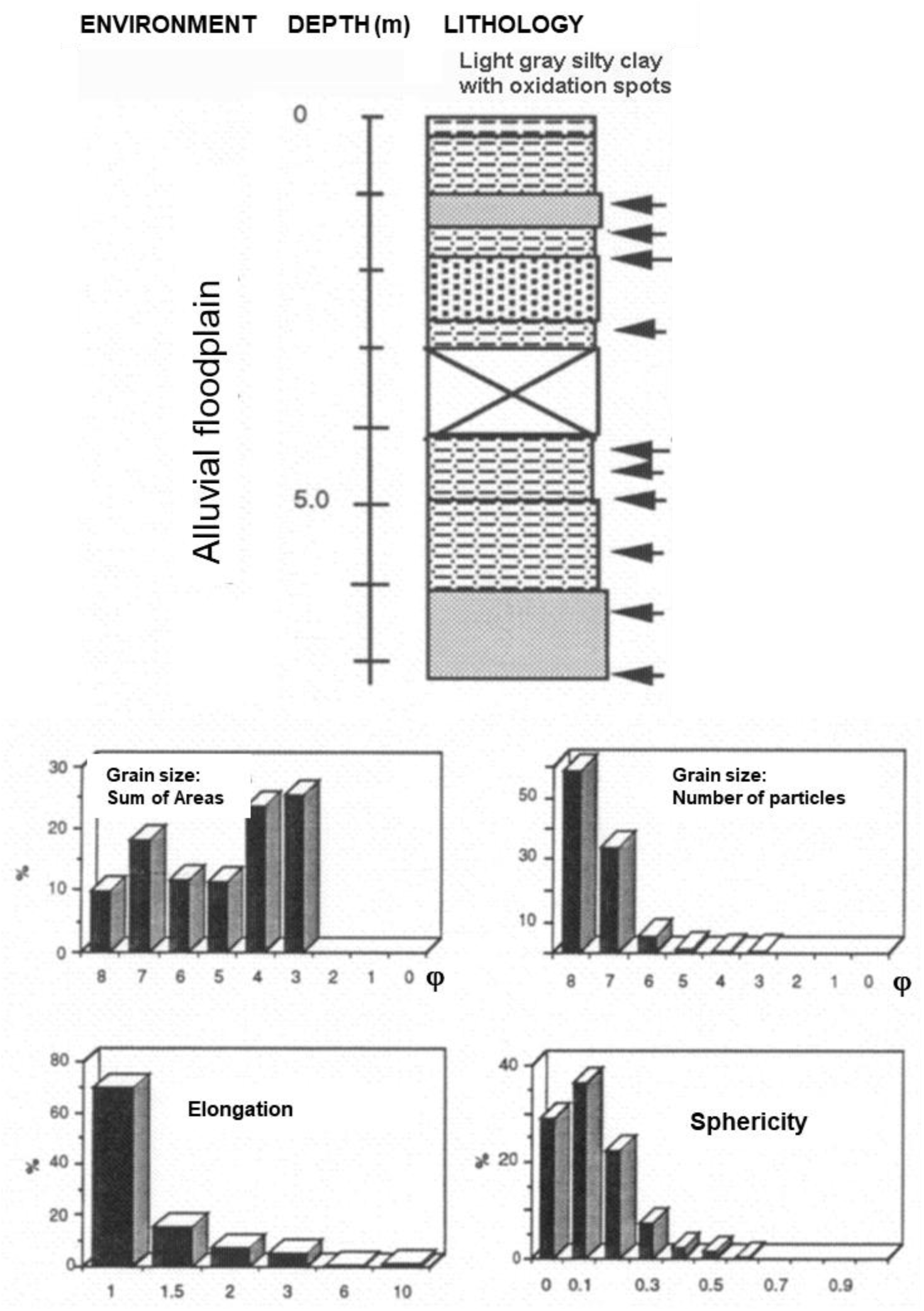
El Limon core section POM quantitative morphological characterization.

The assemblages had grass pollen and only very occasionally composite pollen or poorly preserved unidentifiable grains.

Fungal remains are present in those few samples with sporomorphs.

##### POM facies

Palynological organic matter (POM) recovery from this section was extremely low, approaching 0 % concentration. Nevertheless, some organic matter was recovered.

The POM composition recovered from the 1.37 m to 5.55 m depth interval is of ‘organo-mineral gel” type with comparatively minor amounts of amorphous opaque POM, occasionally present are fungal remains, woody, and other plant tissues.

Samples 1.05 m and 6.25 m to 7.05 m depth contain some amorphous finely dispersed POM.

Finely dispersed organic material tends to hide the rest of the organic components. When removed by sieving, it is possible to determine that all the samples’ organic composition is very similar in the entire section.

Regarding the POM morphological characterization (Figure 6c), grain size distributions range mainly from classes Φ 8 to Φ 3, the grain size distribution histogram, based on the sum of areas, have the first pick at Φ 7 class and a second pick between Φ 4 - Φ 2 classes, usually much more prominent than the first one.

**Figure 6.**
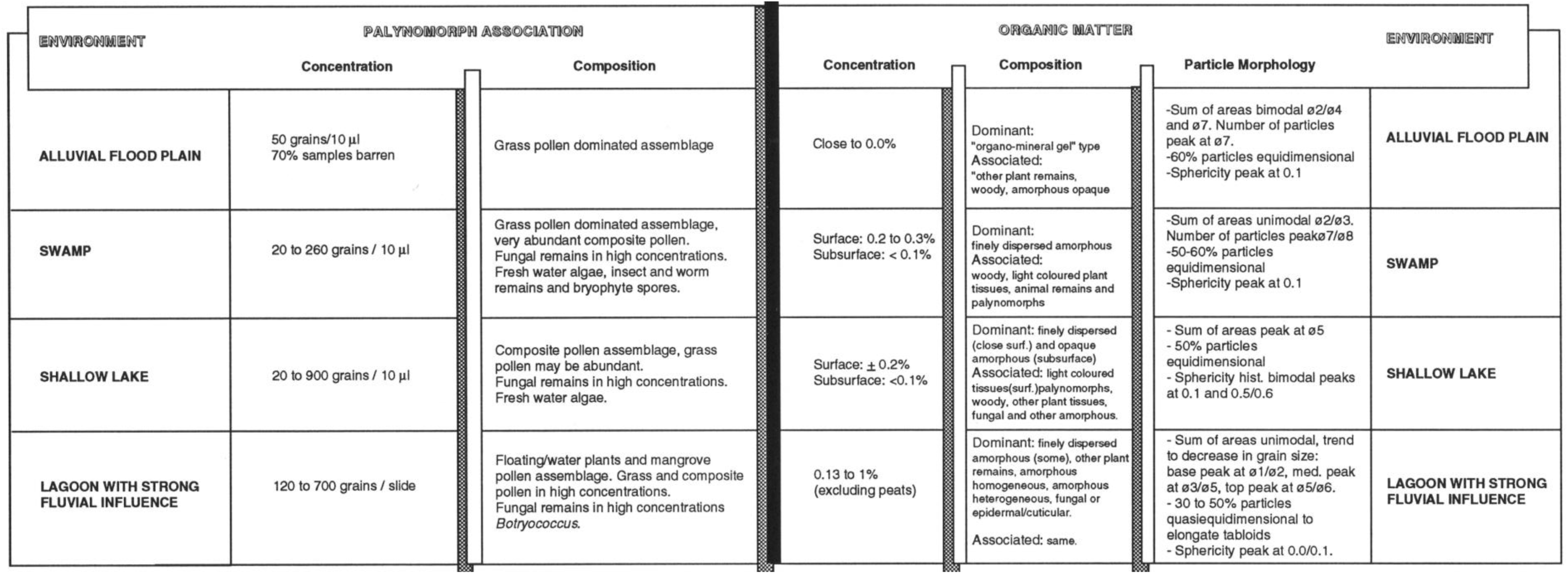
Summary of environments characteristics

In the grain size distribution histogram based on the number of particles, assemblages are dominated by particles in the range of Φ 8 and Φ 7 classes, belonging to the clay and fine silt size range.

More than 60% of all particles fall in the equidimensional range.

## DISCUSSION OF RESULTS

### Comparison of the recent environments

A comparison of the recent environments palynofacies studied is shown in Figure 6. In the next paragraphs, the major differences are discussed.

#### The sporomorphs assemblage

The shallow lake assemblage is dominated by pollen grains of composite followed by grass pollen; water plants’ pollen is present in comparatively lower amounts. The assemblage is, in general, very similar to those recovered from the swamp environment, but in the swamp environment, in general, the grass pollen is dominant over the composites.

The flood basin organic assemblage is usually barren of palynomorphs, and when present, they are pollen grains of grasses and fungal remains.

#### The particulate palynological organic matter (POM)

The organic matter concentration for surface swamp samples varies from 0.1% to 0.3%, while for buried shallow lakes and swamps sediments absolute value is below 0.1%. The flood basin has very low organic content, close to 0.0 % or in the order of 1% of the amount contained in any other environment.

About the type of preserved organic matter, the shallow lake has mainly very dark (opaque unstructured o.m. materials). In contrast, the swamp environment has a clear dominance of finely dispersed unstructured o.m. and “organo-mineral gels”. The scarce material preserved in the flood basin is either the “organo-mineral gel” type or opaque unstructured o.m. or finely dispersed o.m. The shape of particles also differs. In the shallow lake, grain size distribution always has a maximum in class Φ 5 and a bimodal distribution in the sphericity histogram (max. at 0.1 and 0.5/0.6). Those characteristics are systematically absent in the histograms from the swamps.

In the swamp environment, two types of organic assemblages can be differentiated depending on the amount of oxidation: low oxidation and high oxidation. Still, in general, both show a unimodal coarse skewed sum of areas histogram.

The alluvial flood basin has a sum of areas grain size distribution bimodal with the first pick at Φ 7 and a second pick in the coarse silt to sand grain size range (Φ 4 to Φ 2) usually much more prominent than the first one. The grain size distribution histogram is dominated by particles in the range of Φ 8 (clay size range).

### Comparison of recent environments quantitative palynofacies with fossil (Neogene) palynofacies

Available for comparison with a similar set of data for POM textural characterization from the Neogene is the “Marsh-swamp complex” described in Lorente (1986) from two different well cores (VLC 737 and LSJ 3310) from the Miocene Lagunillas Formation of the Maracaibo Lake Basin.

The Neogene swamp environment descriptions fit well with the Cienaga de El Medio cores’ and surface samples findings.

#### Mash - Swamp Palynofacies Comparison

The palynofacies assemblage in the Marsh-swamp complex is highly variable in palynomorphs and organic matter concentration (Figure 7). In the Neogene from the Lake Maracaibo Basin, the palynomorph assemblage is highly variable from one sample to another. When palynomorphs present, the grass pollen dominates the assemblage, with palms, monolete verrucate spores, fungal remains, and freshwater *Pediastrum* colonial algae. This association compares well with the one found in the recent swamp environment from the Lower Magdalena Basin, grass-dominated assemblages, with abundant fungal remains and freshwater algae. Regarding organic matter composition, Neogene’s swamp POM is mainly degraded plant material, with minor amounts of humic gels, woody, and epidermal remains. The composition of the recent assemblages is similar, although the recent swamp environments are dominated by finely dispersed organic matter. The finely dispersed POM was not reported in the Neogene samples, probably due to some differences in the sample preparation (e.g., sieving vs. not sieving).

**Figure 7:**
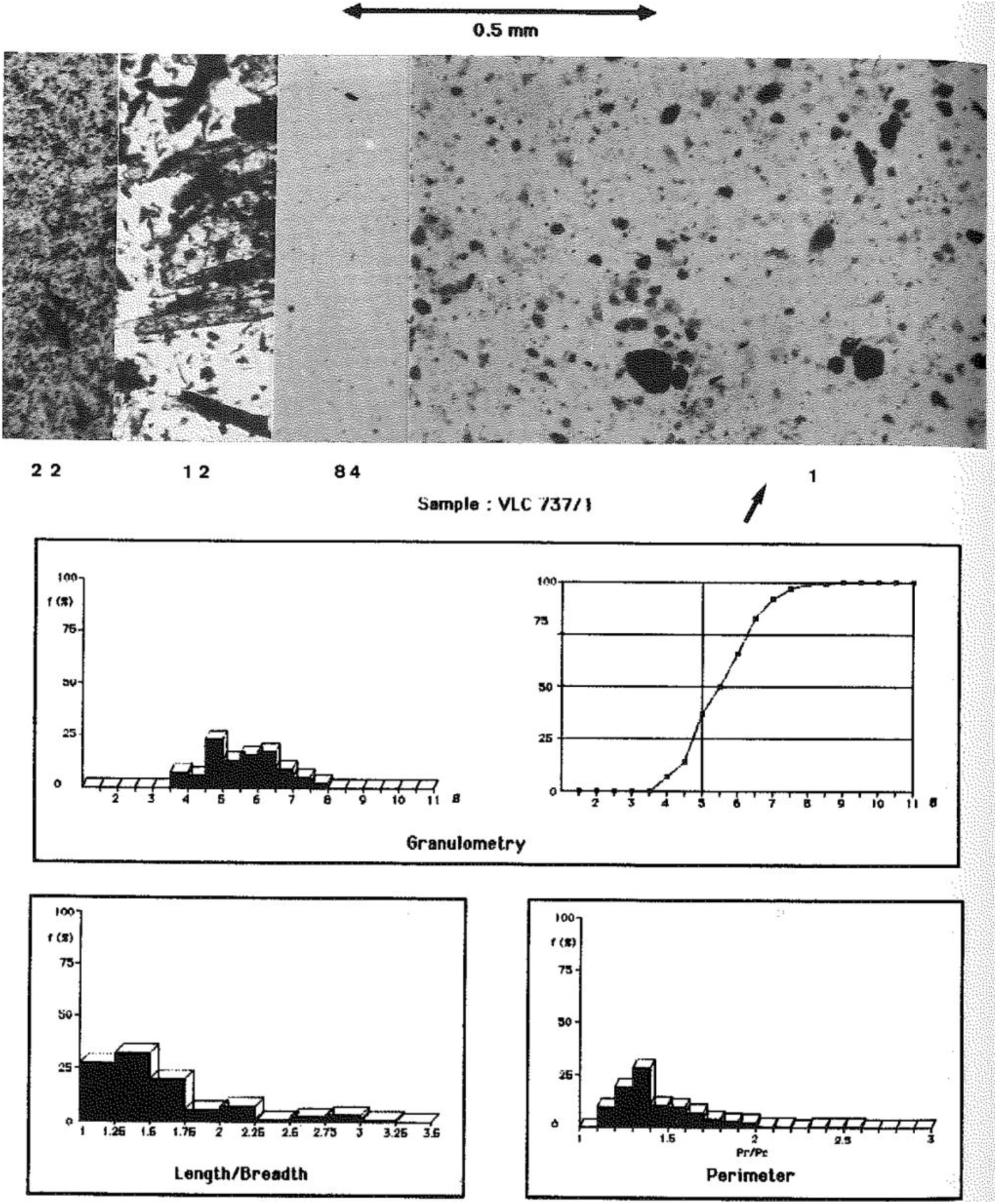
Quantitative palynofacies of Neogene Swamp-marsh complex, Lake Maracaibo, Venezuela (Lorente, 1986).

The POM texture in Neogene sections shows a close relation with the associated lithology. Grain size mainly in the range of Φ 3.5 to Φ 8 class, and most particles equidimensional to quasidimensional in shape (Figure 7). Similar things happen in recent environments, with grain size mainly in the range Φ 2 to Φ 8 class and 50% to 60% of equidimensional shape particles (Figure 4c).

##### The last 1000 yr. climate in the area

The climate on the basin was influenced by E.N.S.O. (El Niño Southern Oscillation) events. That type of event may alter the climate and hydrologic conditions even for decades, with negatives in rainfall during El Niño low–warm phase (Poveda and Mesa, 1997), thus favoring oxidative periods for the POM. Sediment yield is in general controlled by factors like “local topography, soil properties, climate, vegetation cover, morphology, drainage network characteristics, and land use” (Restrepo et al., 2009). But also other factors influence the sediment yield of the Pacific and Caribbean rivers of Colombia as the “(1) the annual shift of the Inter-Tropical Convergent Zone, (2) climatic anomalies due to the El Niño Southern Oscillation during its positive phase of La Niña, (3) low-pressure systems from the Caribbean Sea, (4) orographically driven rainfall within the Andes cordilleras, and (5) cold fronts from the Amazon basin” according to Restrepo et al. 2009.

Although the Magdalena River is a relatively low sediment yield river (Restrepo et al. 2009) with almost no gradient in some parts of the Low Magdalena longitudinal profile, those low gradient areas are associated with the large flood plains like the ones in the Mompos area that comprises 80% of the so-called Cienagas, a kind of natural lakes lying on Cretaceous-Tertiary bedrocks, that act as traps for about 14% of the Magdalena sediment load, being flooded most of the year (van der Hammen, 1986, Restrepo, 2008), with a general sedimentation rate of 2.0 mm/yr (Restrepo, op. cita).

Van der Hammen created frequency curves based on C14 dated peat and dark humic layers, and Cleef (1992) based on the works by Plazas et al., 1988 and van der Hammen (1986), those curves showed three dry periods for the last 1500 years: (I) 1300-1500 years bp, (II) 750-650 years bp., and (III) 400-500 years bp. Some of those are recognized on the studied cores.

##### Freshwater shallow Lake: Ayapel

The generally homogeneous lithology points out to quite stable conditions in the environment during the recovered core sedimentation. Minor exceptions are the five intervals with roots and root casts (0.25 m; 1.65 m) or plant remains (1.05 m; 2.25 m; 3.20 m) that point to “minor” variations within the same environment. The root casts and remains may indicate shallower conditions and or withdrawal of the lakeshore due to dryer periods that allowed land plants to advance over the shallow lakeshore, while the intervals rich in plant remains may be indicative of abundant rain periods favoring the transport of plant material by torrential flows from the adjacent shore areas to the lake.

Based on the average sedimentation rate for the area estimated as 2.7 mm/year (H.I.M.A.T., 1977), ages for the samples were calculated (Table 1).

**Table 1:**
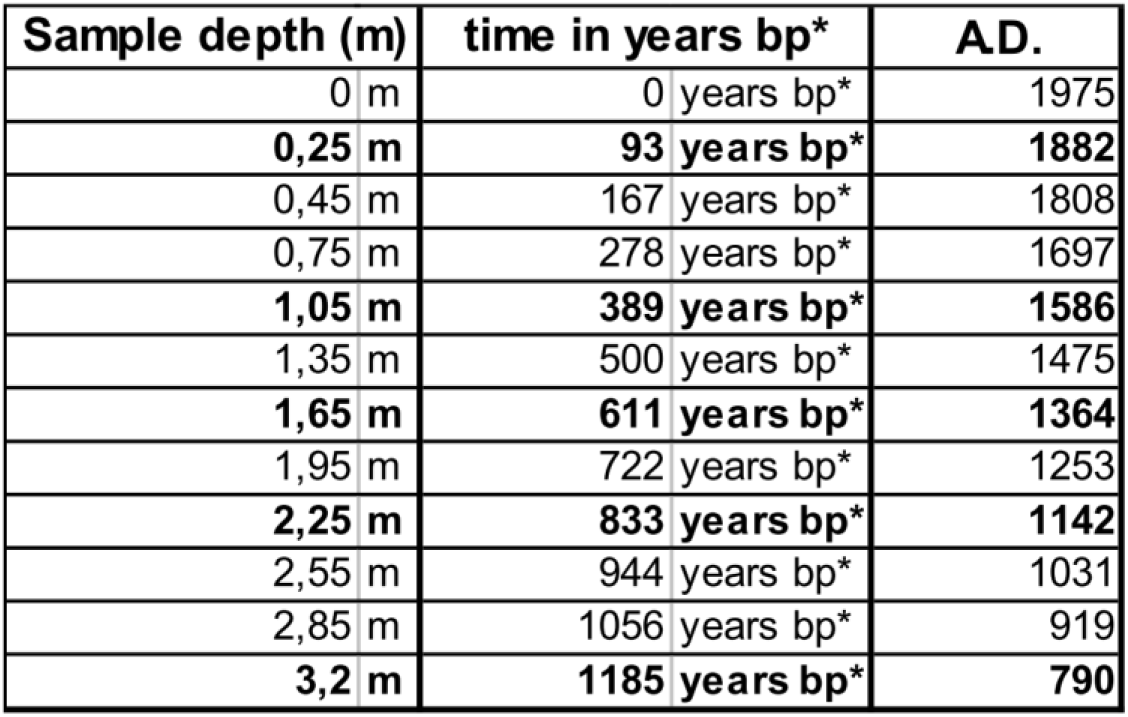
Calculated ages for the Ayapel Lake Core samples in this study. * = years before present. 1975 is the year established in this work as present because it was the approximate year of sampling. A.D. = Anno Domini, referring to years after the birth of Jesus Christ.

| **Sample depth (m)** | **time in years bp\*** | **AD.** |
| --- | --- | --- |
| 0 m | 0 years bp* | 1975 |
| **0,25 m** | **93 years bp\*** | **1882** |
| 0,45 m | 167 years bp* | 1808 |
| 0,75 m | 278 years bp* | 1697 |
| **1,05 m** | **389 years bp\*** | **1586** |
| 1,35 m | 500 years bp* | 1475 |
| **1,65 m** | **611 years bp\*** | **1364** |
| 1,95 m | 722 years bp* | 1253 |
| **2,25 m** | **833 years bp\*** | **1142** |
| 2,55 m | 944 years bp* | 1031 |
| 2,85 m | 1056 years bp* | 919 |
| **3,2 m** | **1185 years bp\*** | **790** |

From the perspective of the type of POM facies preserved in this environment, it varies dramatically with burial history (Figure 8).

**Figure 8:**
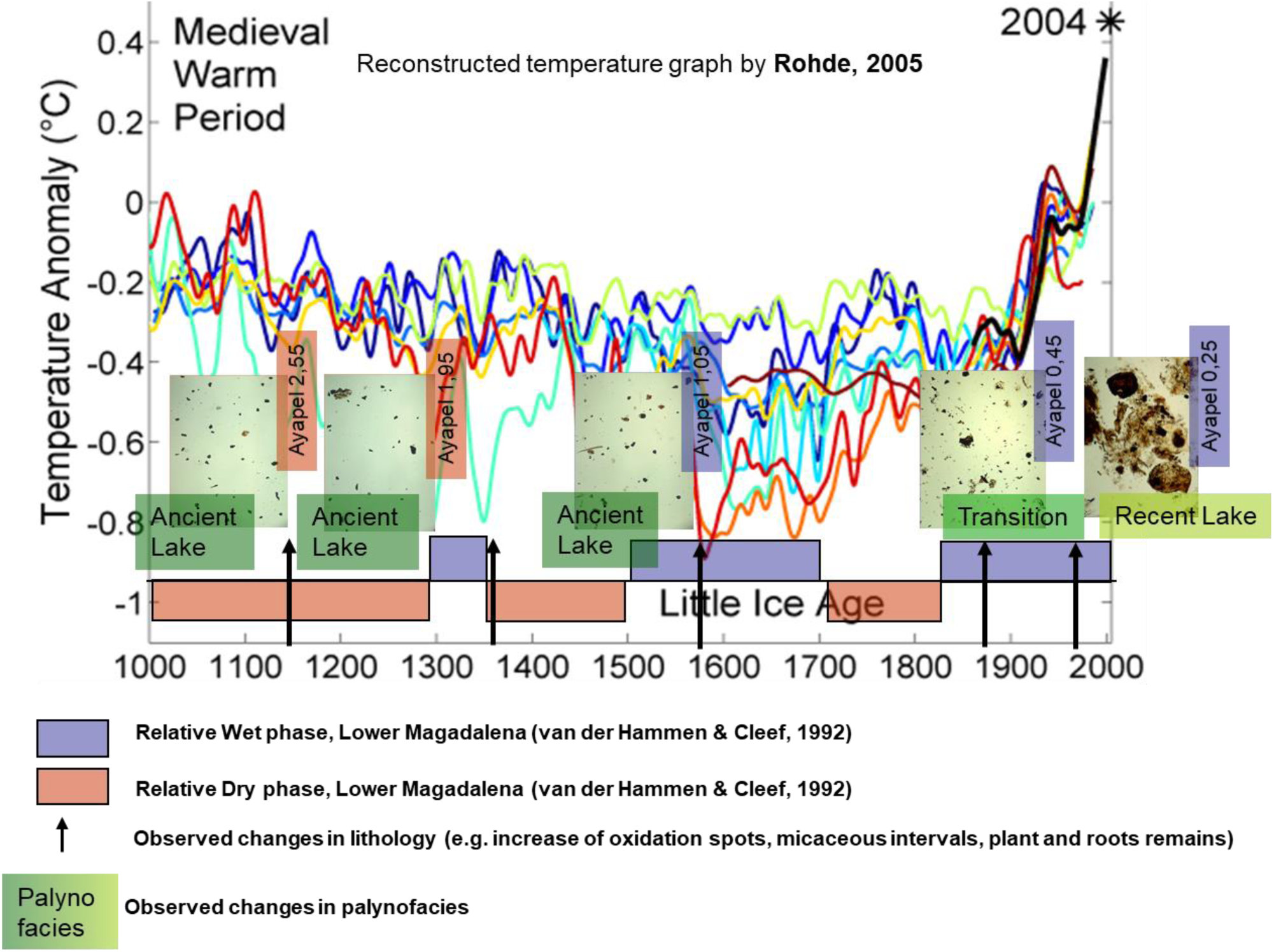
Ayapel Lake Core palynofacies related to the temperature variation (Rohde, 2005) and the wet-dry phases (van der Hammen & Cleef, 1992) during the last 1000 yr.

The three key intervals identified are:

I – The “water-sediment interface” or “present lake” (0 to 0.25 m depth). This zone coincides with a wet and warm phase (Figure 8).
II - The ‘transition zone’ (0.25 m – 0.50 m depth). This zone coincides with a wet but colder phase (Figure 8).
III - The “ancient lake” (below 0.50 m depth). This zone contains the wet and cold phase known as the “Little Ice Age” but also encompasses other dry and warmer phases, as shown in Figure 8.

Rainy vs. dry seasons POM facies assemblages were described previously in short-term cycles from the Upper Orinoco Delta’s recent environments (Lorente, 1990). Those short-term cycle assemblages are characterized by the alternation of coarse, light-colored, cuticles and woody frequently elongate, poorly sorted p.o.m, with assemblages of finer, denser, frequently equidimensional, well-sorted, unstructured opaque POM.

On the other hand, van der Hammen and Cleef (1992) described zones of marsh vegetation develop along the shores of the back swamp lakes, with “floating meadows” of grasses that root at the shore (Figure 9), those floating meadows according to those authors gave origin to peat layers intercalated on the clayey, silty and sandy deposits formed in those shores during the low water dry periods. Those authors correlated the peaty layers with periods of lower rainfall. But no peat layers were identified on the sampled core.

**Figure 9:**
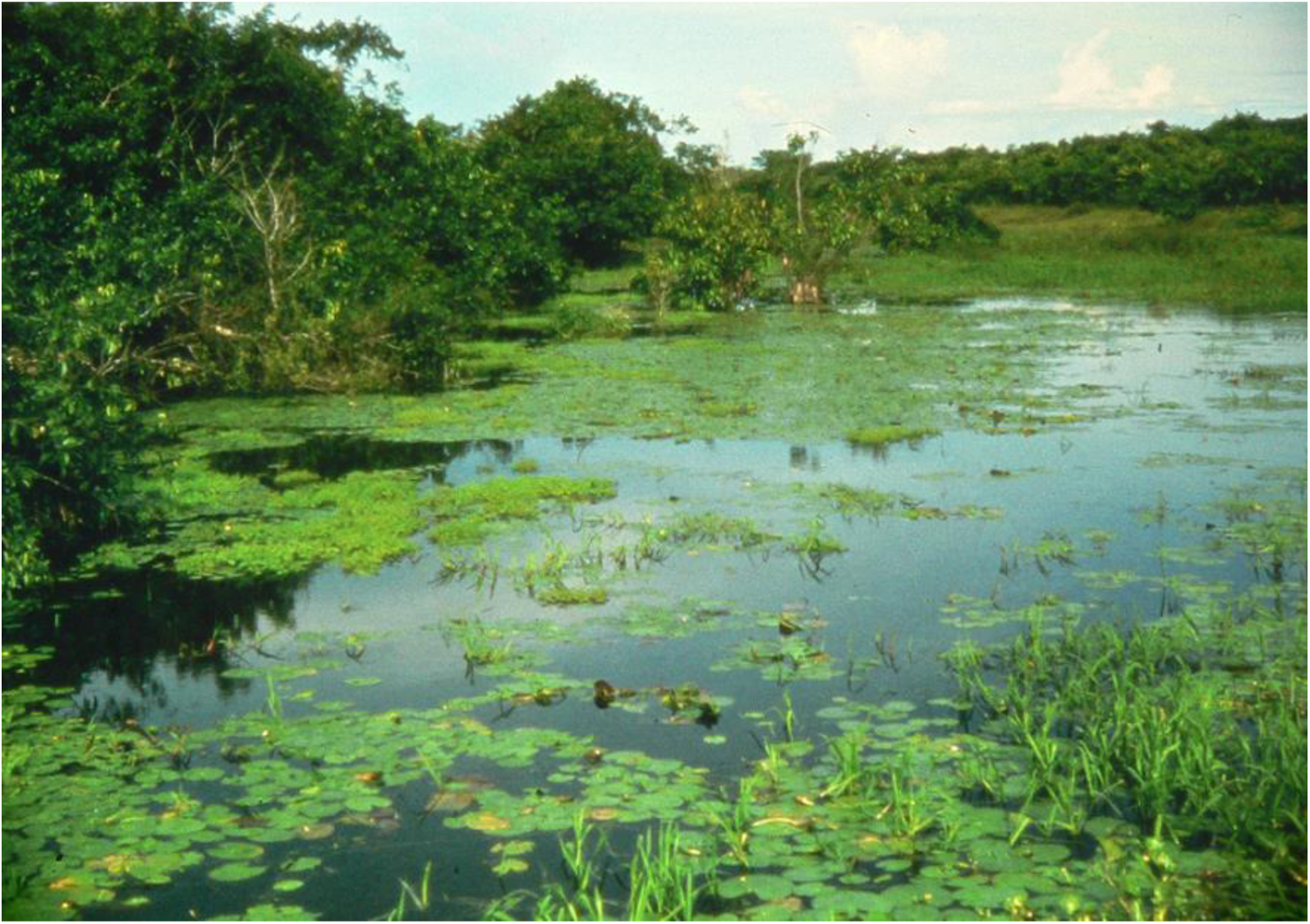
Ayapel lake “floating meadows” and shore vegetation. The authorship of the photo is unknown.

Integrating lithology and POM facies assemblages information, it is possible to visualize a colder and dryer scenery than present-day conditions during the interval from 1100 A.D. until 1850 A.D. (3.20 m to 0.5 m), however, interrupted by warmer and relatively humid shorter events (Figure 8). This trend changed about two centuries ago when a progressive shift towards a warmer and humid climate was registered in the interval from 0.45 m depth to the surface.

##### Swamps

The swamp environment was characterized through two different sets of samples, the surface samples, and the Cienaga de El Medio core. Surface samples characterize the palynofacies assemblage as is deposited under present-day conditions. The core allowed us to understand the changes in the palynofacies assemblages within the first meters of burial history (Figure 10). Considering an average sedimentation rate of 2.92 mm/year (H.I.M.A.T., 1977), with a maximum value of 4.0 mm/year for the period 1941 – 1975, approximate ages were estimated (Table 2) for the interval.

**Figure 10:**
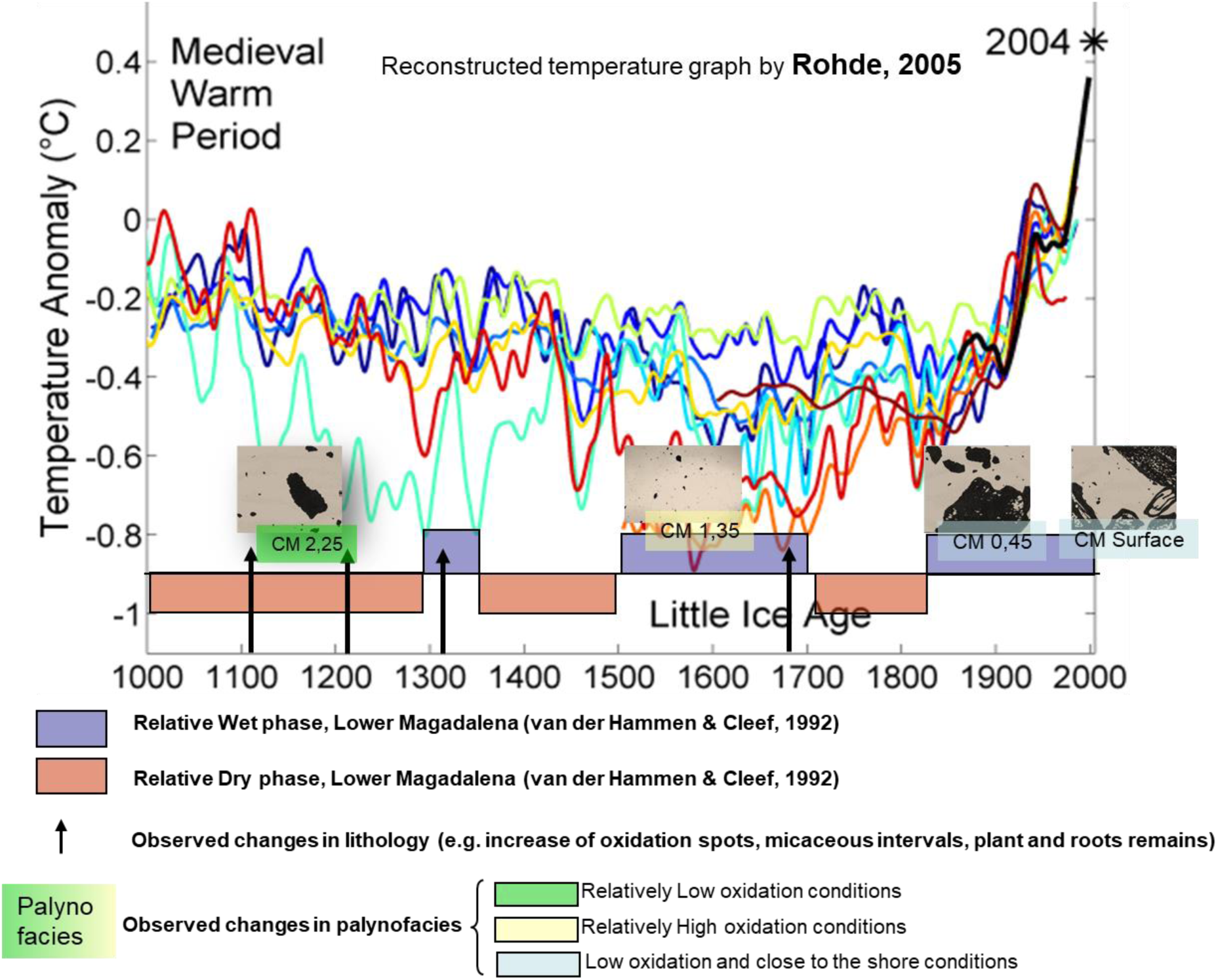
Cienaga El Medio Core palynofacies related to the temperature variation (Rohde, 2005) and the wet-dry phases (van der Hammen & Cleef, 1992) during the last 1000 yr.

**Table 2:** Calculated ages for the Cienaga El Medio Core samples in this study. * years before present, where 1975 is the year established in this work as present because it was the approximate year of sampling. A.D. = Anno Domini, referring to years after the birth of Jesus Christ.

| **Sample depth (m)** | **time in years bp\*** | **A.D.** |
| --- | --- | --- |
| 0 m | 0 years bp* | 1975 |
| 0,13 m | 33 years bp* | 1943 |
| 0,25 m | 74 years bp* | 1901 |
| 0,45 m | 142 years bp* | 1833 |
| 0,75 m | 188 years bp* | 1788 |
| **1,05 m** | **290 years bp\*** | **1685** |
| **1,35 m** | **450 years bp\*** | **1525** |
| 1,65 m | 496 years bp* | 1479 |
| **1,95 m** | **656 years bp\*** | **1319** |
| **2,25 m** | **759 years bp\*** | **1216** |
| **2,55 m** | **861 years bp\*** | **1114** |
| 2,85 m | 964 years bp* | 1011 |

The “present-day sedimentation” or water-sediment interface sporomorph assemblage is similar to the one described for the upper part of Ayapel Lake, so as expected, the present sedimentation conditions registered in both localities are very similar.

Two intervals are recognized based on lithology and POM characteristics:

- High Oxidant Conditions Assemblage (HOCA) (1,95 – 1,05 m).
- Oxidant Conditions Assemblage (NOCA) (2,85 – 2,25 m).

The observed variations in the POM assemblages interpreted as related to the increase/decrease of oxidation due to the sediment input and water levels are assumed to correlate with climate conditions changes.

When plotted on the climate chart of the last 1000 years superimposed with the dry/wet phases of van der Hammen 1986, NOCA Assemblage coincides with a dry phase, while the HOCA Assemblage coincides with wet phases, which includes the “Little Ice Age”.

##### Alluvial flood basin

The alluvial flood basin is a very difficult environment for organic matter preservation due to the high levels of oxidation, favored by short flooded times and longer periods under dry conditions with the sediments interface exposed. Energy is also highly fluctuating in this environment.

No specific information was available from the H.I.M.A.T. Project about the sedimentation rate of this area. I used the equivalent rates of 3.8 mm/yr average determined by Smith (1986) for the Magdalena river basin’s extensive wetlands to calculate the probable age of the only event singled out in the last 1000 y. Characterized by an increase in the pollen count with grasses and fungal remains as only components of the palynomorphs assemblage and some unstructured organic matter, it may have occurred circa 1699 A.D. close to the end of the “Little Ice Age” based on the probable age calculation.

## CONCLUSIONS

- Swamp and shallow water lake surface palynofacies may seem indistinguishable, but:
- Each environment (swamp, shallow water lake, and alluvial flood basin), when analyzed in detail, has its own set of characteristic palynofacies that reflect probably seasonal changes within the environment.
- Most recent palynofacies within one environment do not necessarily conform to older palynofacies within the same environment. This is due to climate, and other local conditions change through time (oxygen content, energy, vegetation, availability of sediments, temperature, rainfall, etc.).
- Palynofacies changes in depth (time) within a single environment are related in this study with climate changes.
- It was possible to correlate the observed changes with the dry/wet phases described by van der Hammen (1986) in the Lower Magdalena – San Jorge River Basin.
- When carrying out this type of studies is necessary to keep in mind that the complexity of the changes observed within the first meters of burial history may have different origins, and with no intention to be exhaustive, examples of those are:
- Decomposition of organic matter due to biological activity (bacterial, microbial, fungal decomposition)
- Decomposition of organic matter due to chemical processes like oxidation or replacement, e.g., generation of inert materials: opaque unstructured and “organo-mineral gel” POM types.
- Variations on the amount of organic matter input may be due to oscillations of the environment’s energy, related to variations of the amount of rain in the catchment area (climate).

## Supporting information

Attachment 1

## ACKNOWLEDGEMENTS

The author dedicates this paper *in memoriam* of two outstanding South America tropics researchers: Prof. Dr. Thomas van der Hammen my mentor, and Dr. T. A. Wijmstra, my guide and inspiration for this research project.

This research was supported by S.I.P.M, The Hague, and the Hugo de Vries Laboratorium of the Universiteit van Amsterdam.

## Notes

### Competing Interest Statement

The authors have declared no competing interest.

