## Supplementary material for "Palynofacies, environments, and climate changes in the Magdalena River Basin": Attachment 1

UNIVERSITEIT VAN AMSTERDAM **Hugo de Vries-Laboratorium** VAKGROEP SYSTEMATIEK, EVOLUTIE EN PALEOBIOLOGIE
AFD. PALYNOLOGIE EN PALEO/ACTUO-OECOLOGIE
Prof. Dr. H. Hooghiemstra

### FIRST RESULTS FROM A QUANTITATIVE PALYNOFACIES STUDY OF RECENT ENVIRONMENTS FROM THE LOWER MAGDALENA RIVER BASIN, COLOMBIA.

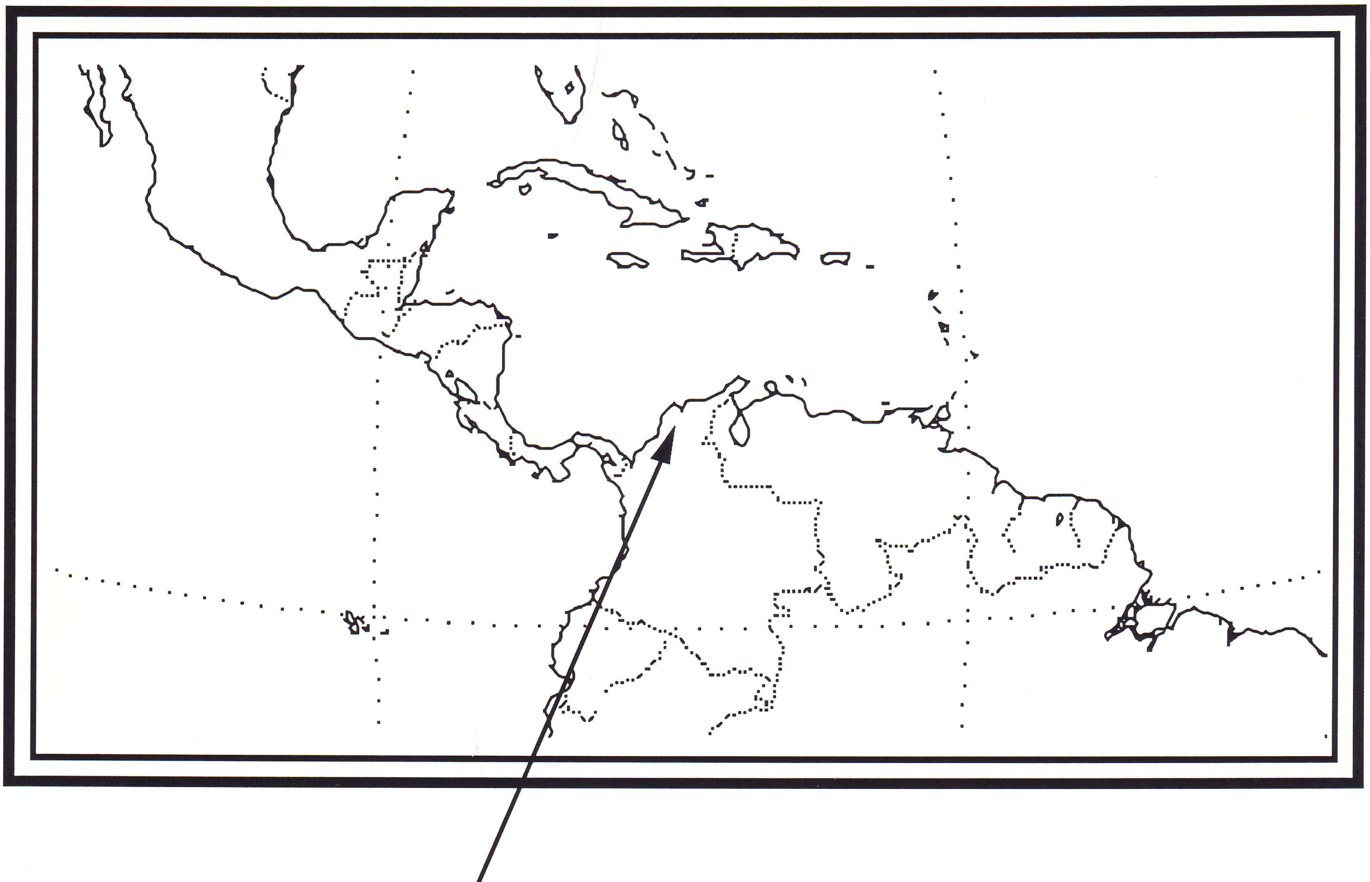

Lower Magdalena River Basin

by: **M. A. LORENTE**

**January, 1992**

**FIRST RESULTS FROM A QUANTITATIVE PALYNOFACIES STUDY OF RECENT ENVIRONMENTS FROM THE LOWER MAGDALENA RIVER BASIN, COLOMBIA.**

M. A. LORENTE

HUGO DE VRIES LABORATORIUM, KRUISLAAN 318 1098 SM AMSTERDAM.

**CONTENT**

- INTRODUCTION 3

Magdalena River General Information 3
Vegetation in the Lower Magdalena River Basin 4

- PALYNOLOGY AND PALYNOFACIES OF RECENT ENVIRONMENTS 6

The Ayapel Section 6
Location and general information
Lithology and sample description
Palynology
Pollen and spores diagram
Palynomorphs diagram
Ecological groups diagram
Palynofacies
Organic matter concentration
Organic matter composition
Organic matter morphological characterization

The Cienaga de El Medio (CM) Section 18
Location and general information
Lithology and sample description
Palynology
Pollen and spores diagram
Palynomorphs diagram
Ecological groups diagram
Palynofacies
Organic matter concentration
Organic matter composition
Organic matter morphological characterization

The El Limon Section 29
Location and general information
Lithology and sample description
Palynology
Pollen and spores diagram
Palynomorphs diagram
Ecological groups diagram
Palynofacies
Organic matter concentration
Organic matter composition
Organic matter morphological characterization

The Santa Marta (BL) Section 39
Location and general information
Lithology and sample description
Palynology
Pollen and spores diagram
Palynomorphs diagram
Ecological groups diagram
Palynofacies
Organic matter concentration
Organic matter composition
Organic matter morphological characterization

The surface samples 50
Location and general information
Lithology and sample description
Palynology
Pollen and spores diagram
Palynomorphs diagram
Ecological groups diagram
Palynofacies
Organic matter concentration
Organic matter composition
Organic matter morphological characterization

- DISCUSSION OF THE ENVIRONMENTS 59

The fresh shallow water lake environment 59
The swamp environment 60
The flood alluvial basin environment 63
The Lagoon with strong fluvial input environment 64

- COMPARISON OF THE ENVIRONMENTS 65

The palynomorph association
The organic matter

- GENERAL REMARKS AND RECOMMENDATIONS 67

- REFERENCES 70

- ANNEX
1.- Methodology for organic matter concentration determination 71
2.- Amsterdam Palynological Organic Matter Classification 72
3.- Measurements taken by OMAS (Magadalena Project) 74
4.- Relative elongation scale 75
5.- Summary of the Environments 76

#### - INTRODUCTION

This report presents the initial results from the study of a first set of 61 sediment samples taken from Recent fluvial to coastal environments, as part of the project "Recent palynofacies in the Lower Magdalena River basin" carried out at the University of Amsterdam with the sponsorship of S.I.P.M.

The study has been focused on the characterization of recent environments throughout the quantitative study of palynofacies. A set of cores from different environments from the Lower Magdalena River basin in Colombia were investigated as well as a limited set of surface samples (fig. 1).

Most of the original cores and surface samples were taken during the development of the multidisciplinary project designed to recover the flooded area of the Lower Magdalena River for agricultural purposes. That project was developed as part of a cooperation program between the Dutch and the Colombian Governments. The original material was kept in very good conditions at the INGEOMINAS installations in Bogota, Colombia, were the author was allowed to sample it.

In this research each step, from the sample preparation to the final search of the palynological residues, was made in a systematic and as quantitative as possible way, to ensure comparable results.

General methods followed here are similar to the ones described in Lorente 1986, 1990a/b, for quantitative studies of palynofacies. A different process was designed to improve the measurement of the amount of organic matter recovered after sample preparation and calculation of organic matter concentration. The process was already described in the first presentation of this project to S.I.P.M (annex 1).

The organic matter classification used follows the guide-lines of the "Amsterdam Palynological Organic Matter Classification" (annex 2) as was approved on June 1991, after the Open Workshop on Organic Matter Classification, organized by the University of Amsterdam as part of this research, and that was partially sponsored by Shell Nederland.

Quantification of morphological characteristics of the organic particles was done by means of the O.M.A.S. system (annex 3), as developed at the Hugo de Vries Laboratory.

##### Magdalena River General Information

The tributary basin of the Magdalena River is located between 2º and 11º N, restricted to equatorial latitude, covering a drainage area of 257.000 km2 (Magdalena - Cauca). The total river length (from Barranquilla to Paramo de las Papas) is 1550 km, with a flow rate of 6700 m3/sec. The precipitation average in the area is 2000 mm (from 5000 mm maximum to 800 mm minimum).

The swampy depression ("depresion cenagosa" or "zona lacustrina") has an area of 207.700 Km2. The statistics from this area show that the "flooded area" cover 16% of the total swampy depression while the "dry area"covers about 84%. The flooded area has the following regime:

6 - 12 months inundated: 19%
3 - 6 months inundated: 30%
1 - 3 months inundated: 14%
less than 1 month inundated: 21%

The average annual sedimentation rate (swampy depression, last 1500 years) is 2.9 mm, but there are significant variations related with local conditions, the main areas are:

Palmitas: 2.7 mm
Sucre: 3.3 mm
Boquillas: 4.0 mm
Mompos: 1.7 mm

**Vegetation in the Lower Magdalena River Basin**

The flora in the lower course of the Magdalena river has been described by Cuatrecasas (1958) and Wijmstra (1967).

The following types of vegetations were described from the river-plain of the Magdalena River: grassy plains with scattered groups of trees or open vegetation, gallery forest along the riversides or dense vegetation and the vegetation belts around the cienagas.

The open vegetation typically occur under rather extreme climatic conditions, e.g. with rain fall concentrate in few months every year and long dry periods or when big changes of the ground-water-table occur (Wijmstra, *op. cita*). It consist of two different types depending on vegetation density:
- The grassy plains: with grasses as *Andropogon bicornis*, *Paspalum millegranum*, *Panicum vulgaris*, etc.; palms and trees as *Mauritia minor*, *Hirtella elongata*, *Bowdichia*, *Byrsonima crassifolia*, *Curatella americana*, *Palicourea rigida* and *Cecropia.*
- The "savanna woodland" type: with all the above mentioned plant association plus *Croton glabellum* and *Ficus elliptica*

The gallery forest along the river sides, usually is characterized by a dominant species of tree as *Inga calliandra*, *Albizzia* sp., *Heliocarpus* sp., *Salix* sp., *Cecropia* sp., or *Celtis* sp. Other trees present in the association belong to the Bombacaceae, Anacardiaceae, Polygalaceae, *Hedyosmum*, *Miconia* and various species of Palmae. Gramineae and Cyperaceae are in the places usually inundated by the river.

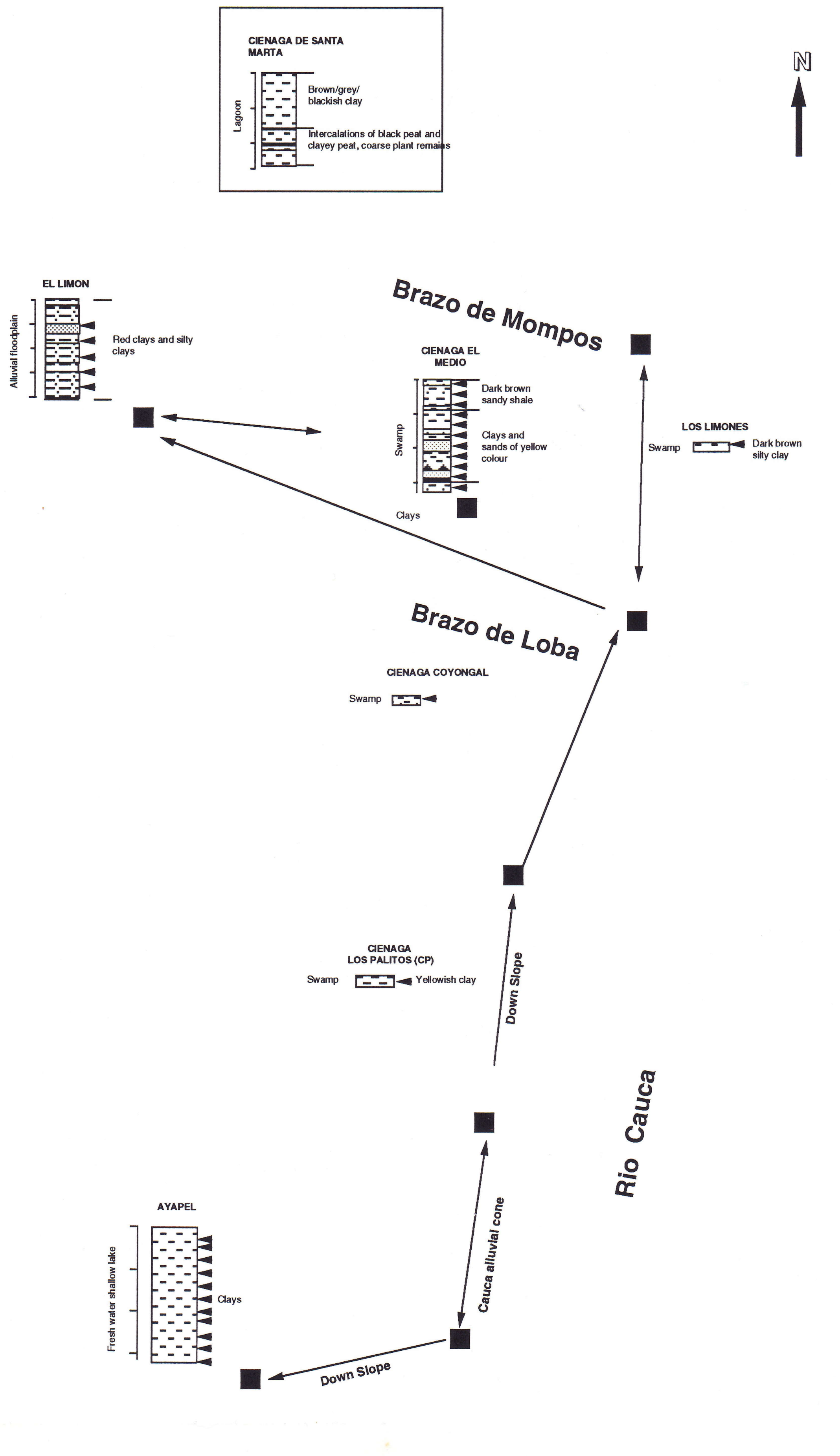


Figure 1: Studied samples and sections, Lower Magdalena River Basin, Colombia

The vegetation belts around the cienagas are composed of:

- Inner belt, where the water is deeper with: *Nymphaea*, *Limnanthemum*, *Trapa* and *Cabomba.*

-First vegetation belt landward, partly continuously inundated and partly temporarily inundated with: *Eichhornia crassipens* and *Pistia stratiotes*. It can also be founded *Salvinia*, *Marsillia*, *Ludvigia* (Jussiaea) and *Polygonum*.

-Second vegetation belt landward, vegetation inundated during greater part of the year, characterized by grasses and sedges. Getting drier gradually. Grasses may consist of one (*Paspalum*) or more species (*Eriochloa*, *Panicum*, *Leptochloa*, *Eragrostis*, *Echinocloa* and of the Cyperaceae *Cyperus*, *Ligularis* and *Imperata contracta*. With more consolidated soils and mineral- water-management improved appear *Hymenache*, *Gynerium sagittatum* and *Tessaria integrifolia*.

- Third vegetation belt landward, consists of brush- and tree-vegetation

- Fourth vegetation belt landward, where the ground-water-level is at its lowest, trees dominated. Present in the association are: *Cecropia*, *Ficus*, *Celtis*, *Psychotria*. Also can be found *Zygia*, *Inga* and other Mimosaceae.

Plants like *Salix humboldtium*, *Alnus* and *Myrica* only occur in the upper course of the Magdalena river, in the Andes where they grow between 2000 and 3500 m.

#### - PALYNOLOGY AND PALYNOFACIES OF RECENT ENVIRONMENTS

#### The Ayapel Section

##### Location and general information

Ayapel is located in the swampy depression ("depresion cenagosa") very close to the right bank of the San Jorge River, between San Jorge River and Cauca River in the vicinity of the localities of "Ayapel" and "El Cedro". ( app. borehole coordinates: 75º03'W - 8º24'N). Numerous water courses and creeks drain the area, among them: Caño Muñoz, Caño Cedro, Caño Aguas Claras, Quebrada Escobillas and Quebrada Quebradona .

Ayapel is situated within the Palmitas area, which has an average sedimentation rate of 2.7mm/year. The lake with an irregular shape has a main axis of about 25 Km length  and a total area of water coverage about 250 km2.

**Lithology and sample description** (fig. 2)

A total of 11 samples were studied (sample depth shown in m):

0.25 : Silty clay, light gray colour, with oxidation spots and roots remains

0.45 : Silty clay, light gray colour, with oxidation spots

0.75 : Silty clay, light gray colour, with oxidation spots

1.05 : Silty clay, light gray colour, with oxidation spots and plant remains

1.35 : Silty clay, light gray colour, with oxidation spots

1.65 : Silty clay, light gray colour, with oxidation spots and root casts

1.95 : Silty clay, light gray colour, with oxidation spots

2.25 : Silty clay, gray colour, with oxidation spots and abundant plant remains

2.55 : Silty clay, light gray colour, with oxidation spots

2.85 : Silty clay, light gray colour, with oxidation spots

3.20 : Silty clay, light gray colour, with oxidation spots and thin layers of plant remains.

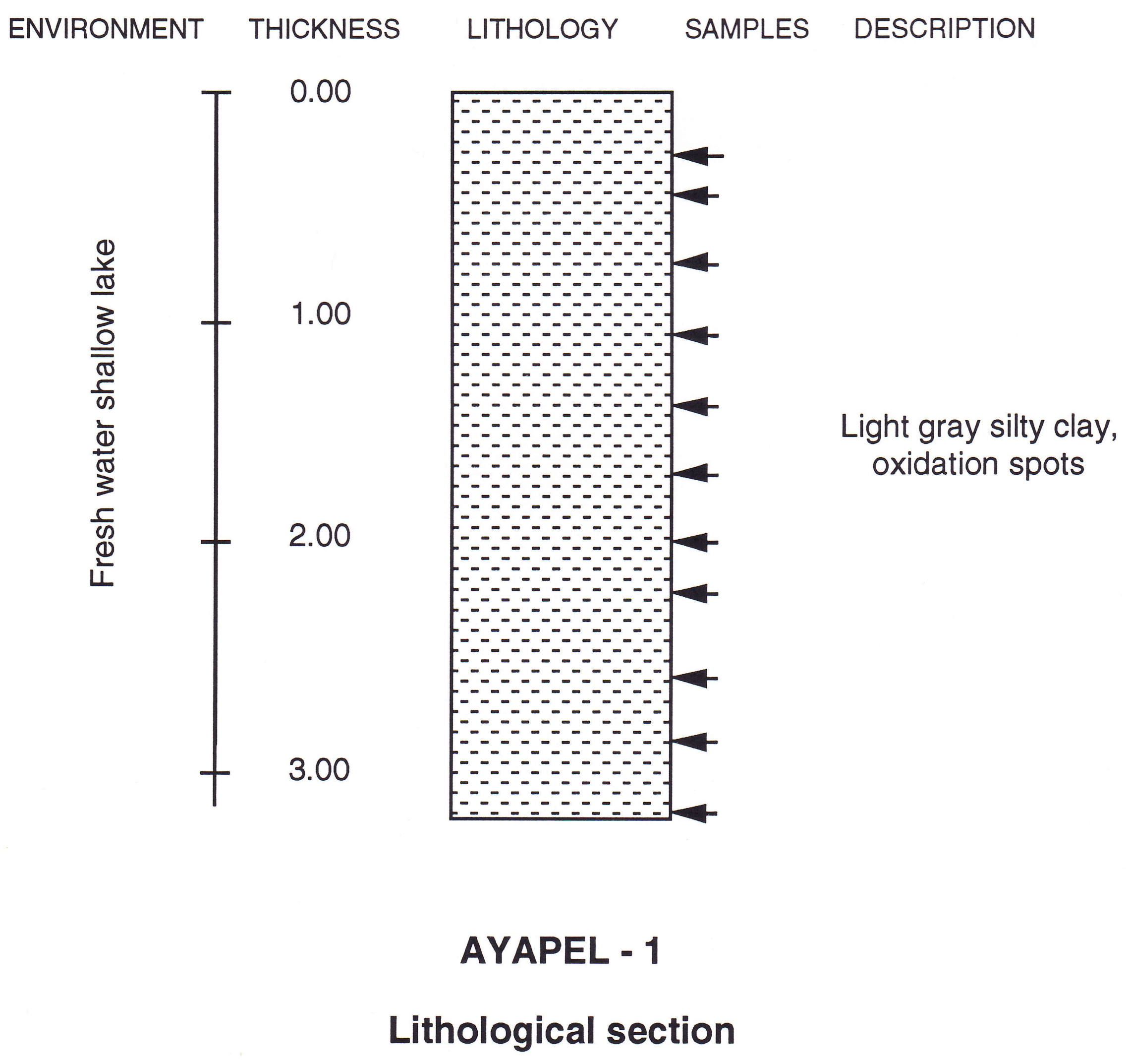


Figure 2: Lithology and sample position Ayapel core section

**Palynology**

Ayapel samples show variations in the sporomorph content (fig 3). The flora is in general richer in specimens between 1.65 and 0.45 m than between 3.20 and 1.95 m. The relative low value founded at 0.25m is maybe due to the diluting effect of more abundant organic matter at that depth.

- Pollen and spores & ecological groups diagram (fig. 3 a & b)

The flora of Ayapel section from 2.55 m - 2.85m, close to the base of the section, shows a pollen diagram with grass pollen dominant over composites while pollen from fresh water/floating plants and palms are present in significant amounts.

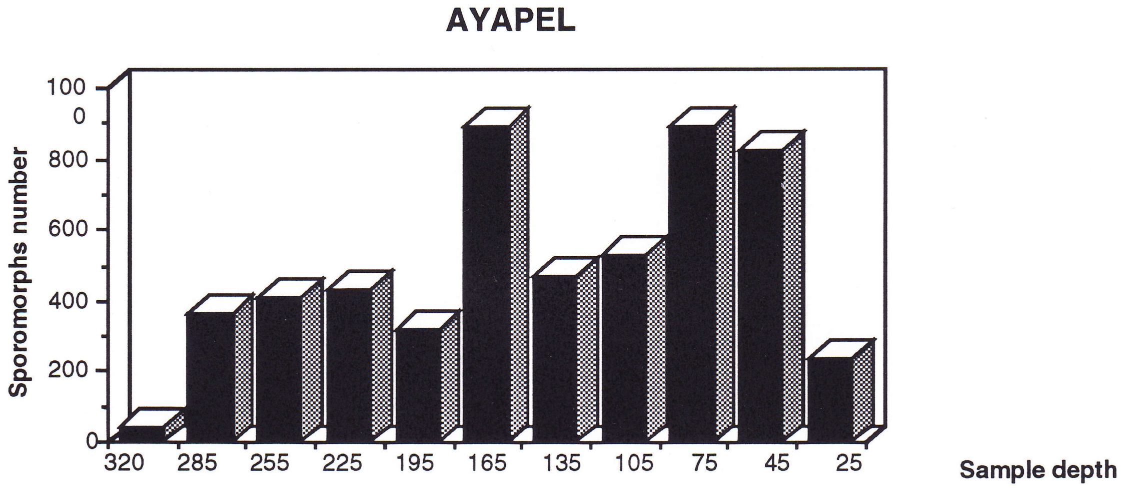


Figure 3: Sporomorph concentration. Sporomorphs /10µl.

The deepest sample (3.20 m) is very poor in sporomorphs.

From 2.25 m to 0.25 m the flora is dominated by composite and grass pollen. From 1.65 m upwards the abundance of sporomorphs increases (fig. 3). Pollen from floating and water plants is present throughout the interval but in minor concentrations.

- Palynomorphs diagram (fig.3c)

In general, sporomorphs tend to be more abundant than fungal remains in this section. Fresh water algae (including *Botryococcus* sp.) are present throughout the core. From 1.65 m upwards the amount of fungal remains increase, as well as the amount of fresh water algae .

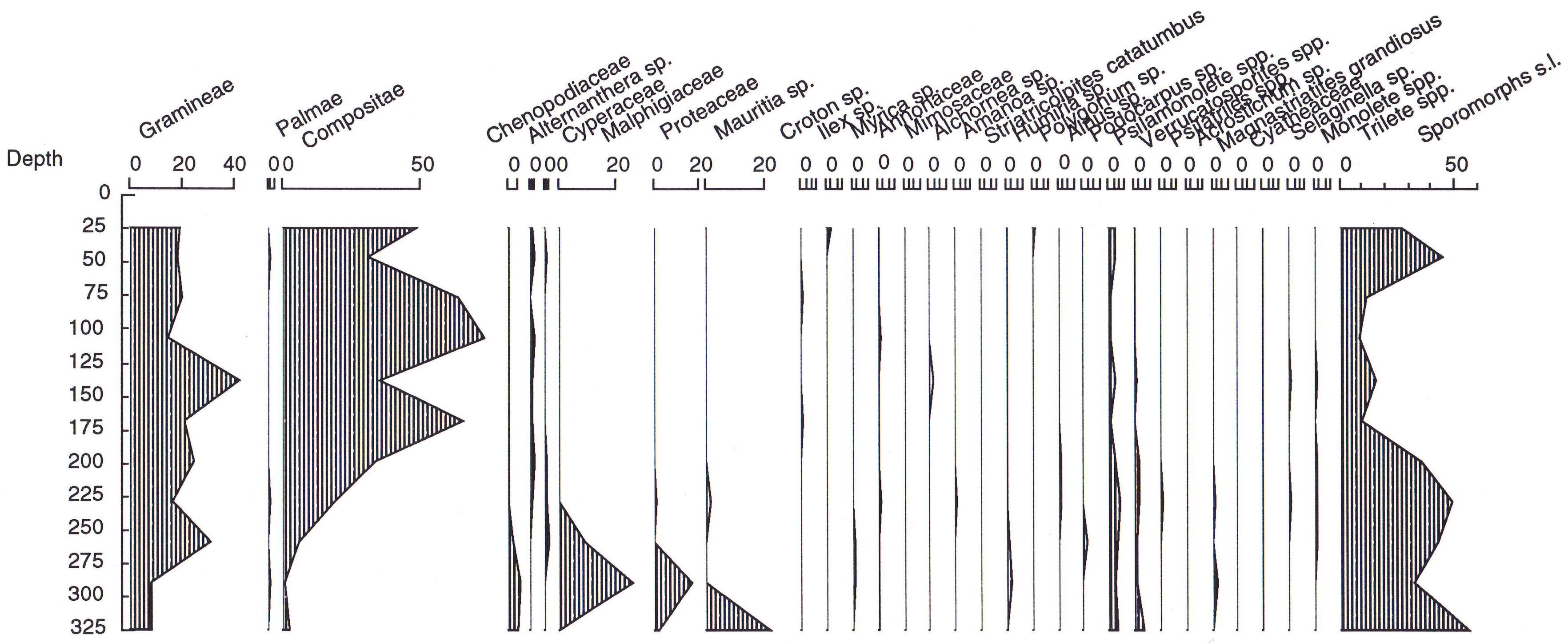


Figure 3a: Sporomorphs diagram. Ayapel section

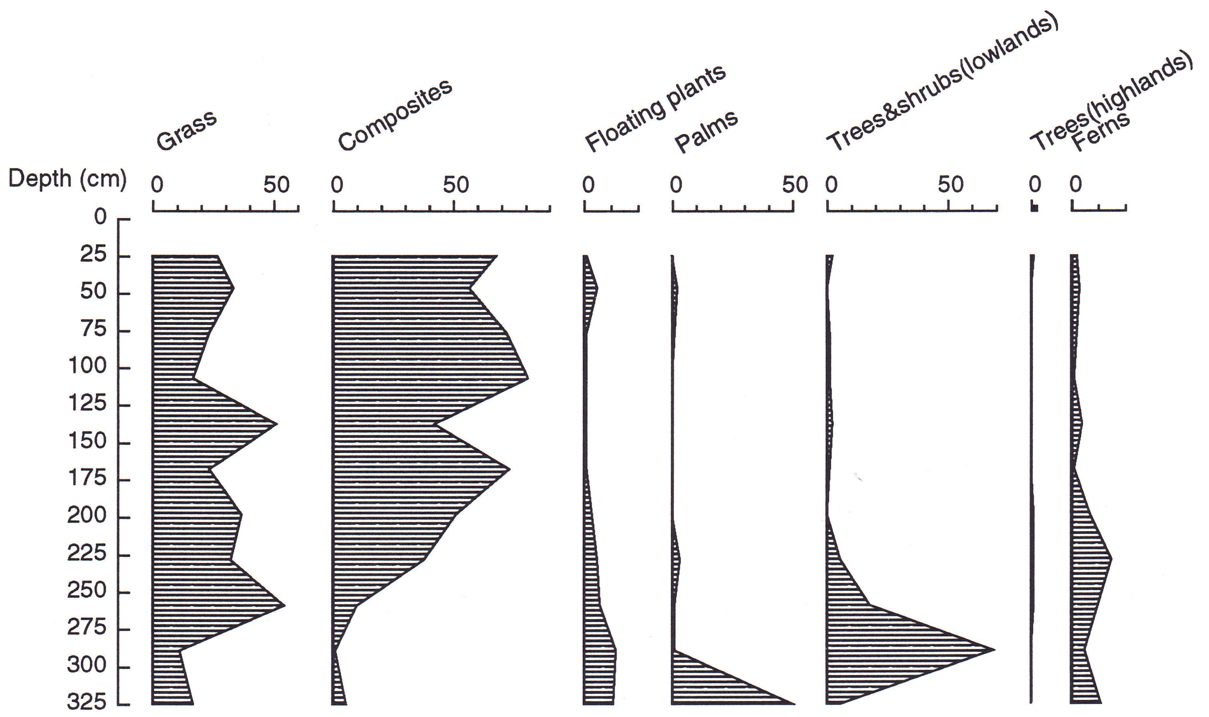


Figure 3b: Paleoecological groups diagram. Ayapel section

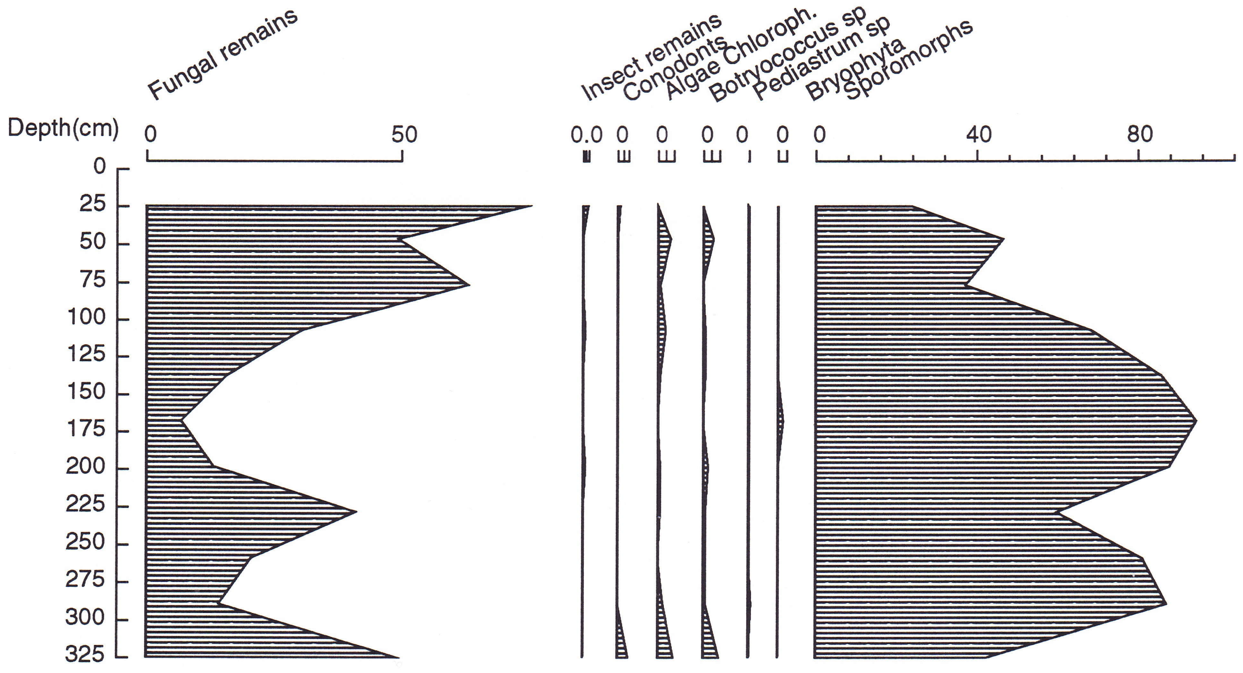


Figure 3c: Palynomorphs diagram. Ayapel section

**Palynofacies**

- Organic matter concentration (fig.4a):

The organic matter concentration is in general low with values below 0.1%, with the only exception of sample 0.25 m (the closest to the surface) with slightly more than 0.2%.

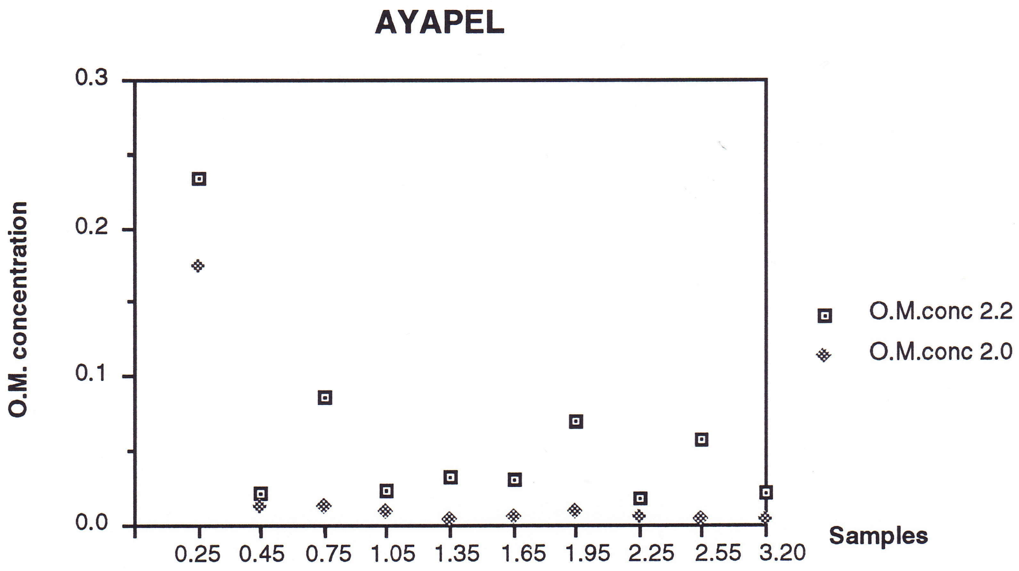


Figure 4a: Organic matter concentration as measured after 2.2 and 2.0 heavy liquid separation.

- Organic matter composition (fig. 5)

Samples from 1.05 m to 3.20 m (3.20 m is very poor) are dominated by amorphous opaque materials. Palynomorphs,woody, other plant tissues, fungal remains and other amorphous are present but in very low concentrations (fig. 5)

Sample 0.75 has very abundant amorphous homogeneous, amorphous opaque and fungal remains.

Sample 0.45 m has amorphous heterogeneous and amorphous finely dispersed as main materials, with minor amounts of amorphous opaque, light coloured plant tissues and palynomorphs.

Sample 0.25 m is different from the rest, with very abundant light coloured plant tissues as epidermal/cut. and other plant remains. Present in lower concentrations are opaque amorphous materials, palynomorphs, animal, fungal and woody remains. The finely dispersed organic matter is very abundant .

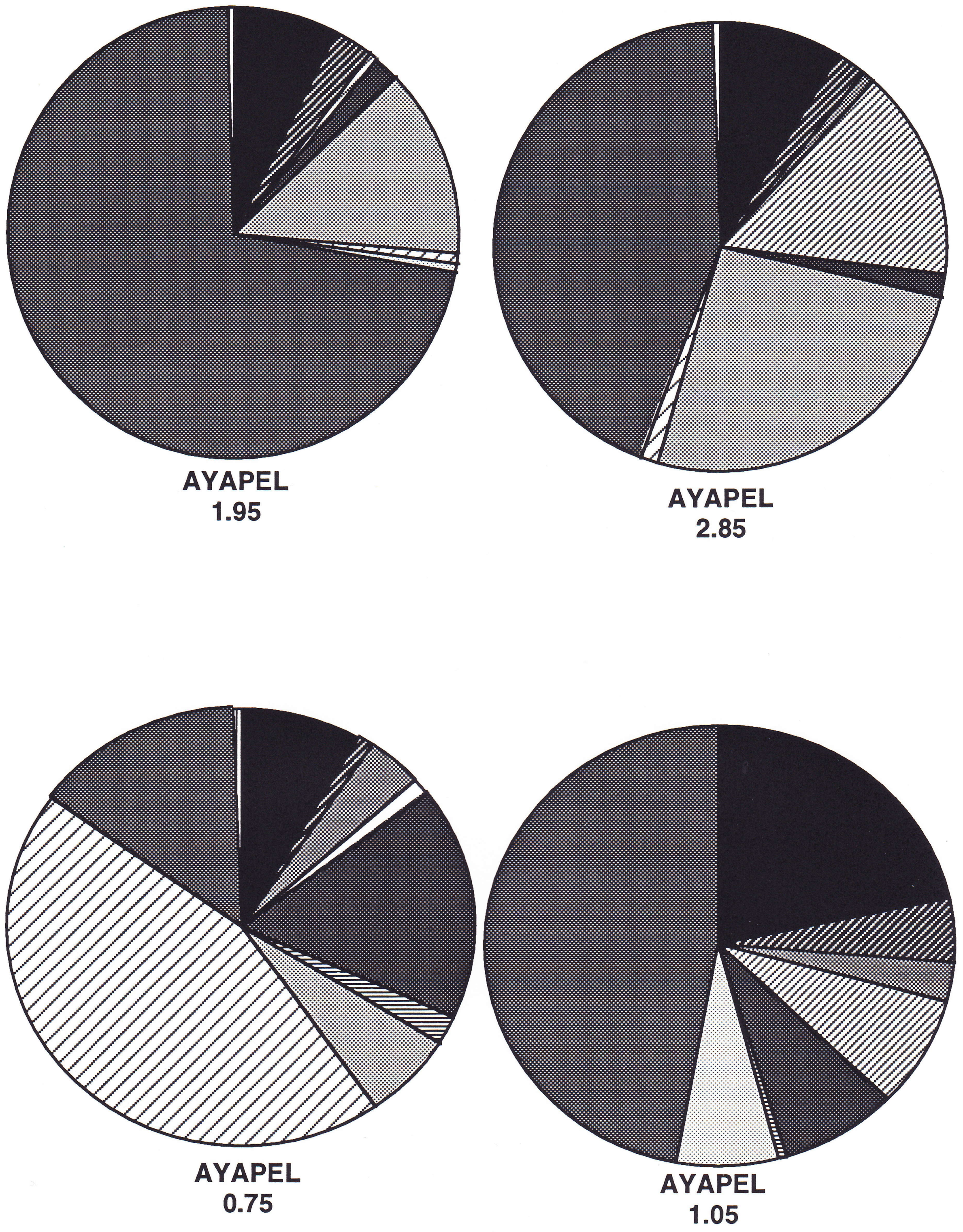


Figure 5: Organic matter composition.Selected samples.
Ayapel section

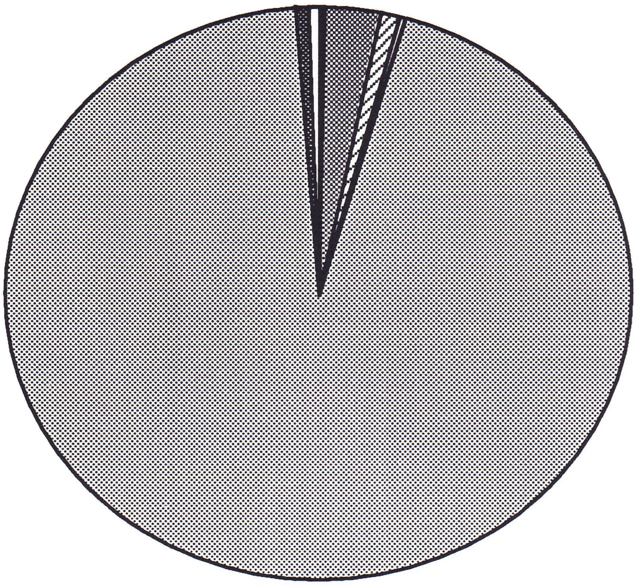

**AYAPEL**
**0.25**

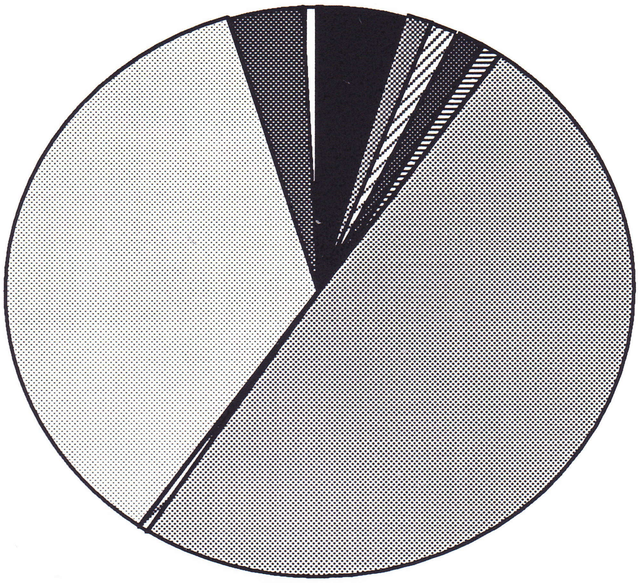

**AYAPEL**
**0.45**

#### Legend

**(Amsterdam Palynological Organic Matter Classification, 1991)**

- Palynomorphs
- Woody
- Epidermal/Cuticular
- Other plant remains
- Animal remains
- Fungal remains
- Opaque structured debris
- Finely dispersed amorphous
- Amorphous homogeneous
- Amorphous heterogeneous
- Amorphous opaque
- Indeterminated opaque
- Indeterminated others (organo-mineral gel)

Figure 5: Organic matter composition.Selected samples. Ayapel section

- Organic matter morphological characterization (fig. 6)

The section comprised between 3.20 m and 0.45 m is characterized by a maximum in the sum of areas grain size distribution mainly in ø5 or rarely together with ø4 (medium to coarse silt equivalent size).

Relative elongation histograms show that more than 50% of the particles are equidimensional, but in general between 15% to 20% have clearly elongate shapes.

The sphericity histogram for samples from 3.20 m to 0.75 m (only exception sample 2.85 m) shows a bimodal distribution with a first maximum at 0.1 and a second maximum at 0.5 or 0.6.

Sample 0.25 m has different morphological characteristics, with about 80% of the total analyzed area covered by particles within the ø3 to ø1, medium to very fine sand, size range. Numerically more than 55% of the particles belong to classes ø7 to ø 4, silt size range. Class ø8 (clay size materials) has about 45% of the particles but cover only 2% of the total analyzed area.

More than 60% of the particles fall in the equidimensional shapes, but there is a relatively important 15%, that clearly have elongate to strongly elongate shapes.

Sphericity values show that particles in the 0.0 and 0.1 classes are the most abundant with more than 60% of the particles comprised within these two classes.

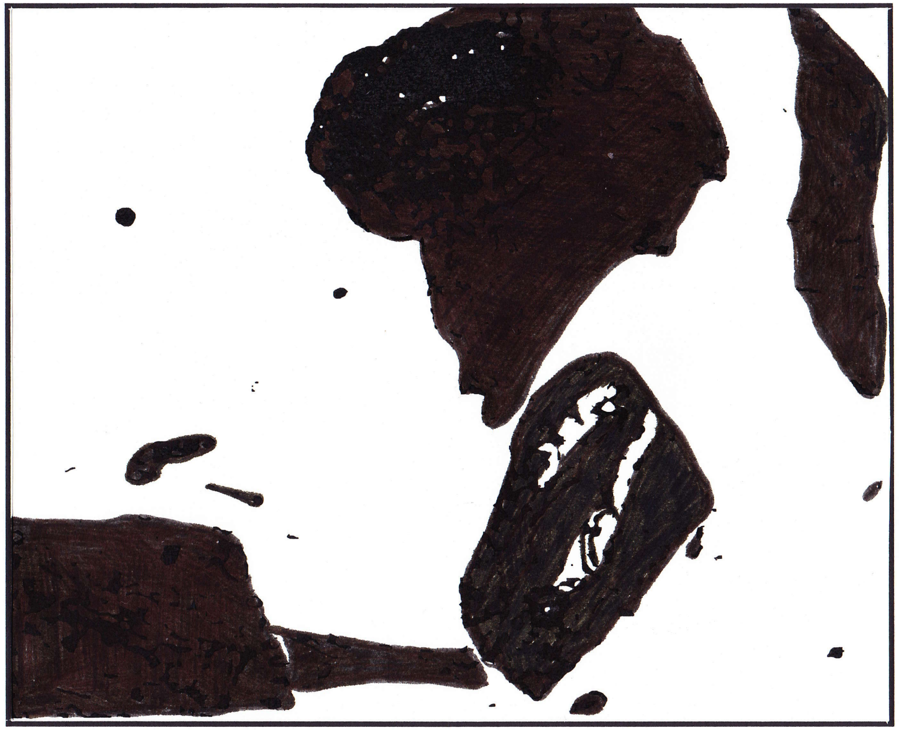

AYAPEL 0.25

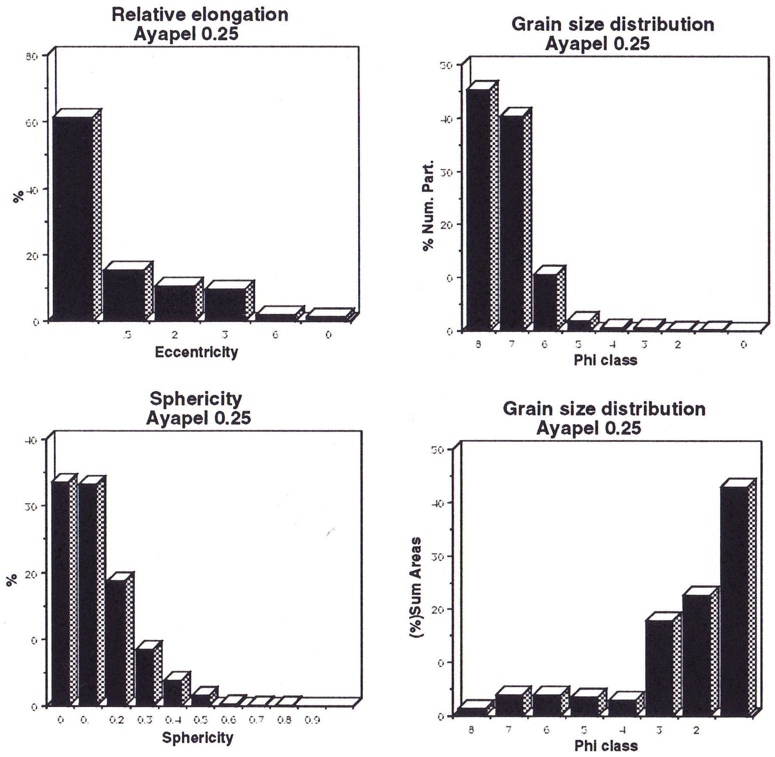


Figure 6: Organic particles morphological characterization. Selected samples from the Ayapel core section

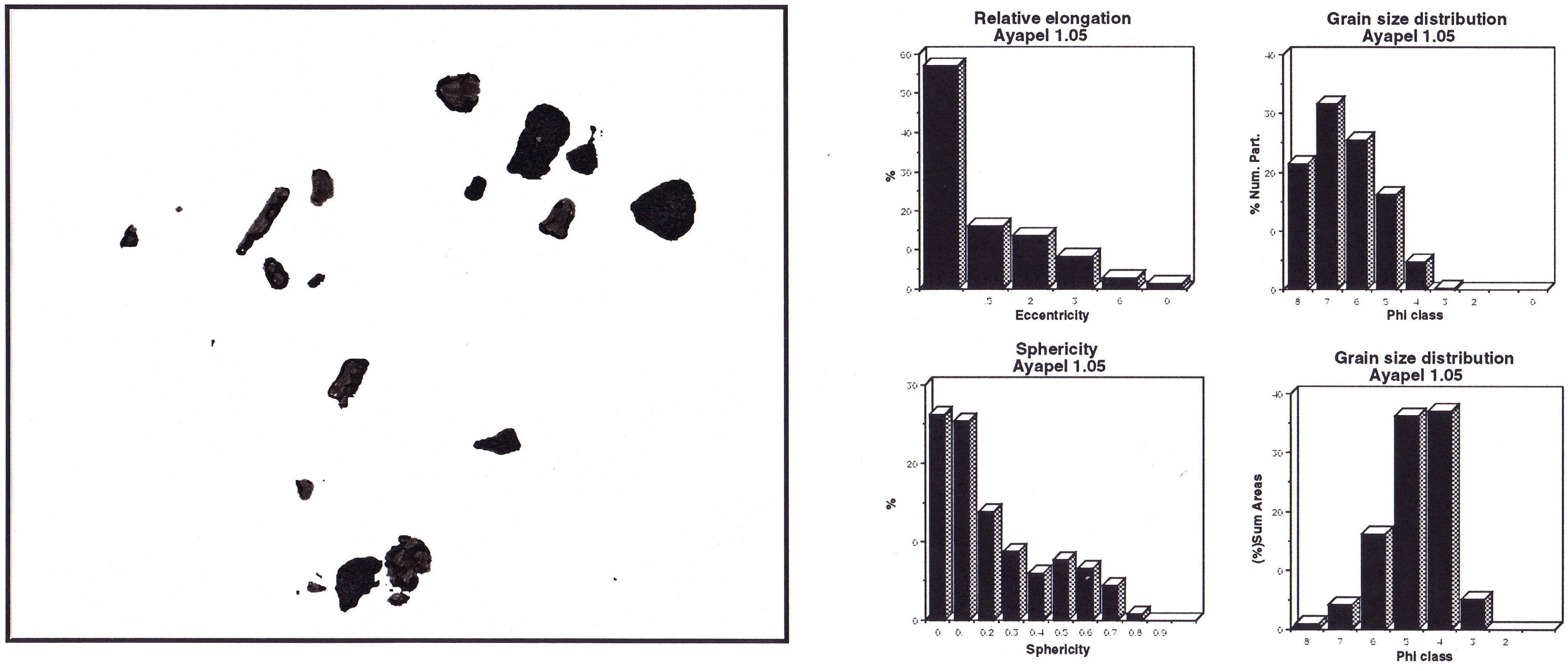


AYAPEL 1.05

Figure 6: Organic particles morphological characterization. Selected samples from the Ayapel core section

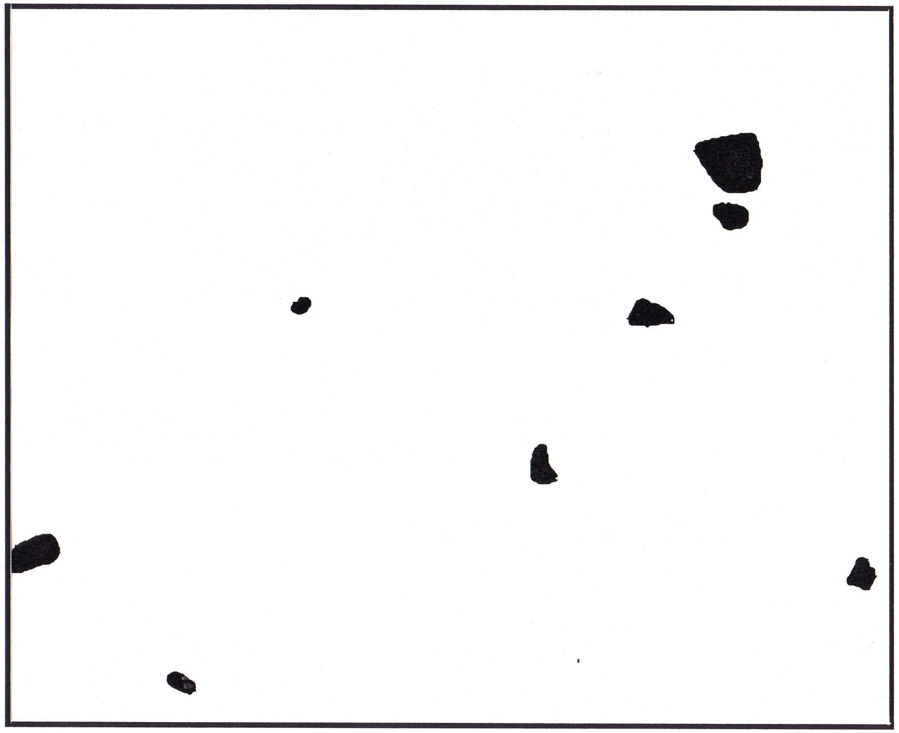

AYAPEL 1.95

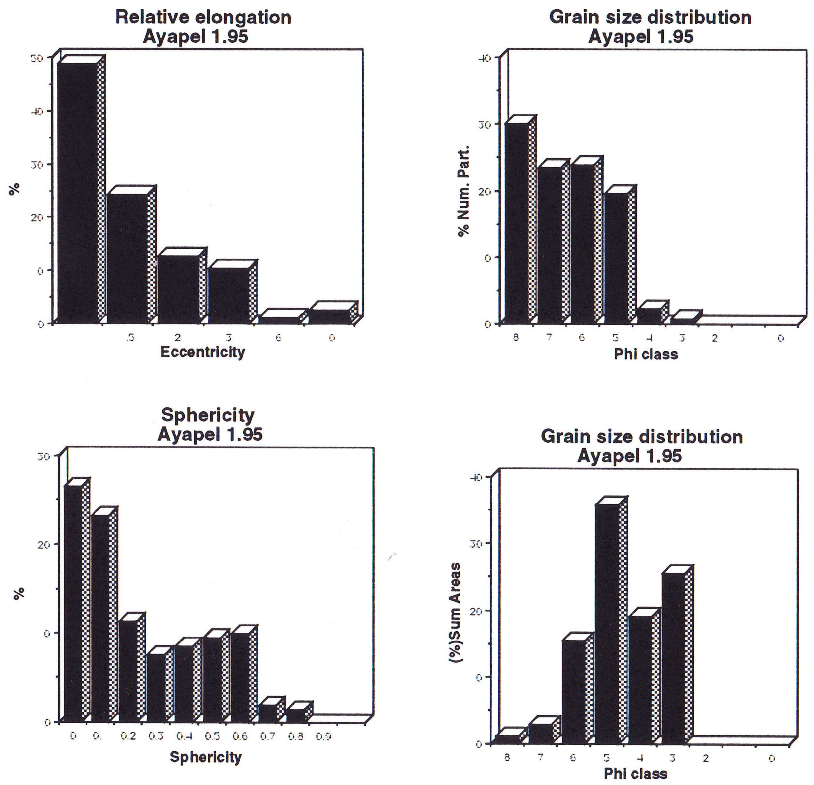


Figure 6: Organic particles morphological characterization. Selected samples from the Ayapel core section

#### The "Cienaga de El Medio" (CM) Section

##### Location and general information

The "Cienaga de El Medio" is situated to the north of Caño Violo, between "Las Boquillas" and "Candelaria" (borehole coordinates: 74º30'W - 9º08'N), in the swampy depression, strongly influenced by the Magdalena River flooding. Water stays for long periods covering the area with an average thickness of 3 m during the high stand season.

The sedimentation rate for the Boquillas area are according to H.I.M.A.T. (1977):
in average 2.92 mm/year; maximum value 4.0 mm/year for the period 1941 - 1975.

##### Lithology and sample description (fig.7)

A total of 11 samples were studied (sample depth in m):

- Surface: Silty clay, medium gray colour, with oxidation spots and root remains
- 0.15: Silty clay, medium gray colour, with oxidation spots, irregular fracture
- 0.45: Sandy clay, light gray colour, with abundant oxidation spots, irregular fracture
- 0.75: Sandy clay, light gray colour, with abundant oxidation spots, irregular fracture
- 1.05: Sandy clay, light gray colour, with abundant oxidation spots, irregular fracture
- 1.35: Shaly silt, light gray colour, with oxidation spots, some micaceous material
- 1.65: Sandy clay, light gray colour, with abundant oxidation spots, irregular fracture
- 1.95: Silt to fine sand, micaceous, light gray to brownish, with a lot of oxidation spots
- 2.25: Sandy clay, light gray colour, with oxidation spots and plant remains,micaceous
- 2.55: Clay, light gray colour, with small oxidation spots and carbonaceous material.
- 2.85: Silty clay, light gray colour, with some oxidation spots.

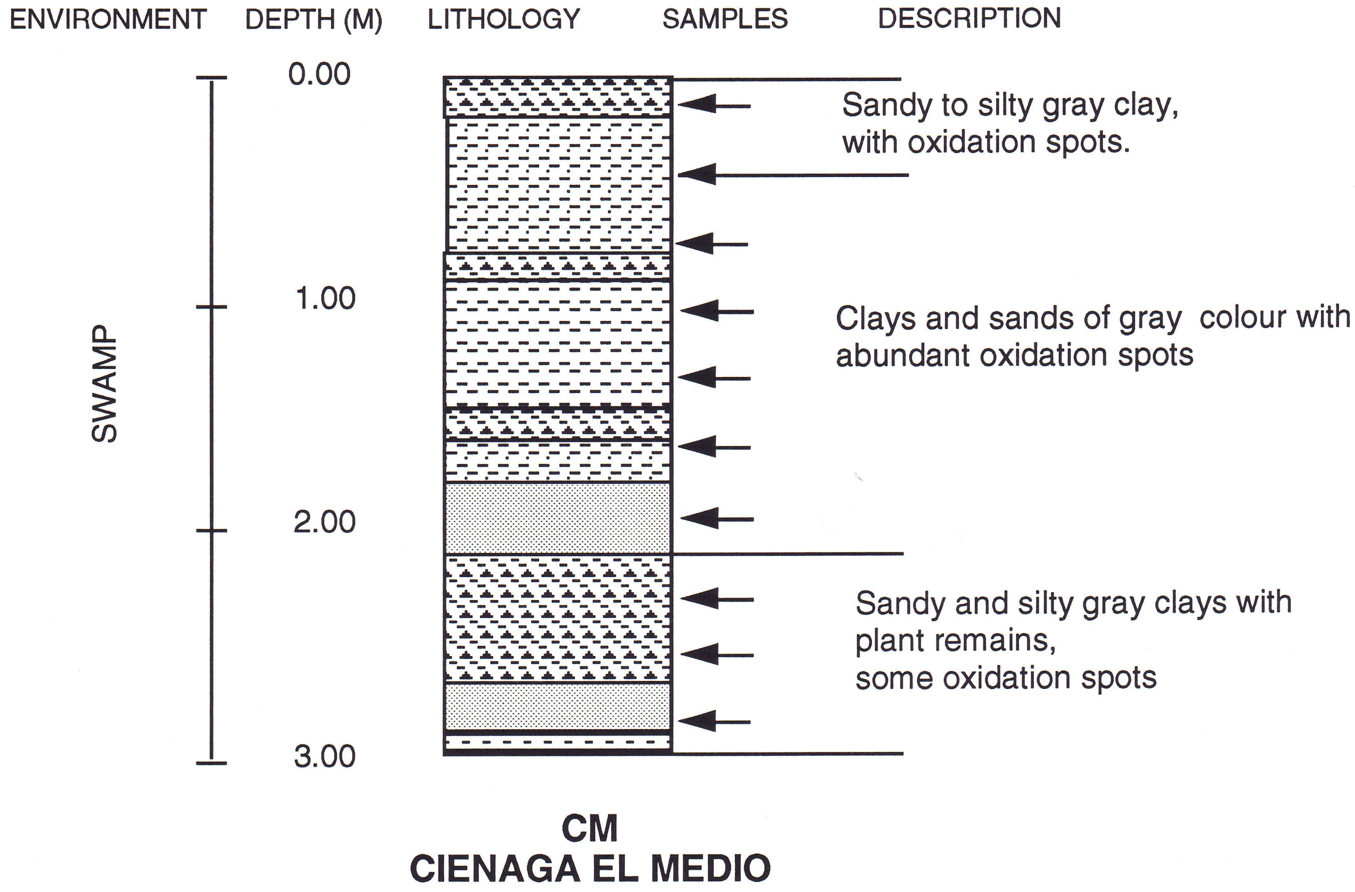


Figure 7: Lithology of the Cienaga de El Medio core section

##### Palynology

The sporomorph concentration has strong variations in the Cienaga de El Medio section (fig. 8). The richest interval was found between 2.85 m and 2.25 m, with a sharp decrease upwards in the section. These decrease is closely reated with the indications of higher degrees of oxidation shown on the lithology.

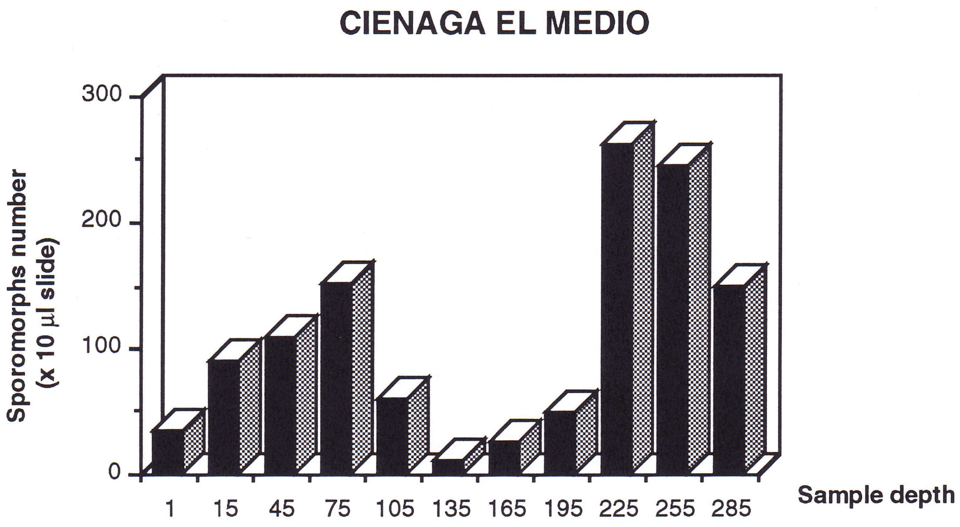


Figure 8: Sporomorph concentration. Absolute amount of sporomorphs / 10µl slide.

- Pollen and spores & ecological groups diagram (fig.8a &b)

The pollen and spore assemblages are in general poor in species and specimens. The richest interval is comprised between 2.85 m and 2.25 m and is dominated by grass pollen and composites alternatively.

Upwards in the section from 1.95 m to 0.45 m the amount of specimens of composites pollen strongly decreases, while gramineae pollen together with fern spores (fig. 8b) are the main components of the assemblage.

Chenopodiaceae, Cyperaceae and *Mauritia* sp. show relative increases. From 0.15 m to the surface, pollen grains of composites become dominant.

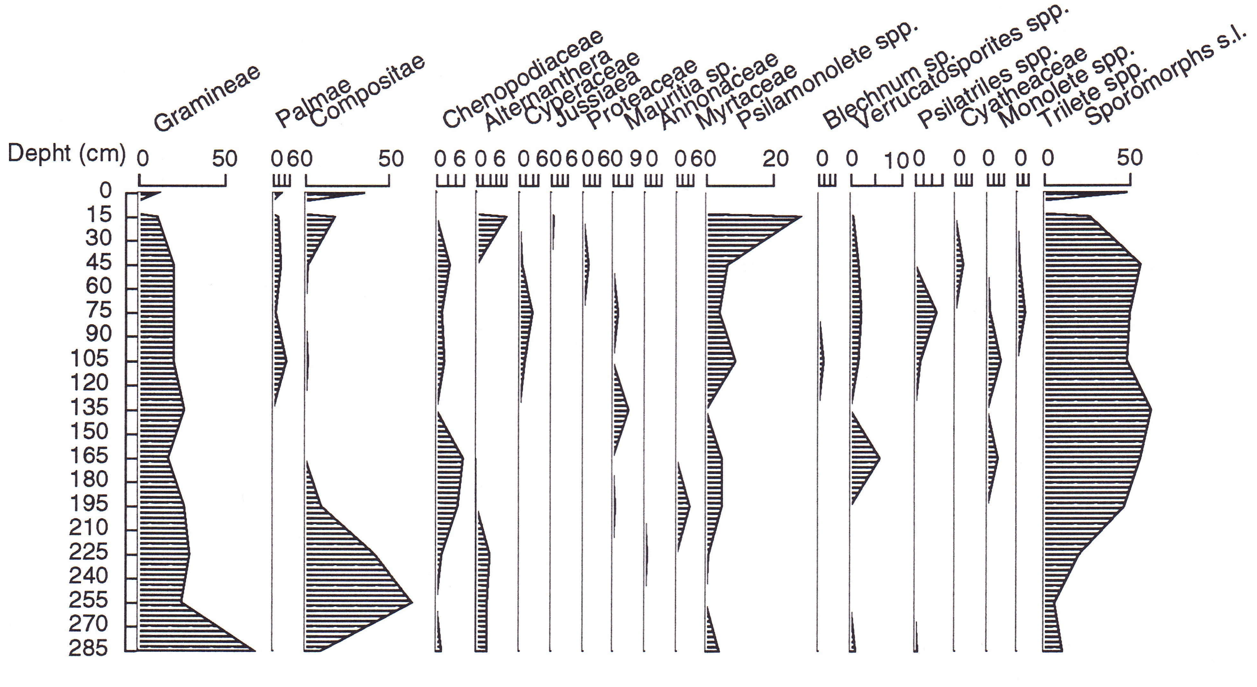


Figure 8a: Sporomorphs diagram. Cienaga de El Medio (CM) core section

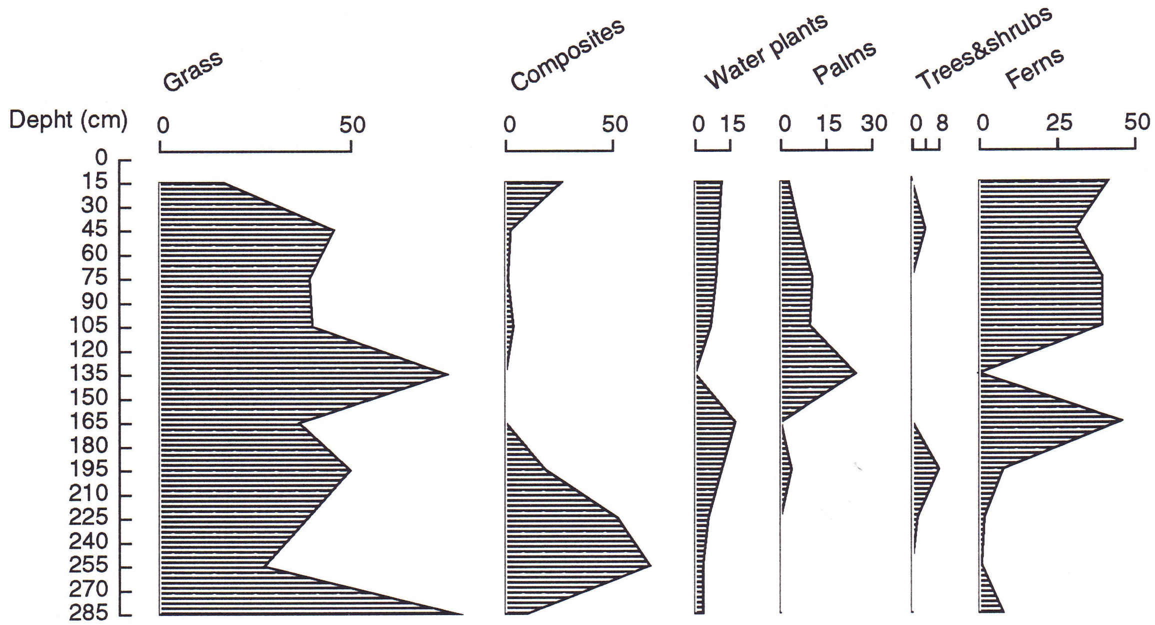


Figure 8 b: Groups diagram. Cienaga de El Medio core section

- Palynomorphs diagram (fig.8c)

The palynomorphs diagram shows that fungal remains are at least as abundant as the sporomorphs in Cienaga de El Medio although the relative abundance varies from one sample to another. The presence of fresh water algae, including *Botryococcus* sp. is constant through out the interval. Spores of bryophyta as well as conodonts and insect remains are restricted only to the upper part of this section.

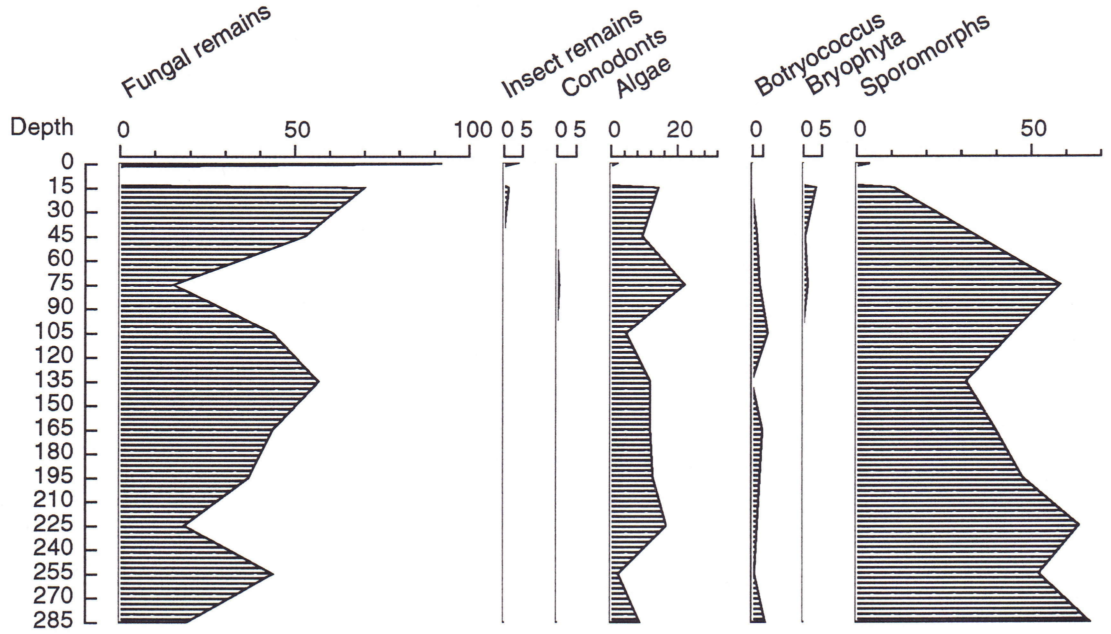


Figure 8c: Palynomorphs diagram. Cienaga de El Medio core section

#### Palynofacies

- Organic matter concentration (fig.4b)

Organic matter concentration in Cienaga de El Medio is quite low, almost always below 0.1%, with the only exception of sample 0.15 m with about 0.14 %.

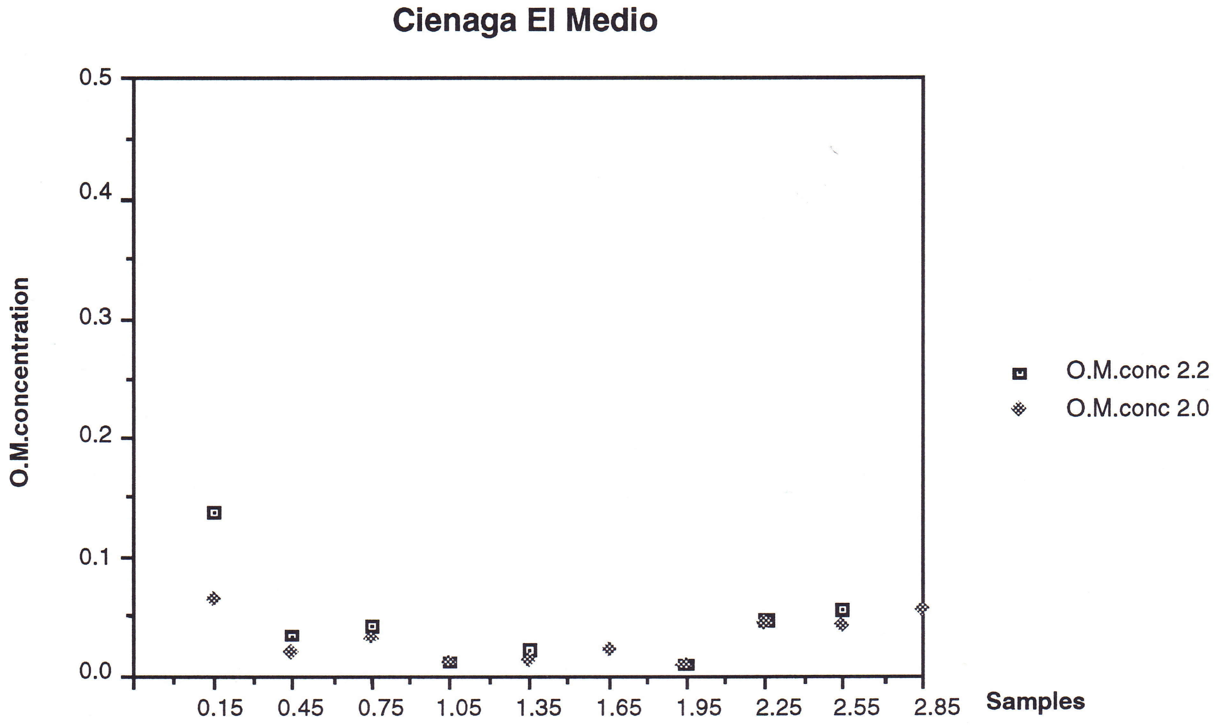


Figure 4b: Organic matter concentration as measured after 2.2 and 2.0 heavy liquid separation

- Organic matter composition (fig.9)

In the Cienaga de El Medio section, changes on organic matter composition follows closely the levels of relative oxidation as they are reflected on the lithology.

From 2.85 m to 2.25 m the material is mainly composed by very fine debris that were classified as amorphous finely dispersed. The rest of the material is mainly composed of “organo-mineral” gels, amorphous opaque, woody, other plant remains and palynomorphs as shown in figure 9a (after removing the finely dispersed amorphous).

From 1.95 m to 1.05 m, the level of oxidation of the material increases, related with this the amount of amorphous opaque and “organo-mineral gels” rises in a significant way, most of the time they account together with the finely dispersed amorphous for more than 85% of all the materials preserved.

From 0.75 m to 0.15 m the organic matter is entirely dominated by amorphous finely dispersed and in a lower proportion by amorphous opaque and "organo-mineral gels "type. The main difference between these set of samples and the set between 1.95 m and 1.05 m is the concentration of the finely dispersed amorphous organic matter. When the rest of the components are compared there is not a significative difference in composition as shown in figure 9b.

The surface sample has a different composition from the rest of the section, with abundance of light coloured plant tissues, and some amount of fungal and opaque materials.

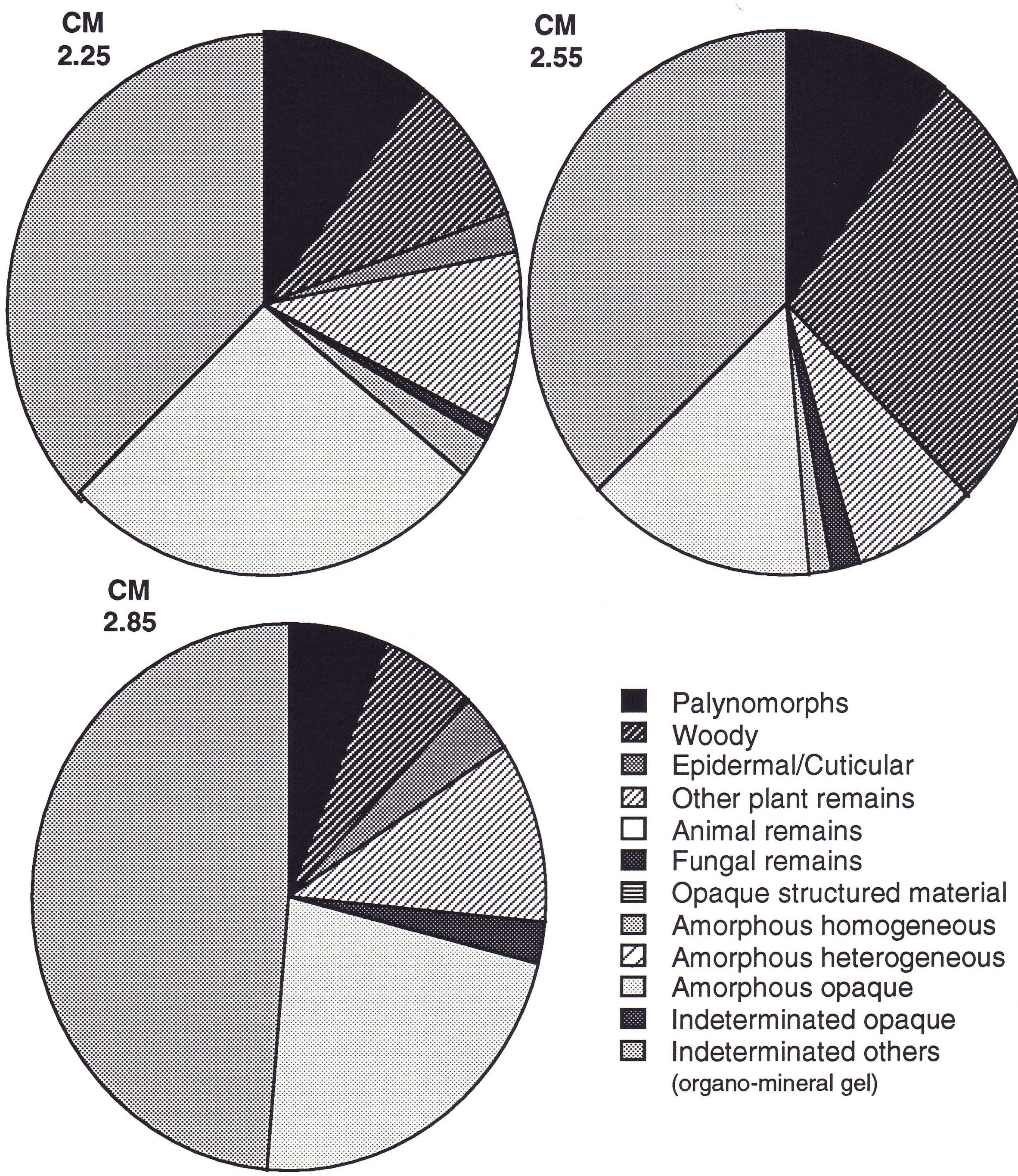


Figure 9a: Organic matter composition. Ciena de El Medio core section

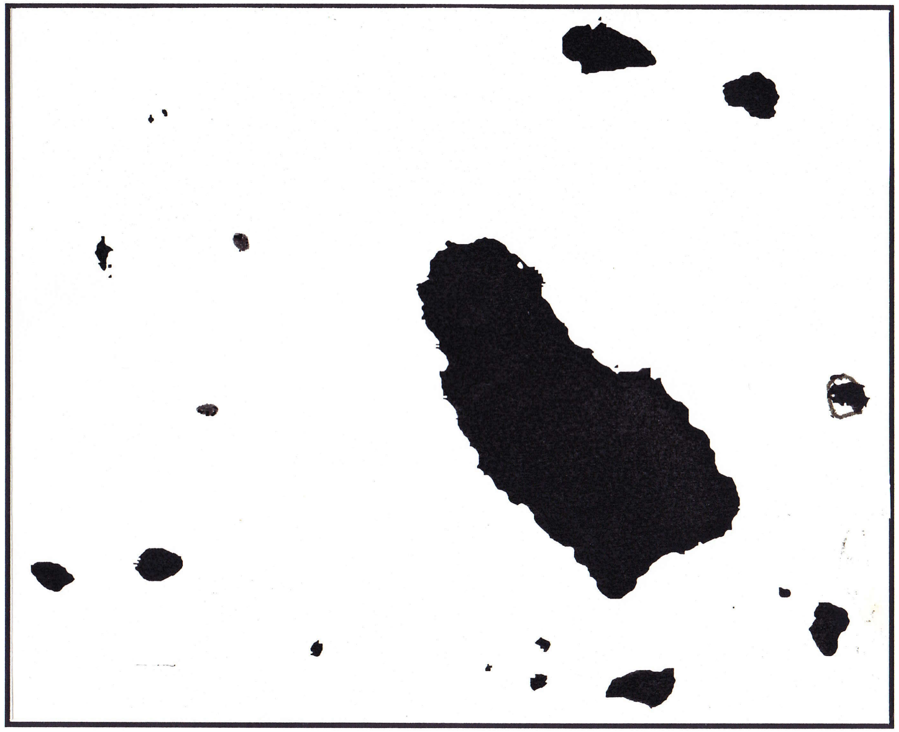

CM 2.25

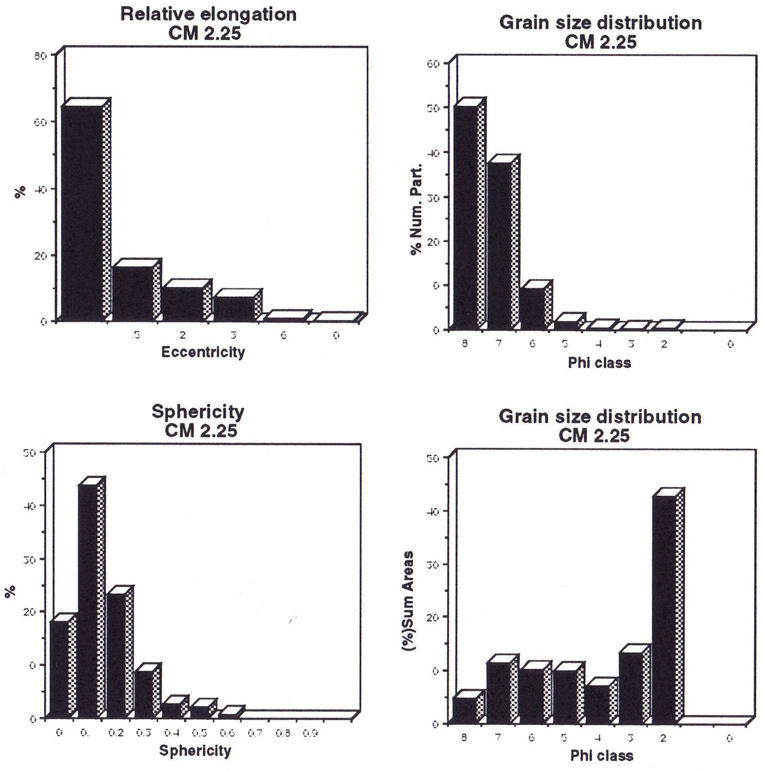


Figure 10: Organic particles morphological characterization. Selected samples from the Cienaga El Medio core section

- Organic matter morphological characterization (fig. 10 and 11)

Grain size of fragments varies mainly between ø 1 and ø8 or medium sand to clay size range. In general the sum of areas grain size distributions have a maximum in the ø 2, 3 or 4 range . Numerically the most abundant particles are in the very fine silt and clay size range. Grain size distribution is related with the composition of the organic matter that in this samples is dominated by finely disperse organic matter and/or by "organo-mineral gels ".

Tree different intervals can be recognized according with the characteristics of the grain size distributions: 1.- 2.85 m to 2.25 m.(fig. 10), 2.- 1.95 m to 0.15 m and (fig. 11) and 3.- Surface (fig. 11 cont.)

The 2.85 m to 2.25 m interval is characterized by small particles (clay size range, ø 8) that although numerically are close to the 50% cover less than 5% of all the particles area. Particles in the categories of silt grain size (ø7 to ø4) account numerically for almost 50% of the total and each category for about 10% of the total particles areas. The exception is the 2.85 m sample were they are more important due to the lack of particles in the size range of medium to very fine sand, ø1 to ø3. Large particles, ø1 and ø2, seldom occur, but due to its size they cover large areas and for that reason they migth seem over represented on the sum of areas diagram.

Between 50% and 65% of particles are equidimensional, with less than 5% in the strongly elongate to filiform shapes.In all samples more than 75% of the particles fall in the sphericity range from 0.0 to 0.20, but it varies up to 0.8. The main peak occurs at 0.1.

The 1.95 m to 0.15 m interval is generally characterized by the numerical dominance 42 to 50% of very fine silt size range, ø7, particles. The ø7 size covers not less than 15% of the particles area and commonly more than 20% of it. Each category between ø6 and ø3, fine silt to very fine sand, cover between 10% and 25%. All this shows a relative increase in particle sizes when compared with the previous interval, this is possibly due to the agglutinated effects of the "organo-mineral gel" particles.

Most of the samples have 52% to 60% of the particles in the equidimensional range, the relative elongation histogram show a step-like distribution. More than 70% of the sphericity values falled on the 0.0 to 0.2 range but it varies up to 0.7.The main peak occurs at 0.1 as in the previously described interval.

The surface sample is numerically dominated by particles in the silt size range (ø7). Particles comprised between ø8 and ø3, clay to very fine sand, cover less than 50% of the particles area with an average of less than 7% per category, whereas particles with sizes larger than ø2 cover more than 50% of the particles area. The relative elongation has a step-like distribution similar to interval 1.95 to 0.15. The sphericity main peak occurs at 0.1 as in the previously described intervals.

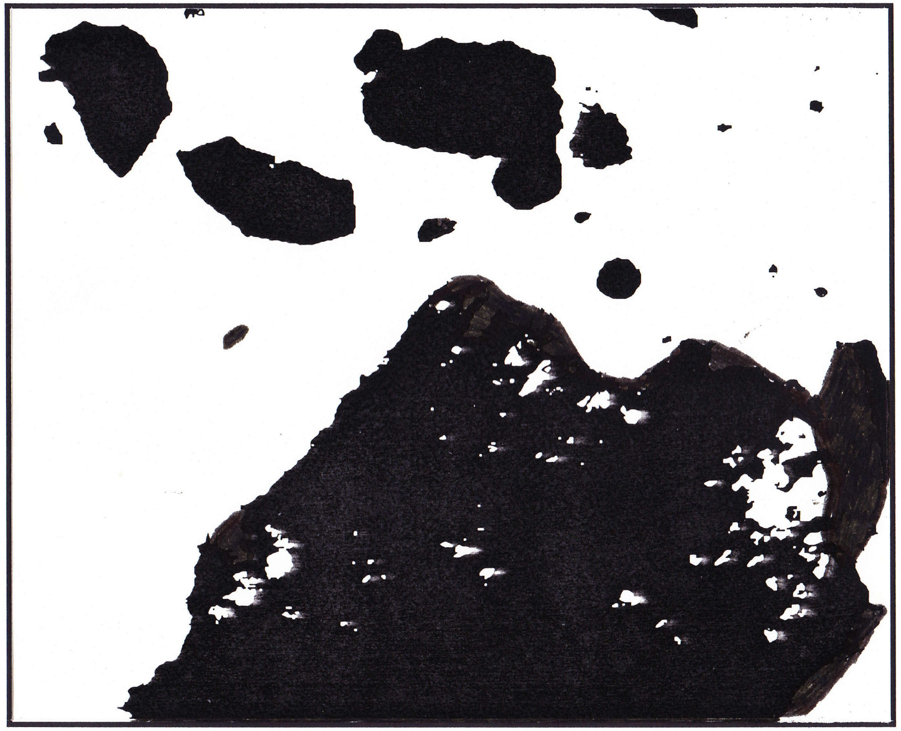

CM 0.45

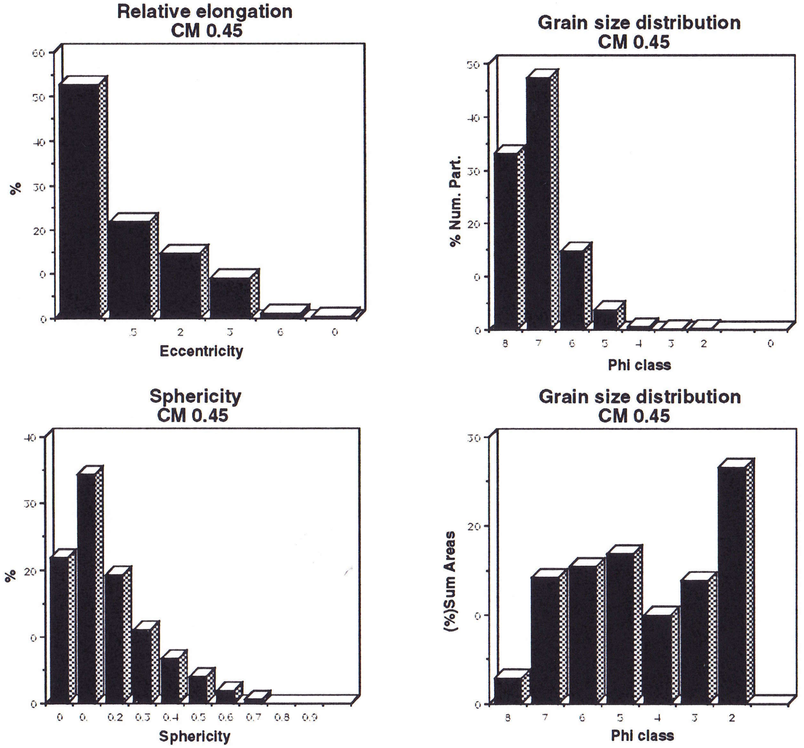


Figure 11: Organic particles morphological characterization. Selected samples from the Cienaga El Medio core section

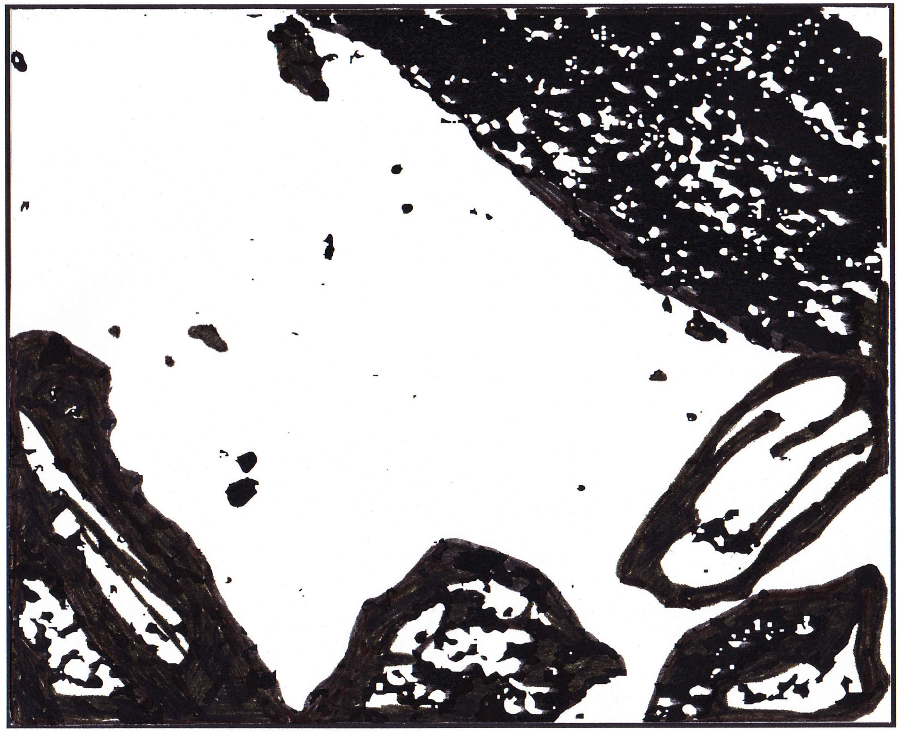

CM Surface

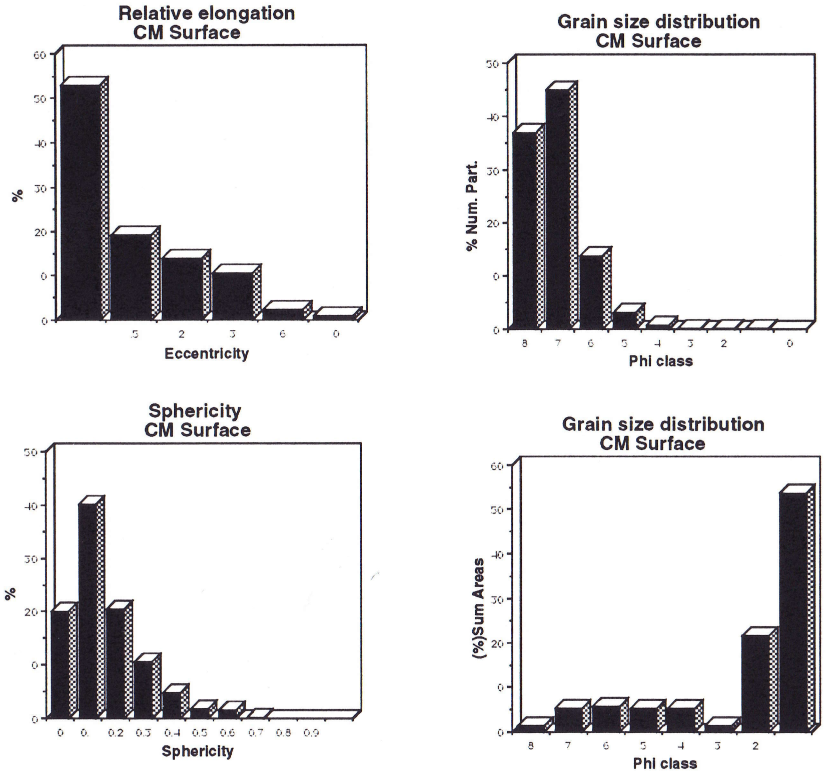


Figure 11: Organic particles morphological characterization. Selected samples from the Cienaga El Medio core section

#### The El Limon Section

##### Location and general information

The El Limon core was taken in an alluvial flood plain, close to the El Limon town (74º39'N 9º17'W) near the intersection between Brazo de Loba and Brazo de Mompos.

##### Lithology and sample description

A set of ten samples were analyzed (fig.12):

1.05 (m) Medium to fine sand, creamy colour, with oxidation spots, non-homogeneous reddish colour.

1.37 (m) Medium to fine sand, creamy colour, with oxidation spots, non homogeneous reddish colour.

1.80 (m) Medium to fine sand, light gray colour, with oxidation spots, non-homogeneous reddish colour.

Between 1.80 m and 2.55 m it is a very poorly recovered coarse sand.

2.87 (m) Medium to fine sand, light gray colour, with oxidation spots, non-homogeneous reddish colour.

Between 3.30 m and 4.05 m there is no material available.

4.05 (m) Coarse silt to fine sand, light gray colour almost white, with oxidation bands, non-homogeneous reddish colour.

4.37 (m) Medium to fine sand, light brown homogeneous colour.

4.80 (m) Medium to fine sand, light gray colour, with oxidation spots, non-homogeneous reddish colour.

5.55 (m) Loose medium to coarse sand, general brown colour, abundant black grains.

6.25 (m) Very soft medium sand, light brown colour, homogeneous colour.

7.05 (m) Very soft medium sand, light brown colour, homogeneous colour.

Note: samples from 0 m to 1.04 m were not available.

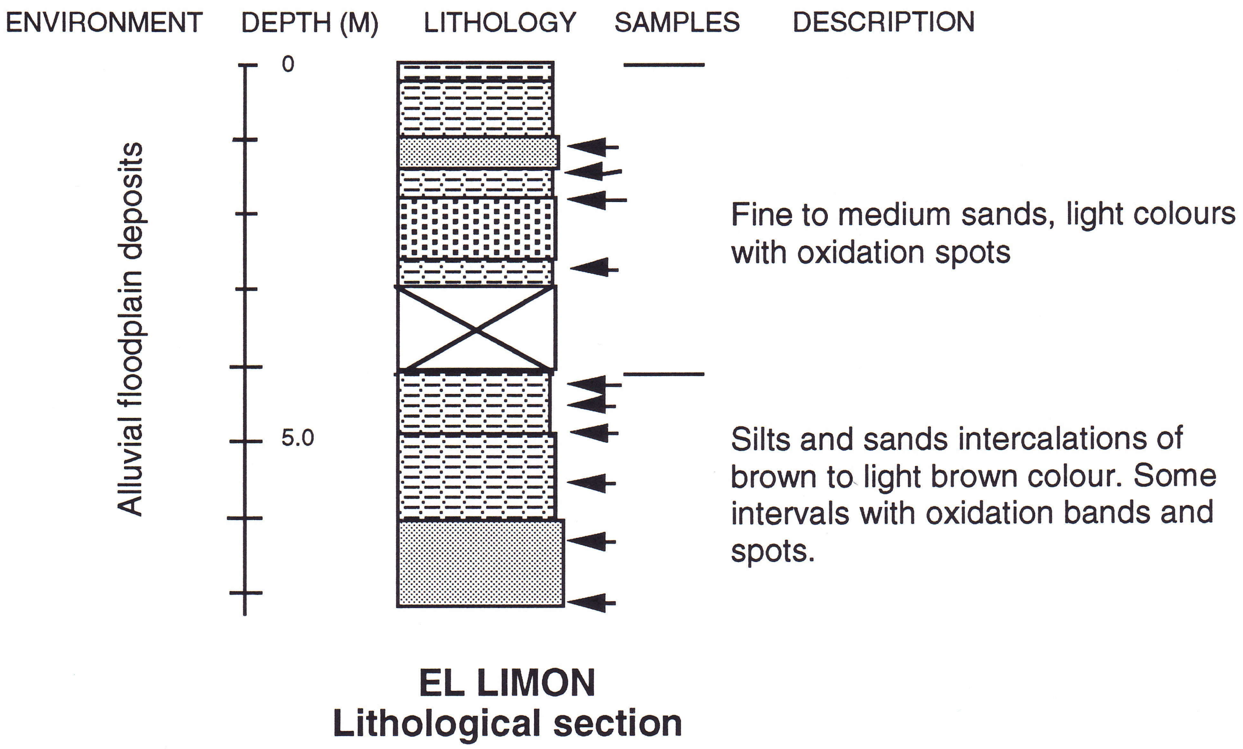


Figure 12: Lithology of the El Limon core section

#### Palynology

- Sporomorph concentration (fig. 13)

Samples are very poor to barren (70%) in this section. Only tree (1.05 m, 4.37 m and 5.55m) proved to have sporomorphs and concentration varies from five to less than 50 / 10 µl.

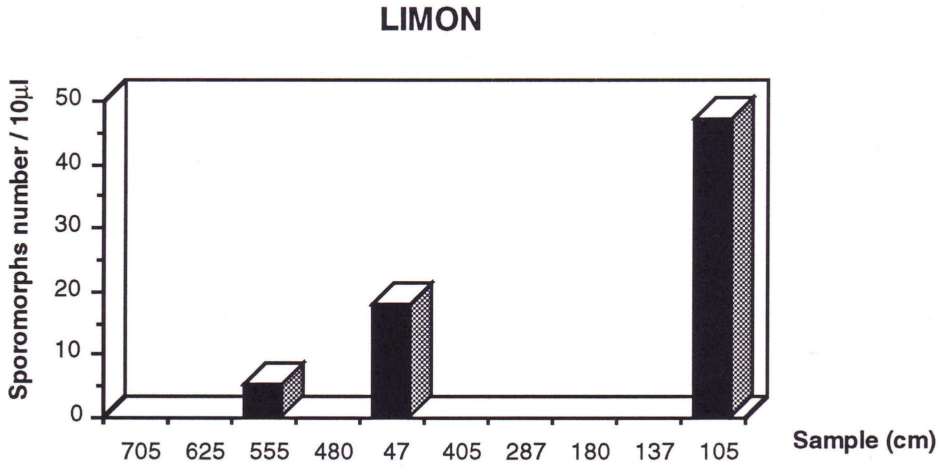


Figure 13: Sporomorph concentration Limon core section. Number of sporomorphs/10µl

- Sporomorphs diagram (fig. 13a)

The recovered assemblages were very poor, with only grass pollen and very occasionally composites pollen or poorly preserved unidentifiable types.

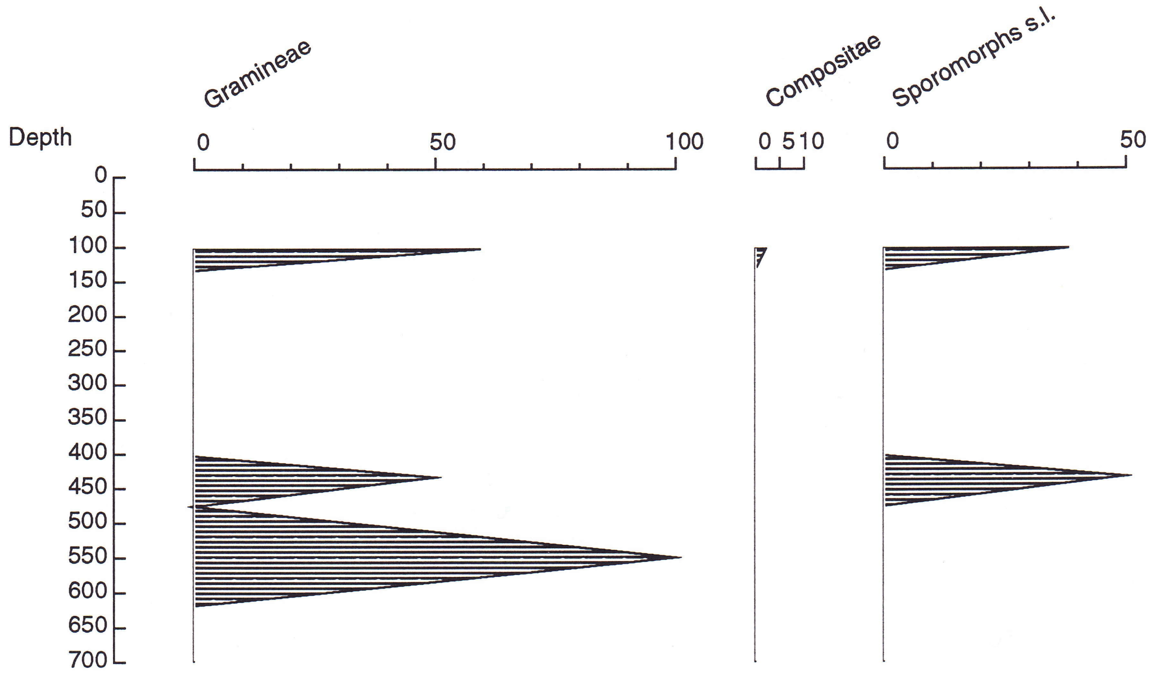


Figure 13 a: Sporomorphs diagram (expressed in %). El Limon core section

- Palynomorphs diagram (fig.13b)

Fungal remains are present in those few samples with sporomorphs.

At 1.05m the fungal remains are very abundant compared with the rest of the section.

In general samples are barren in this environment (70 % of all samples).

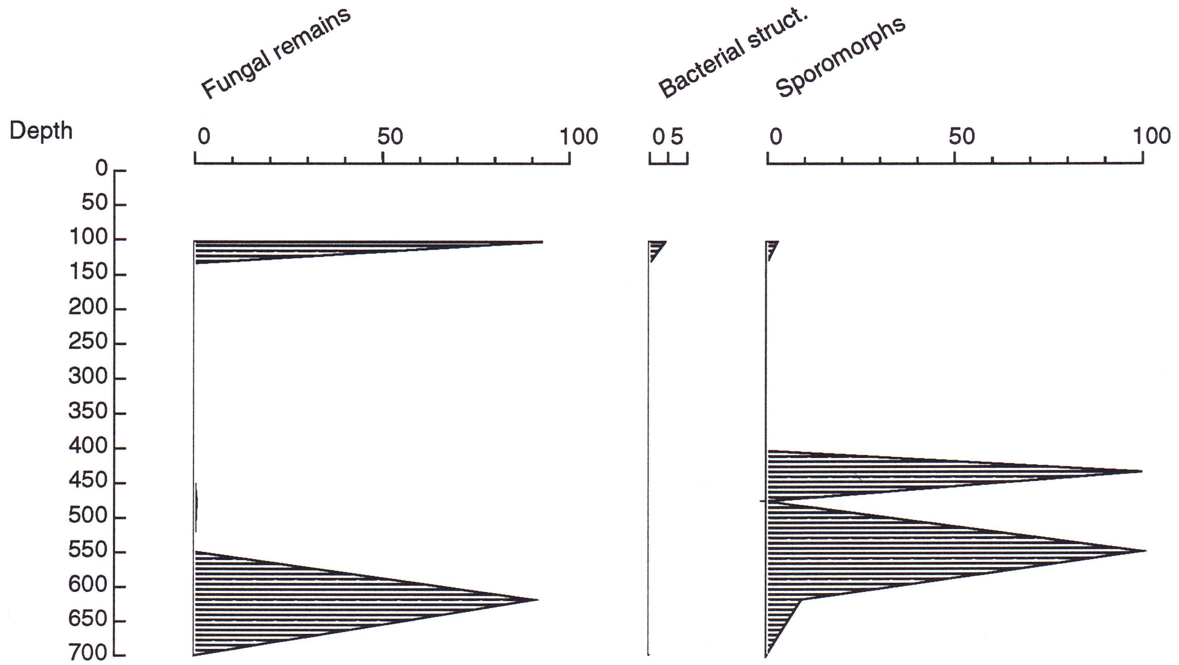


Figure 13 b: Palynomorphs diagram (expressed as %). El Limon core section

#### Palynofacies

- Organic matter concentration (fig.4c)

Organic matter recovery from this section is extremely low approaching 0 % (0.00%) concentration. Nevertheless some organic matter was recovered, but it was impossible to measure it.

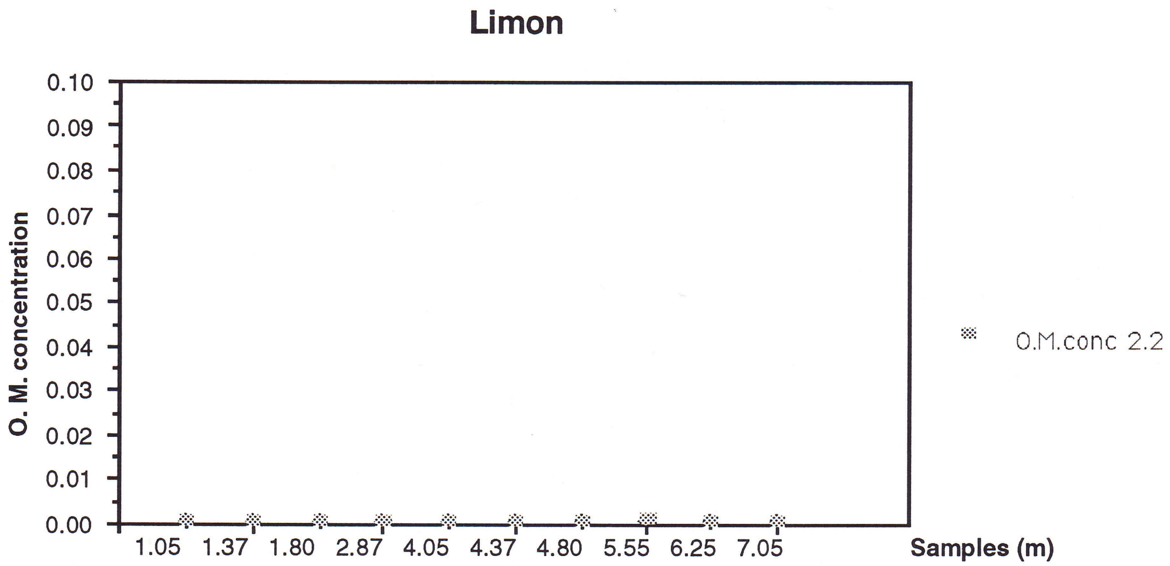


Figure 4c: Organic matter concentration Limon section. Recovered after 2.2 heavy liquid separation.

- Organic matter composition (fig. 14a &b)

Most of the organic matter from 1.37 m to 5.55 m is of "organo-mineral gel" type with comparatively minor amounts of amorphous opaque. Occasionally fungal, woody and other plant tissues are observed.

In samples 1.05 m and 6.25 m to 7.05 amorphous finely dispersed is the dominant type.

Finely dispersed organic material tends to hide the rest of the organic components (fig. 14a), but when is removed is possible to see that the organic composition of all the samples is very similar in the entire section (fig. 14). The same effect has been observed in other sections for that reason the pie diagrams in this report do not include the finely dispersed amorphous organic matter, unless it is specified.

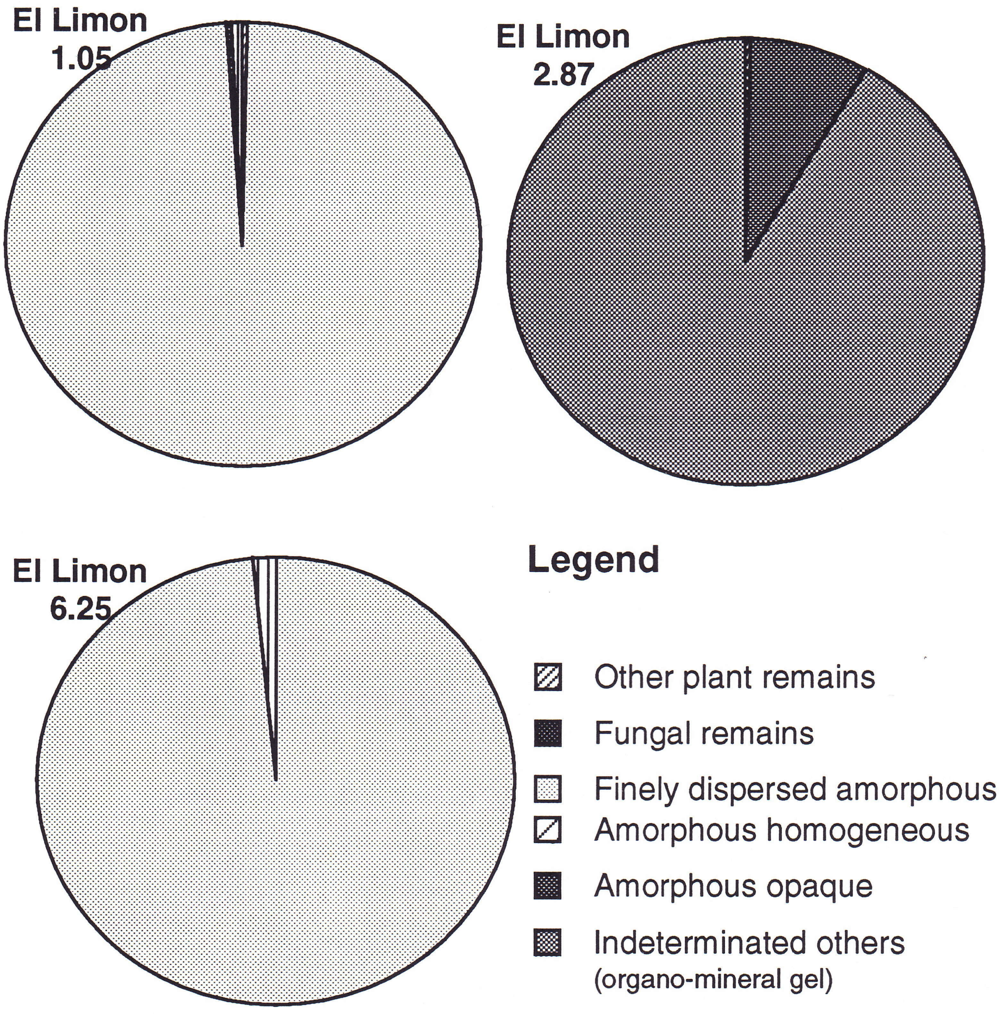


Figure 14a: Organic matter composition of selected samples from El Limon core section. Includes finely dispersed materials.

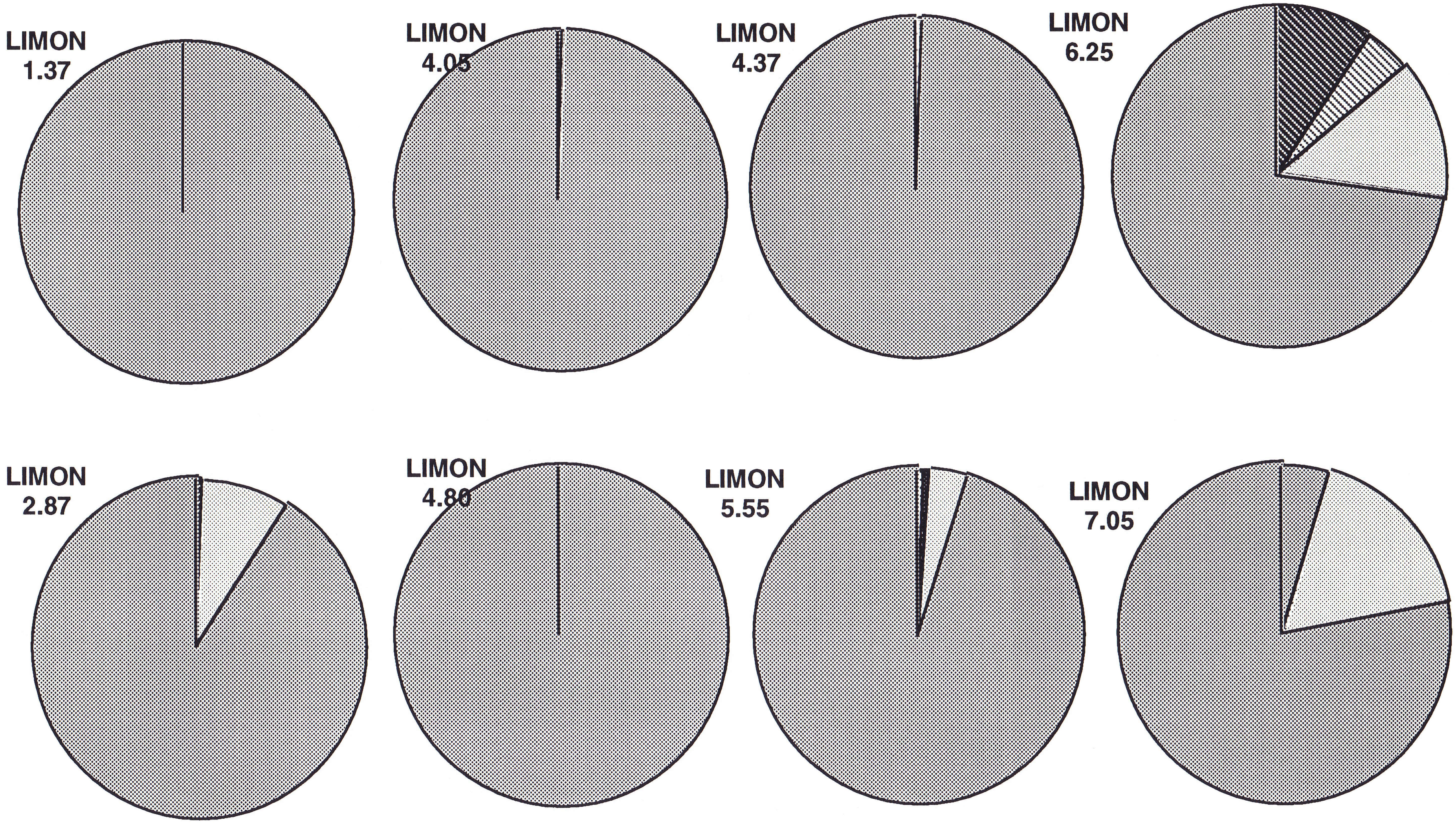


Figure 14: Organic matter composition El Limon core section

- Organic matter morphological characterization (fig. 15)

Grain size distributions range mainly from ø8 to ø3, only the interval between 5.55 m and 4.05 m have particles up to ø2.

With the exception of sample 1.05m, the sum of areas grain size distribution have a first pick at ø7 and a second pick (ø4, or ø3 or ø2) usually much more prominent than the first one.

In the grain size distribution histogram based on number of particles is clear than assemblages are dominated by particles in the range of ø8 and ø7 (clay and fine silt size range).

More than 60% of all particles fall in the equidimensional range, although very occasionally strongly elongate to filiform particles has been measured.

Sphericity varies from 0.0 to 0.8 but the main peak is usually at 0.1.

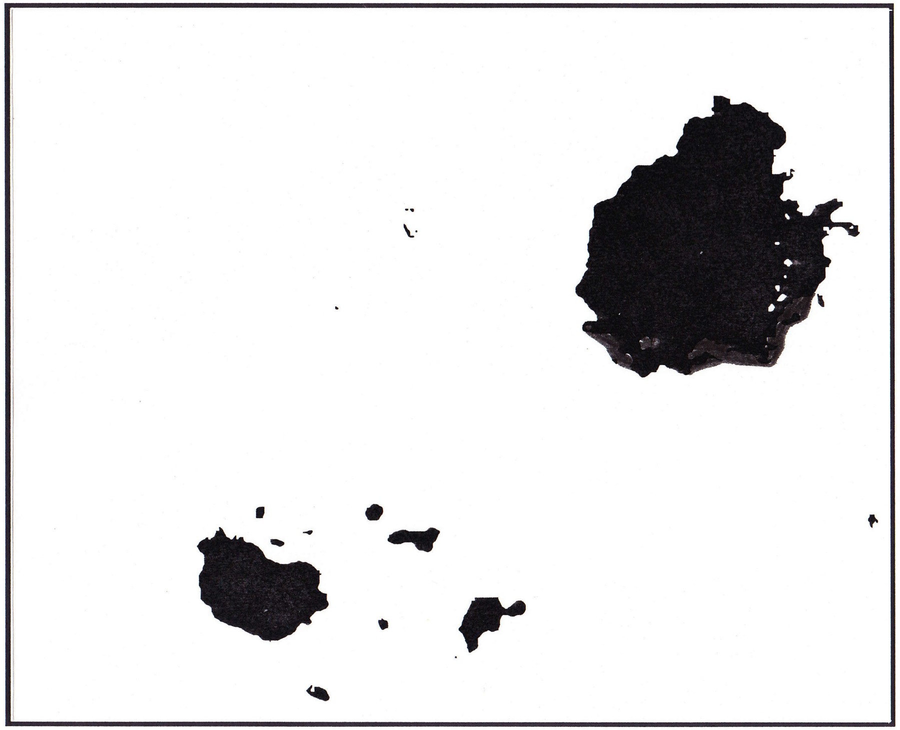

Limon 7.05

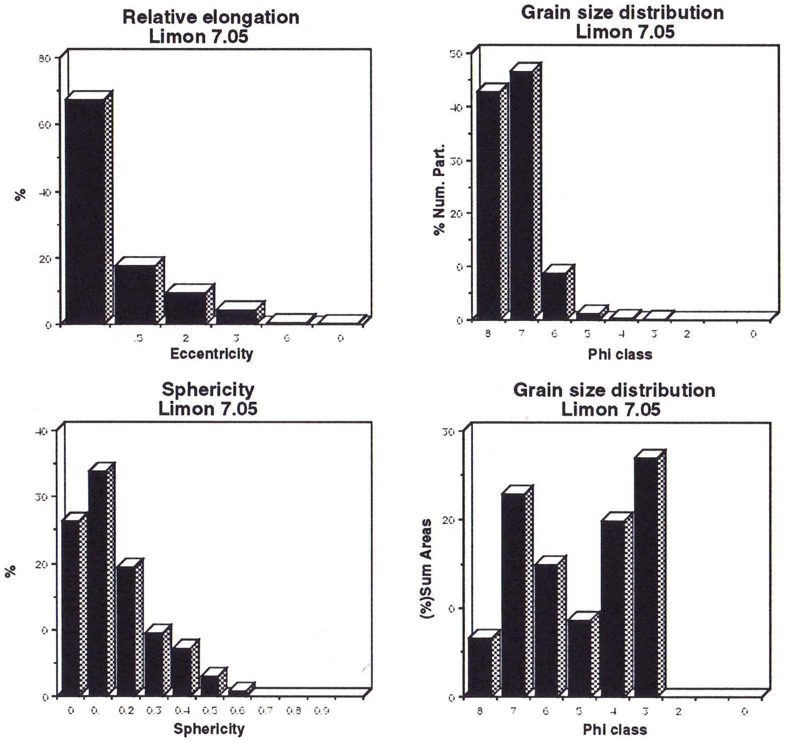


Figure 15: Organic particles morphological characterization. Selected samples from the El Limon core section

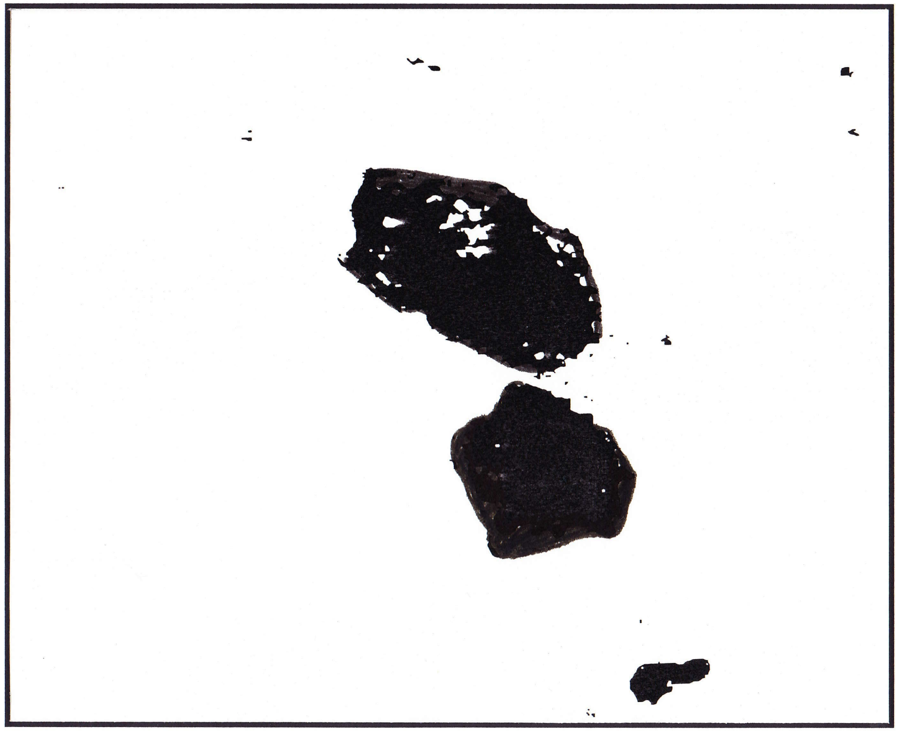

Limon 1.80

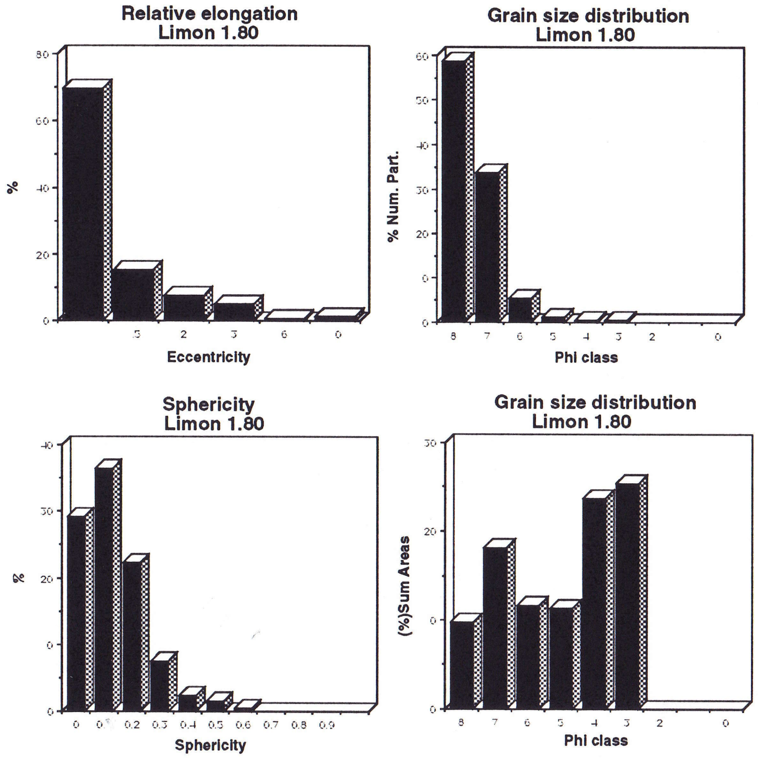


Figure 15: Organic particles morphological characterization. Selected samples from the El Limon core section

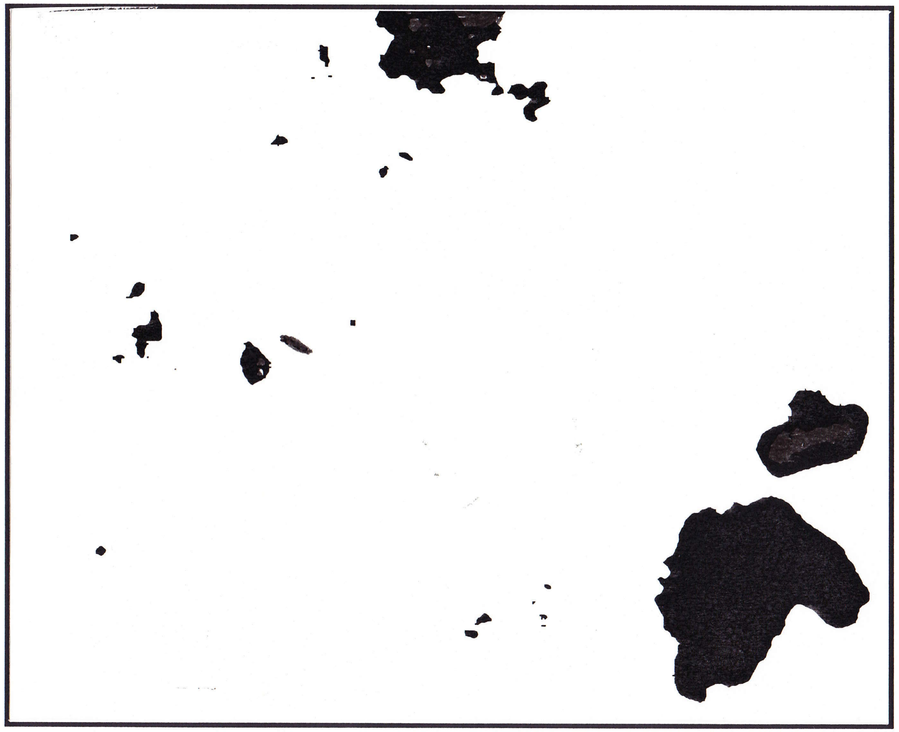

Limon 4.05

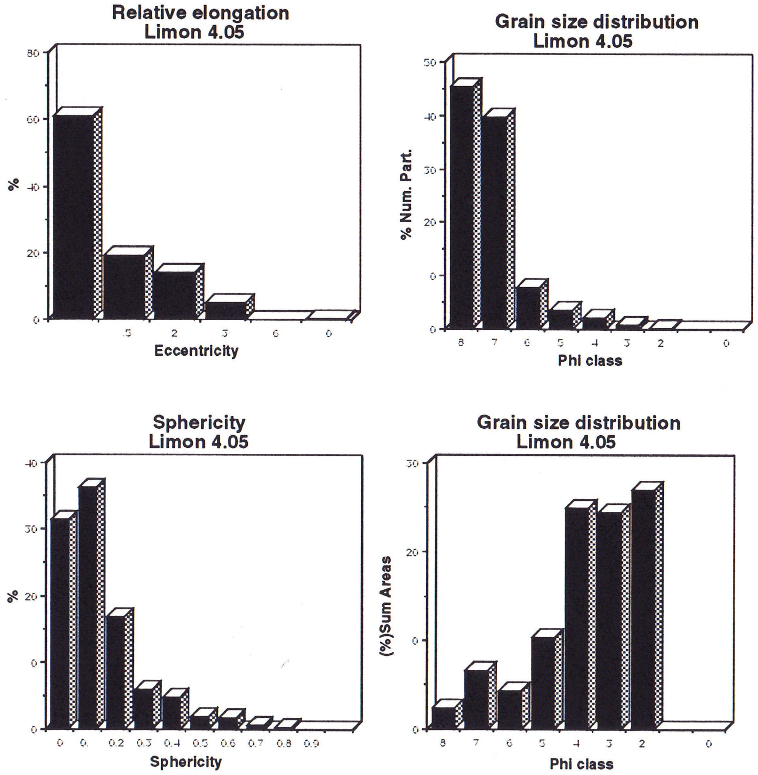


Figure 15: Organic particles morphological characterization. Selected samples from the El Limon core section

#### The Santa Marta (BL) Section

##### Location and general information

The core was taken in the "Cienaga Grande de Santa Marta", 100m in front of the Boca de Lopez mangrove, in a place with 50 cm of water depth.

##### **Lithology and sample description** (fig. 16)

A total of 25 samples were selected for this study:

0.05 Light grey to brownish clay.
0.15 Light grey to brownish clay.
0.25 Light grey to brownish clay.
0.35 Light grey to brownish sandy micaceous clay, with plant remains.
0.45 Light grey to brownish sandy micaceous clay, with plant remains.
0.55 Light grey to brownish sandy clay.
0.65 Light grey to brownish micaceous very sandy clay, with thin bivalvia shells.
0.75 Light grey to brownish micaceous very sandy clay, with bivalvia shells.
0.95 Grey to brownish silty clay
1.05 Grey to brownish silty clay
1.15 Grey to brownish silty clay
1.25 Grey to brownish silty clay
1.35 Grey to brownish silty clay, with plant remains
1.45 Grey to brownish silty clay, with plant remains
1.55 Grey to brownish silty clay
1.65 Grey to brownish silty clay

---------------------contact clay / peat ------------------------

1.70 Peat
1.80 Peat with abundant gypsum crystals
1.90 Peat with abundant gypsum crystals
2.00 Peat with abundant gypsum crystals
2.10 Peat
2.20 Peat
2.30 Grey to brownish clay and peat
2.40 Dark grey to brownish clay with organic remains
2.50 Dark grey to brownish clay with organic remains

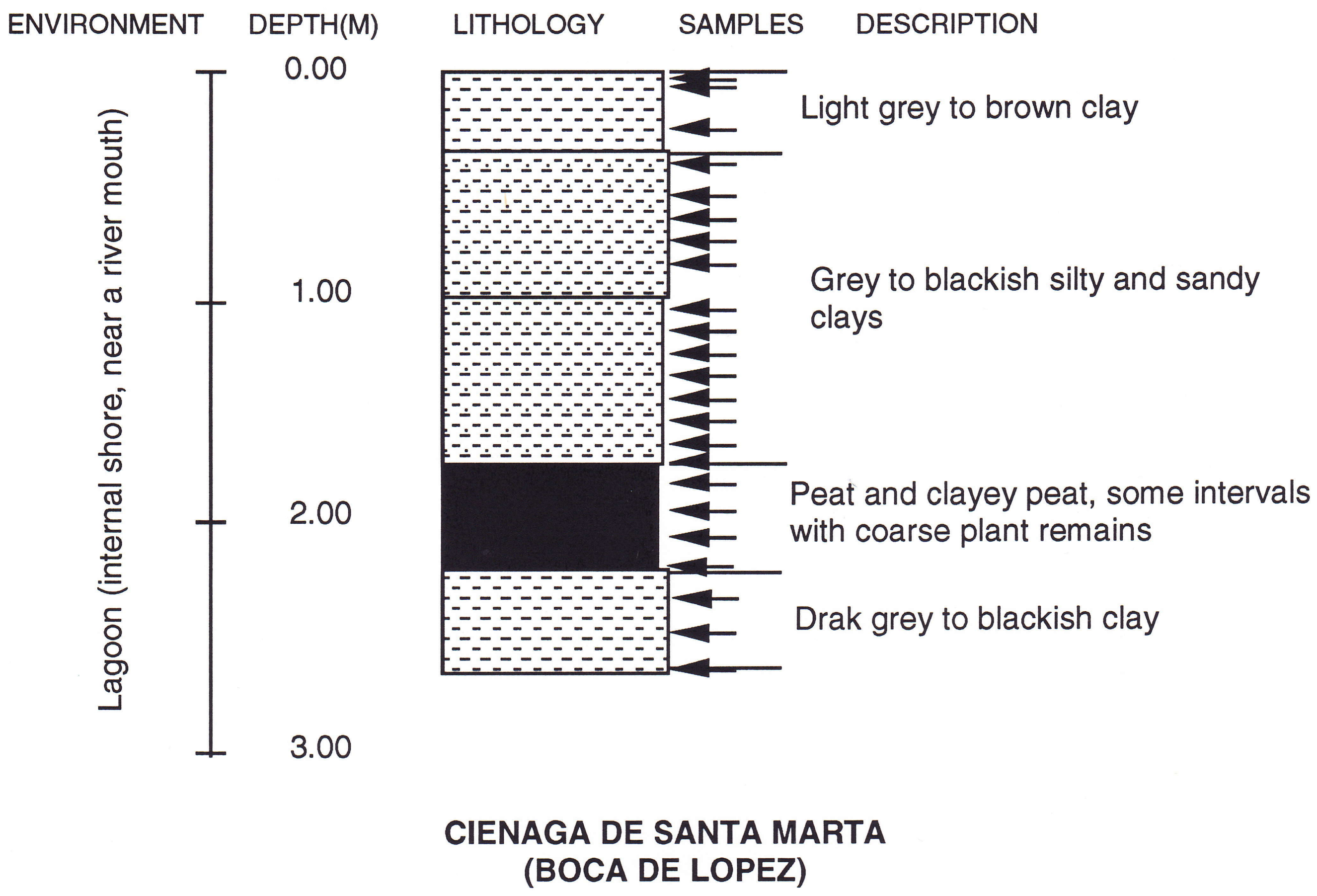


Figure 16: Lithology of the Santa Marta core section

#### Palynology

Sporomorph concentration varies between 100 and more than 700 specimens per slide. This environment proved to be one of the richer in sporomorphs. Only one sample from a total of twenty five was barren (fig. 17a).

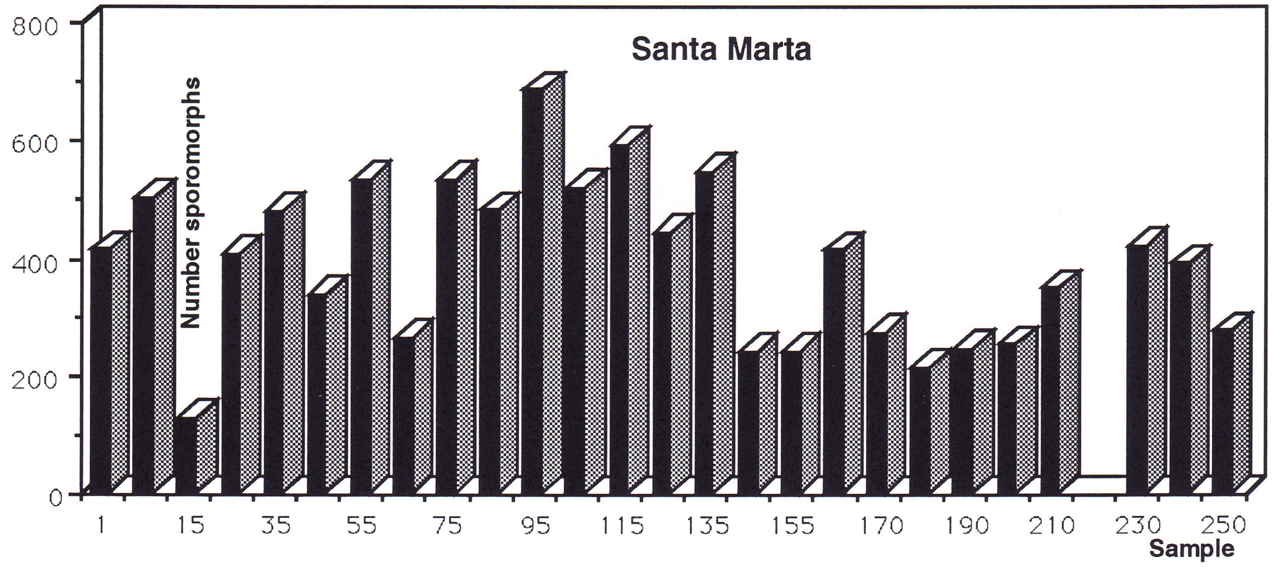


Figure 17a: Sporomorph concentration. Number of grains/slide.

- Pollen and spores & ecological groups diagram (fig.17 and 17b)

The assemblage from the Santa Marta Lagoon is the richest in species and specimens.

Sporomorphs diagram shows two main zones. A lower zone from 2.50 to 1.70 m dominated by mangrove pollen and an upper zone from 1.65 m to 0.05 m. dominated by pollen grains of composites, grasses and water plants.

In the lower zone (2.50m to 1.70 m) two intervals can be recognized: from 2.50 to 2.30 m the lower interval with an assemblage dominated by pollen grains of mangrove and water plants, together with psilate monolete spores. A barren sample at 2.20 m subdivides both intervals. An upper interval with an assemblage dominated by mangrove pollen.

The upper zone assemblage (1.65 m to 0.05 m) prevail the pollen of water and floating plants, present are considerable amounts of the composite, grass, mangrove pollen and psilate monolete spores. A series of types are restricted to this zone as Jussiaea, Proteaceae, *Mauritia* sp. and *Acrostichum* sp.

The group diagram show the same tendency that the sporomorphs diagram. The increase of pollen grains of water plants, composites and grasses in the clay section show a strong contrast with the mangrove sporomorphs association that characterizes the peat interval.

The peat interval has been identified at about the same depth (2 m) in other areas of the lagoon (Wiedemann,1973).

Cohen and Wiedermann (1973) concluded that the peat from the northern areas of the lagoon was a mangrove peat (*Rhizophora mangle*) while the peat from the south was originated in brackish to fresh water (*Acrostichum* sp. and Cyperaceae)

- Palynomorphs diagram (fig.17c)

The same zones can be recognized in the palynomorphs diagram.

The lower zone dominated by fungal remains and the upper zone dominated by sporomorphs.

*Botryococcus* sp. is restricted to the upper zone.

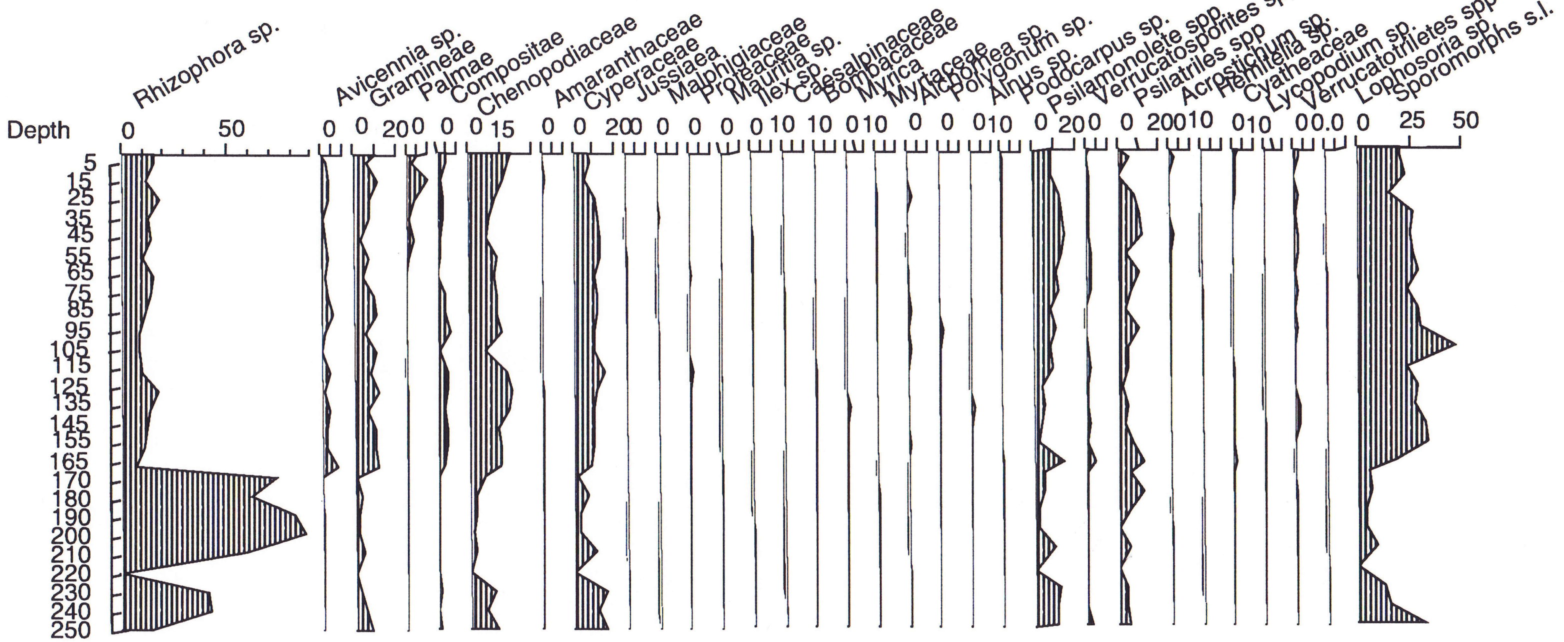


Figure 17: Santa Marta core section. Sporomorphs diagram, bars are % of total sporomorphs countings. Diagrams from this section are based on Mr. G. Noldus countings.

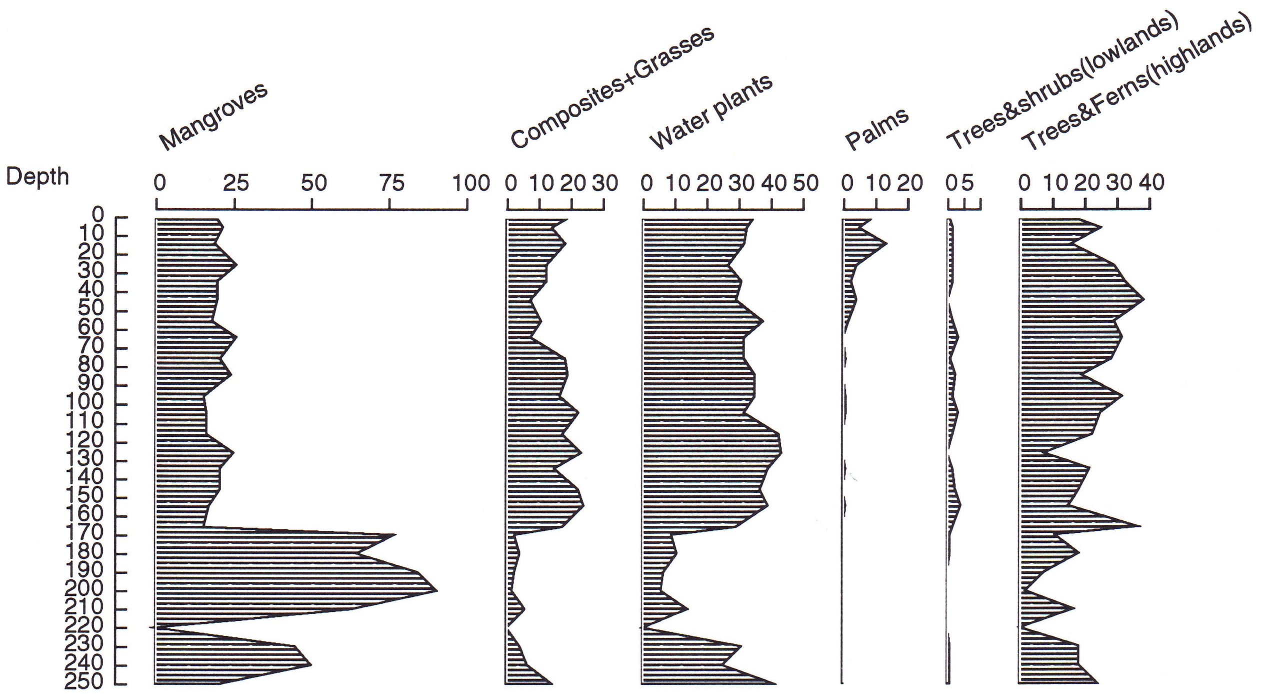


Figure 17b: Group diagram. Santa Marta (BL) core section

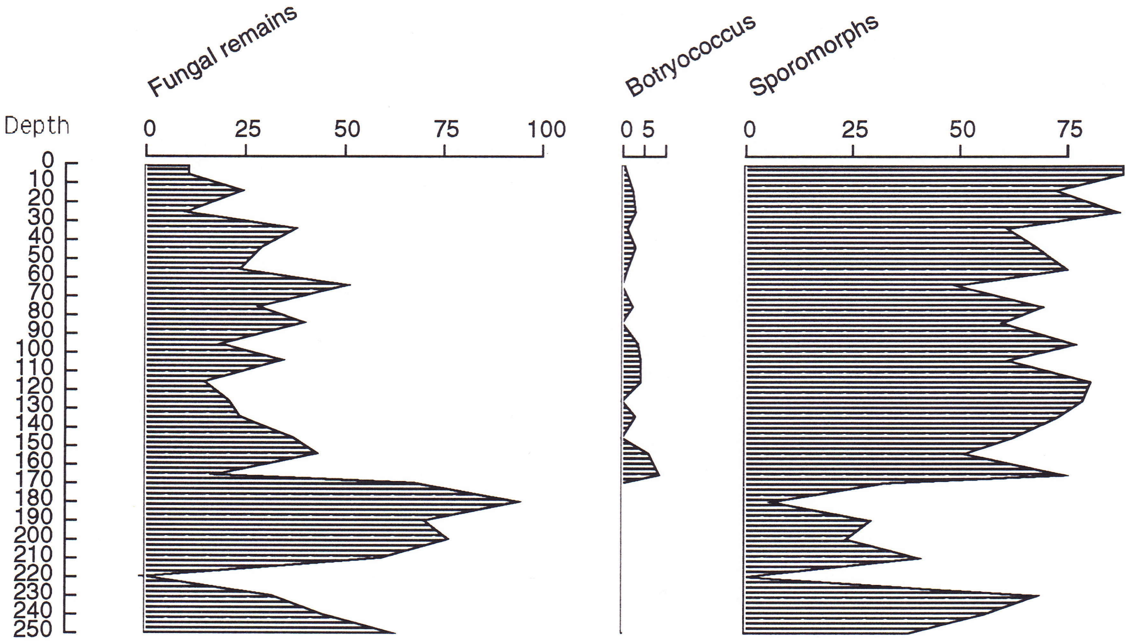


Figure 17c: Palynomorphs diagram. Cienaga de Santa Marta (BL) core section

#### Palynofacies

- Organic matter concentration (fig.4d)

Samples from the lagoon are richer in organic matter than the rest of the studied section. Concentration varies from 0.13% to values close to 1%. Excluding the peats.

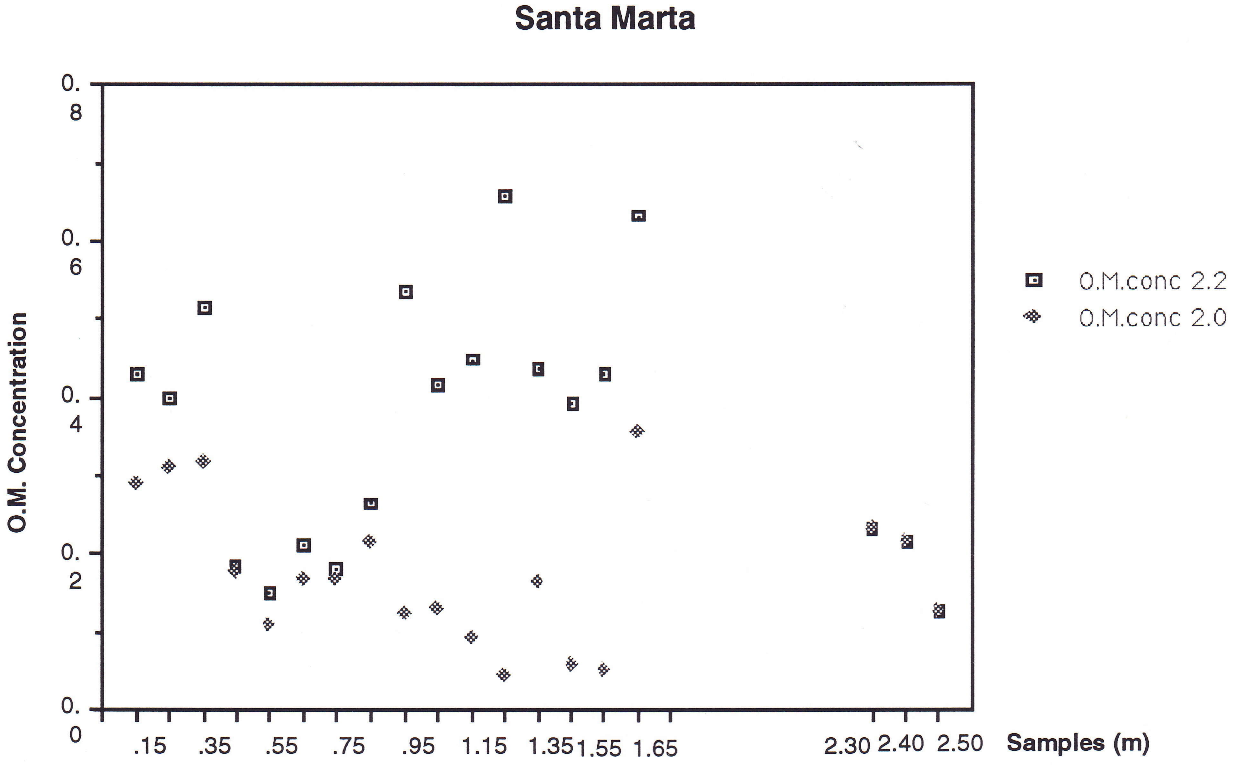


Figure 4d: Organic matter concentration. Heavy liquid separation 2.2 and 2.0

- Organic matter composition (fig. 18)

There are four apparent changes in the organic composition of the assemblages. Changes do not always coincide with changes in the lithology as might be expected.

From bottom to top, the first assemblage (2.50m to 2.30 m) is characterized by relatively more abundant finely dispersed organic matter, amorphous opaque, fungal and palynomorphs.

Sample 2.20 m that represents the starting of the peat section is composed of epidermal/cuticular, woody, "other" plant remains and amorphous homogeneous.

From 2.10 m to 1.80 m. finely dispersed organic matter is very abundant together with fungal remains. Epidermal/cuticular, other plant remains and palynomorphs are also important.

Samples 1.70 m (peat) and 1.65 m.(clay) although having a different lithology, the organic composition is very similar. Mainly amorphous material (homogeneous and heterogeneous), a decrease in the concentration of the finely dispersed material together with an strong diminution of the fungal remains, makes these samples different from the rest of the section.

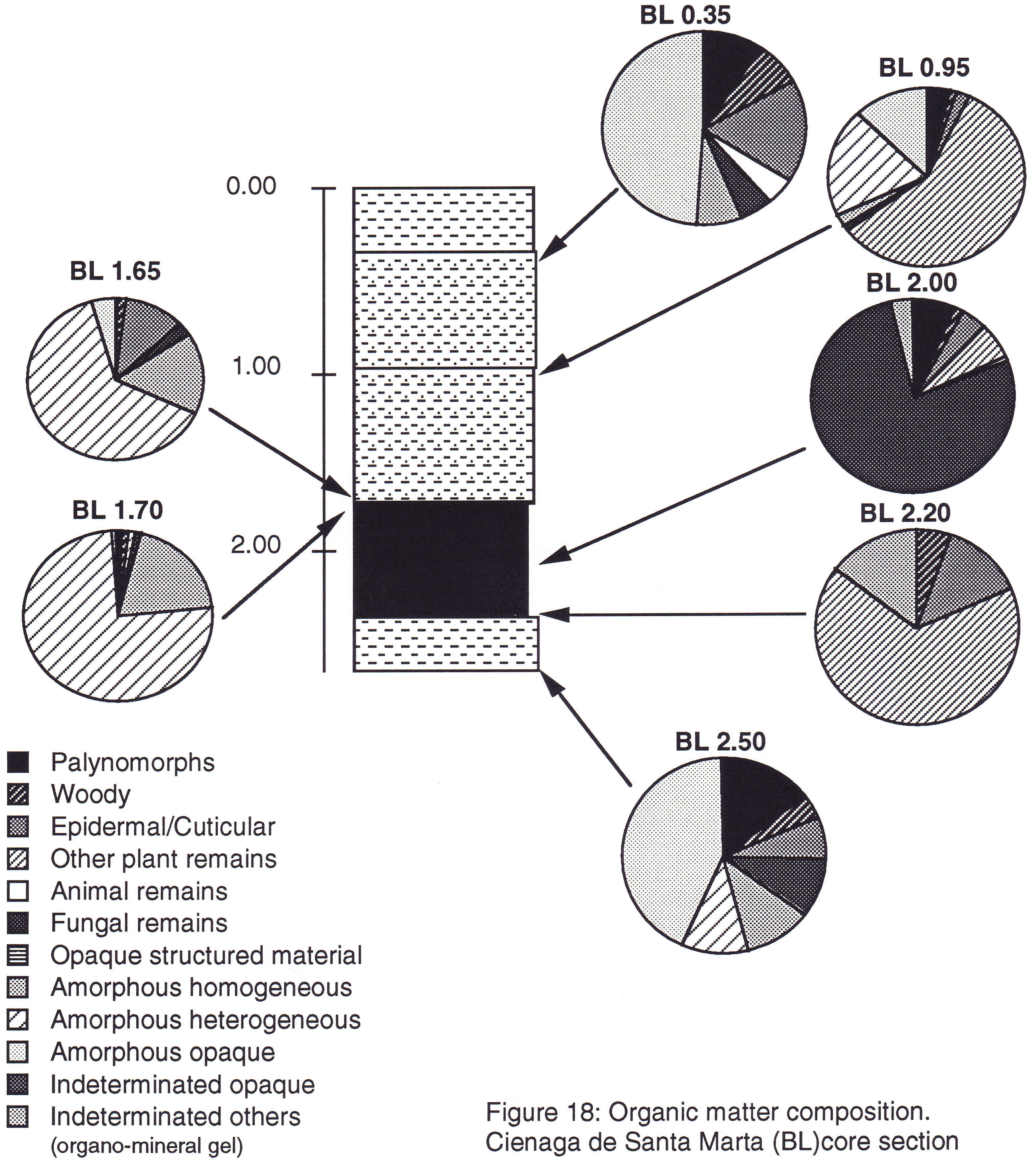


Figure 18: Organic matter composition.
Cienaga de Santa Marta (BL)core section

Samples from 1.55 m to 0.75 m have in general similar composition with the exception of variations in concentration of finely dispersed organic matter that is relatively low compared with the rest section.

Samples from 0.65 m to 0.05 m are characterized by high concentrations of finely dispersed organic matter, that dominates the whole residue. Big fragments of different plant remains, or of amorphous material are 'floating' in a fined grained "matrix".

- Organic matter morphological characterization (fig. 19)

Grain size sum of areas distributions show a general trend to decrease on the average grain size from base to top. Tree main "breaks" can be pointed out:

- Interval 2.50 m to 1.70 m: presents the coarsest grains, up to ø1 registered. The clay samples (2.50 m to 2.30 m) are characterized by bimodal sum of areas grain size distributions and dominance of the 0.1 class in the sphericity diagram. The peat samples show very strong peaks in class ø1 (up to 70%) and some times in class ø2 (about 35 to 40 %). The 0.1 class is also dominant in the sphericity diagram. About 50% of the shapes are equidimensional.

- Interval 1.65 m to 0.95 m: show unimodal sum of areas grain size distributions with the main peak usually between classes ø5 and ø3. Between 40 and 50% of the eccentricity values are concentrated in the range of quasiequidimensional to elongate tabloids, with not less than 10% of the particles belonging to one of these classes, some samples show higher values on the elongated tabloids shapes.

- Interval from 0.75 m to 0.05 m: most samples show unimodal distributions on the sum of areas histograms with maximum around ø6 and ø5, although some samples (0.35 m 0.55 m and 0.65 m) show a maximum in ø3. Most samples have a maximum in the sphericity histogram at 0.1 class. More than 50% of the particles are in the equidimensional range of value.

All these changes are closely related with changes in the lithology, mainly in the relative amount of silt and / or sand in the clays and the general colour of the sediments.

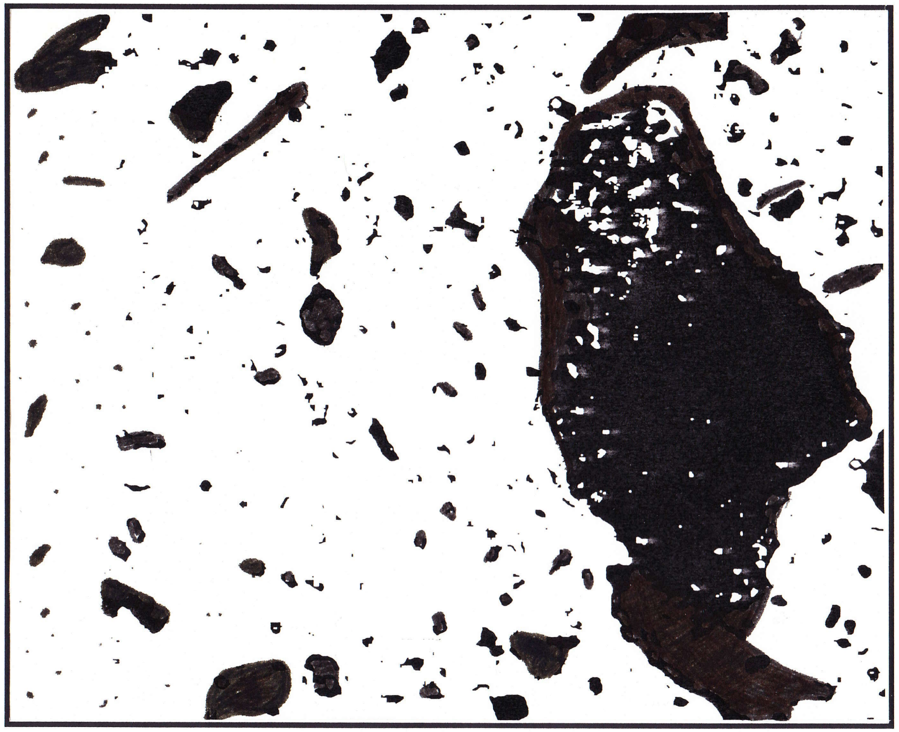

Santa Marta Lagoon (BL 2.40)

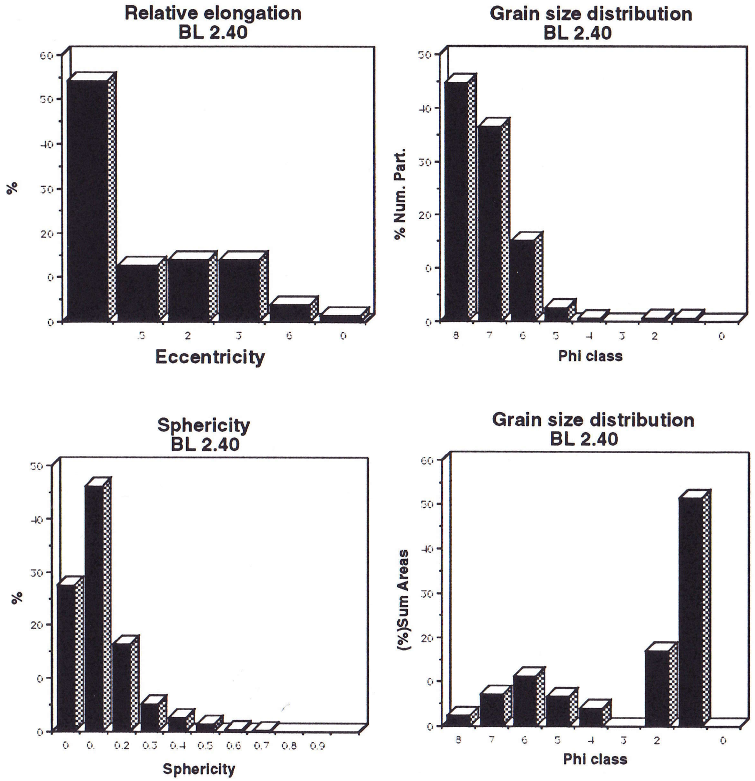


Figure 19: Organic particles morphological characterization. Selected samples from the Santa Marta Lagoon (BL) section

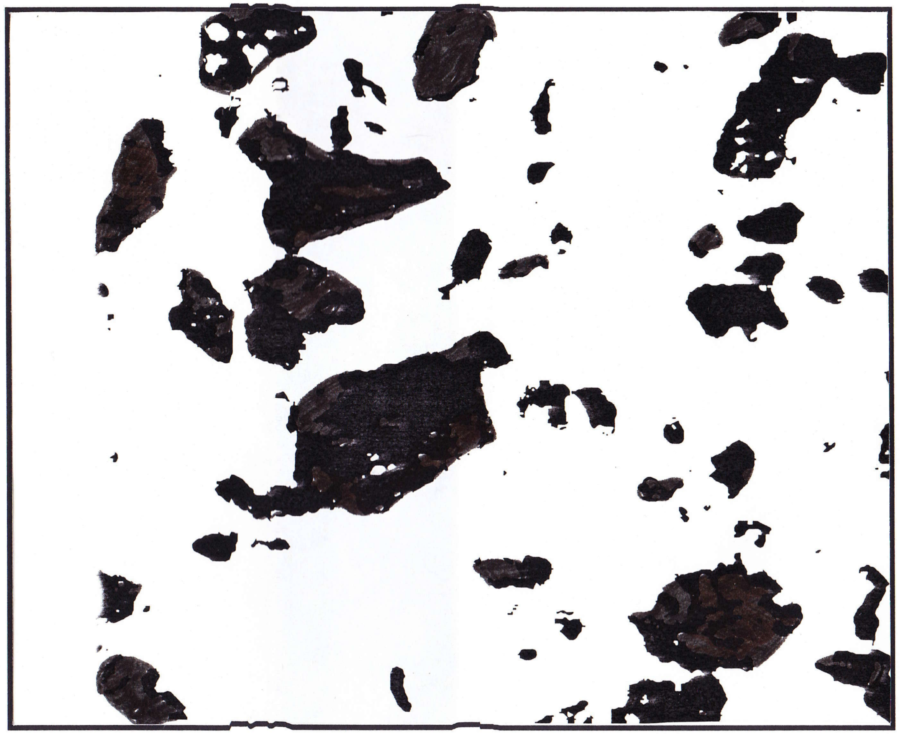

Santa Marta Lagoon (BL 1.25)

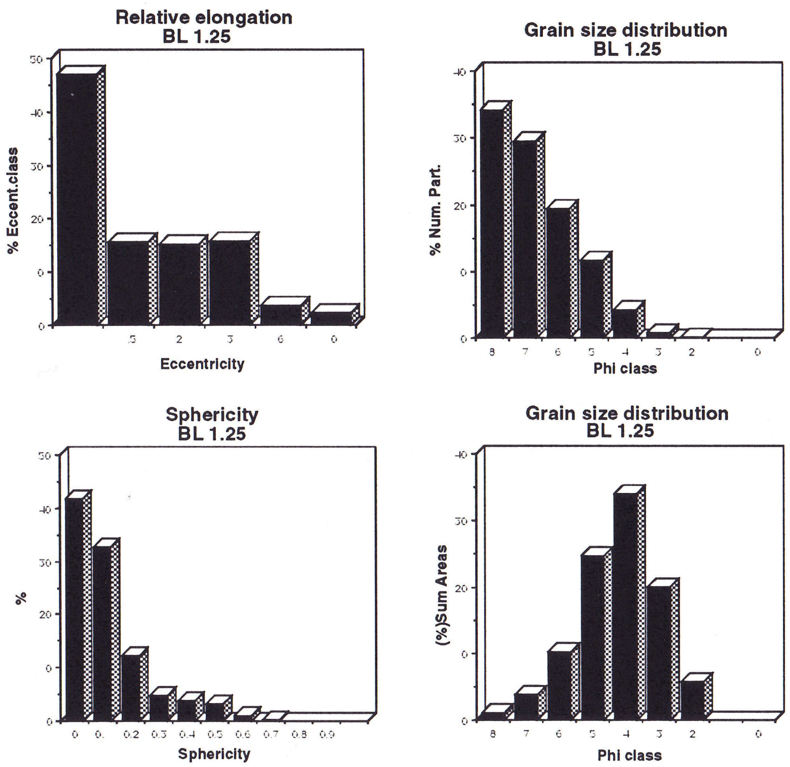


Figure 19: Organic particles morphological characterization. Selected samples from the Santa Marta Lagoon (BL) section

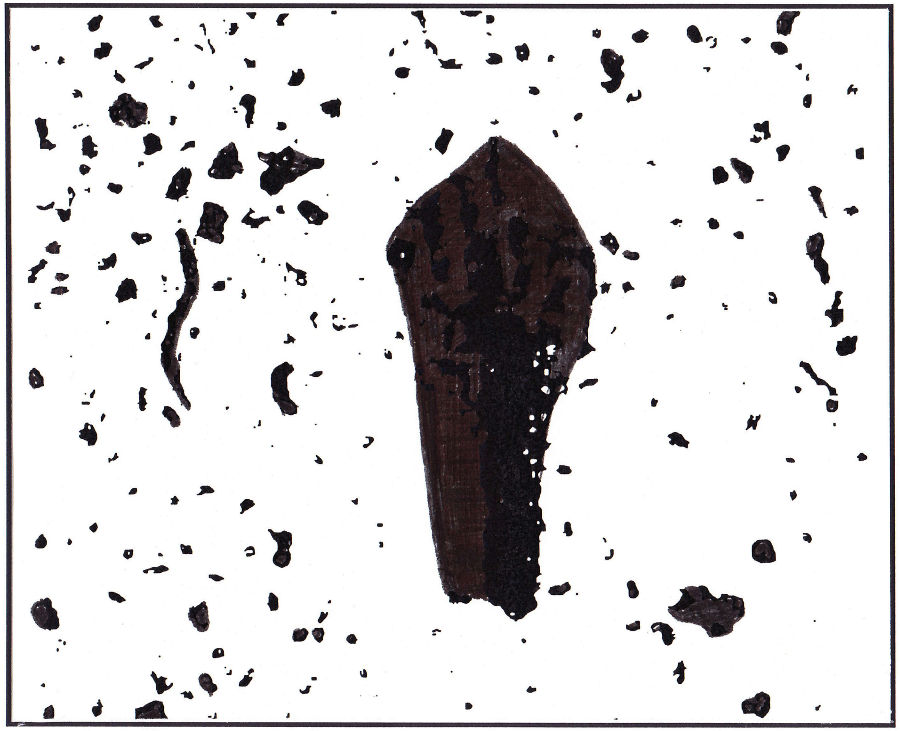

Santa Marta Lagoon (BL 0.45)

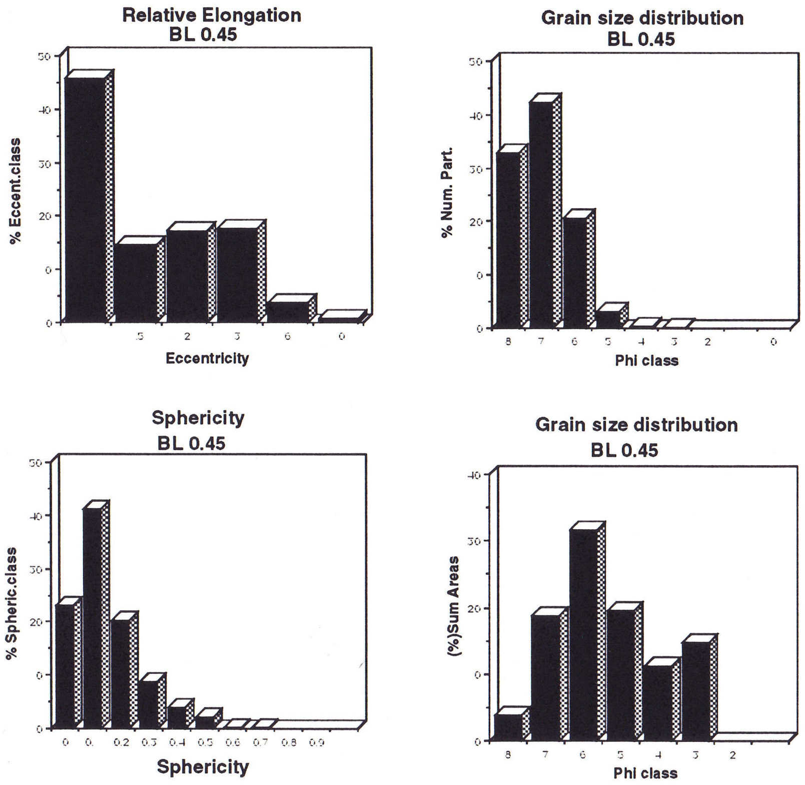


Figure 19: Organic particles morphological characterization. Selected samples from the Santa Marta Lagoon (BL) section

#### The surface samples

##### Location and general information

A set of five samples were taken from the bottom surface of five different swamps from the Lower Magdalena Basin. The sampled swamps were: Cienaga El Medio, Cienaga Pimiento, Cienaga Coyongal, Los Limones and Punta de Blanco. All situated between 8º 40'N - 9º 10' N and 74º95' W- 74º25'W.

##### Lithological description

CM (Cienaga El Medio): Silty clay, medium gray colour, with oxidation spots and roots.

CP (Cienaga Pimiento): Silty clay, medium gray colour, with oxidation spots.

CY (Cienaga Coyongal): Silt with clay, light brown colour, with plant remains.

Los Limones: Silty clay, medium gray colour, with oxidation spots and root remains

PB (Punta de Blanco): Clayey silt, light brown colour, with root remains.

#### Palynology

Sporomorph concentration in the swamp surface samples varies from low concentrations eg. less than 50 grains/10µl to more than 250 grains /10µl (fig. 20a).

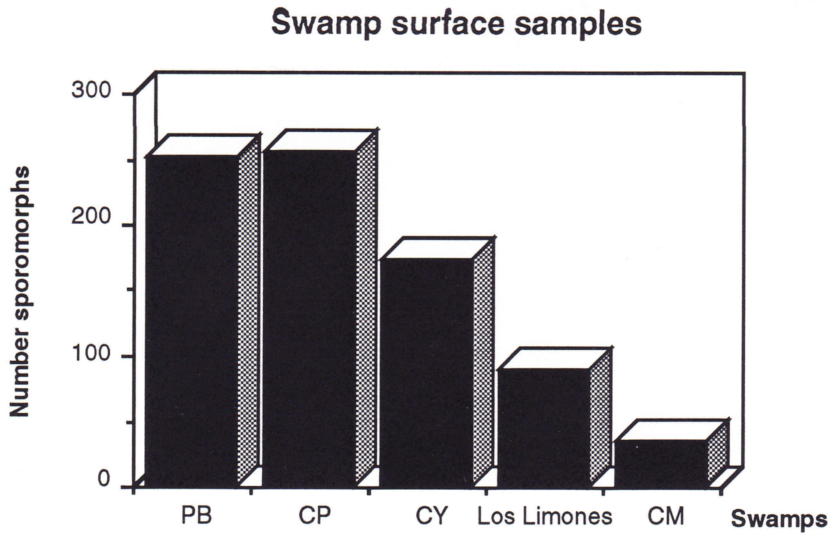


Figure 20a: Sporomorph concentration. Number of sporomorphs / 10 µl.
PB: Punta de Blanco; CP:Cienaga Pimiento; CY: Cienaga Coyongal and CM: Cienaga El Medio.

- Pollen and spores diagram (fig.20)

The Cienaga Coyongal assemblage is characterized by abundance of Gramineae, Compositae, Cyperaceae and psilate monolete spores.

The Cienaga Pimiento: assemblage is characterized by the abundance of Gramineae, Compositae, Jussiaea pollen and psilate monolete spores. Cyperaceae, *Althernantera* and *Croton* pollen are also present.

Los Limones: assemblage is characterized by the abundance of Gramineae, Compositae, *Croton* . pollen and psilate monolete spores. *Althernantera* pollen is also present.

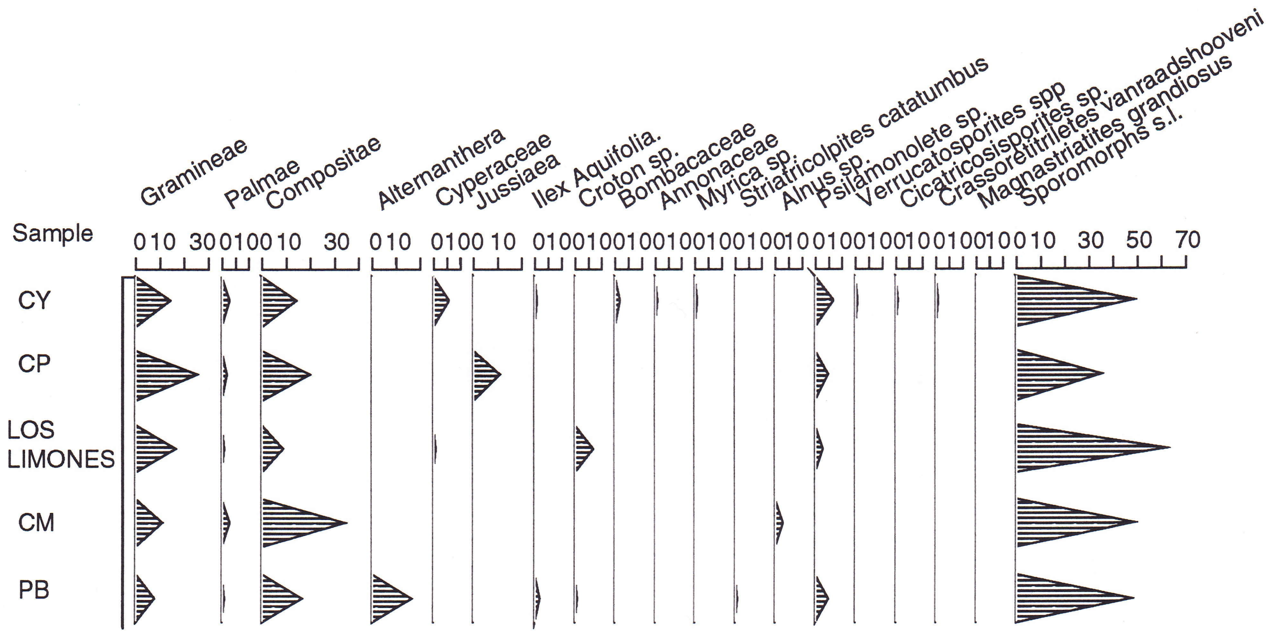


Figure 20: Sporomorphs combined diagram. Swamp surface samples

Cienaga El Medio: assemblage is characterized by the abundance of pollen grains of Gramineae and Compositae.

Punta de Blanco: assemblage is characterized by the abundance of pollen grains of Compositae and *Althernantera* , also important Gramineae pollen and psilate monolete spores. Cyperaceae and *Croton* pollen are also present.

In general the palynomorphs assemblage is dominated by pollen grains of grass and composite and psilate monolete spores. Also present in significative amounts are occasionally *Althernatera*, Cyperacea, *Croton*, and *Ludvigia* (Jussiaea).

- Palynomorphs diagram (fig.20b)

In the Cienagas Coyongal, Pimiento y El Medio fungal remains are the most abundant type of palynomorph.

In Los Limones and Punta Blanco, fungal remains account for between 2.7 and 5% while sporomorphs do not exceed from 0.3% of the total organic residue, after excluding the finely disperse amorphous matter.

Always present in the assemblages are insect remains and fresh water algae (also *Botryococcus*). Occasionally conodonts can be founded in the assemblages.

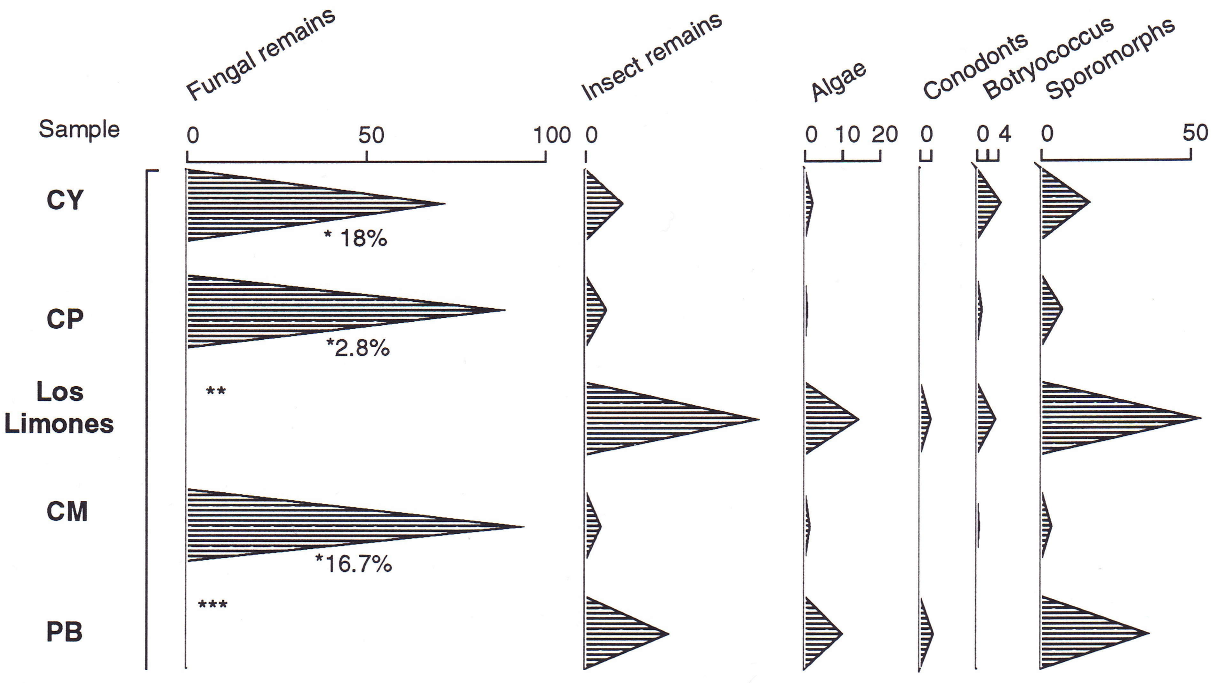


* Percentage of fungal remains as related with the total organic matter excluding amorphous finely dispersed materials)

** Not Quantified (~ 5% of the total organic matter excluding amorphous finely dispersed)

*** Not Quantified (2.7% of the total organic matter excluding amorphous finely disperse

Figure 20b: Palynomorphs combined diagram. Swamp surface samples

#### Palynofacies

- Organic matter concentration (fig.4e)

The swamp surface assemblages are characterized by comparatively high concentrations of organic matter between 0.1 % and 0.3% (after 2.2 heavy liquid separation). The measured concentration drops sharply after 2.0 density separation, probably related with losses of the very fine material.

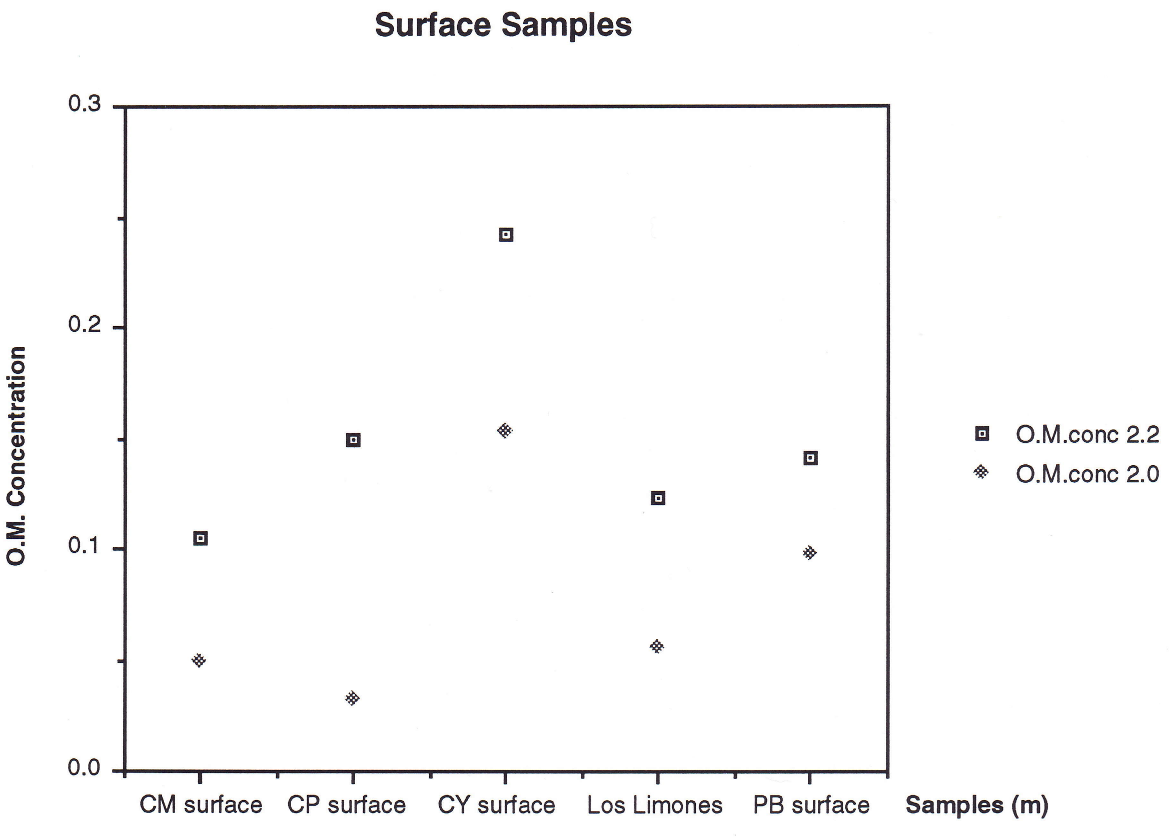


Figure 4e: Organic matter concentration. After 2.2 and 2.0 heavy liquid separation

- Organic matter composition (fig. 21)

The composition of the organic matter show abundance of light coloured plant tissues, but also epidermal/cuticular materials, fungal remains, opaque amorphous, and animal remains, together with very abundant amorphous material finely dispersed.

Figure 21 include the composition of Ayapel 0.25 as comparison between shallow lake close to the surface and the swamp surface.

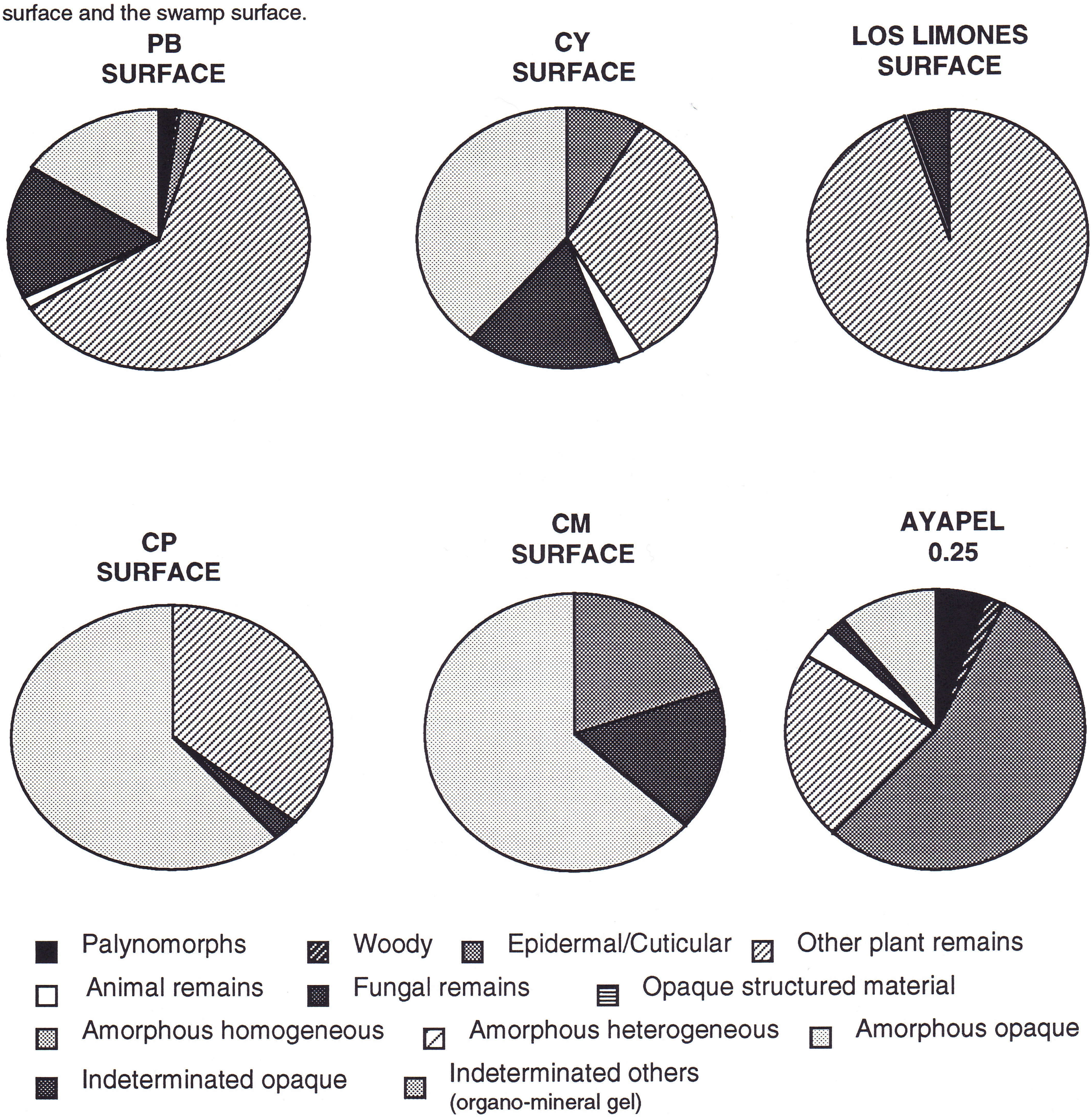


Figure 21: Organic matter composition. Swamp surface samples.

- Organic matter morphological characterization (fig. 22)

In general these assemblages are characterized by a number of particles grain size distribution with materials in the clay size range below 47%, while according to the sum of areas histogram covering only 2 to 3% of the total particle area.  Materials over ø2 (from fine sand upwards) cover more than 60% of the total particle area.

The CM sample is numerically dominated by particles in the silt size range (ø7), but all the particles comprised between ø8 and ø3 (clay to very fine sand) cover less than 50% of the particle area, an average of less than 7% per category, whereas particles with sizes bigger than ø2 cover more than 50% of the particle area.

The relative elongation histograms from swamp surface samples show that although 45 to 60% of the particles fall in the equidimensional category usually not less than 20% fall in the clearly elongate forms.

Most of the sphericity values fall in the 0.0 to 0.2 classes and they can reach up to class 0.8. The exception is the Punta de Blanco sample in which the 0.1 class exceeds even the most abundant classes in more than 20%.

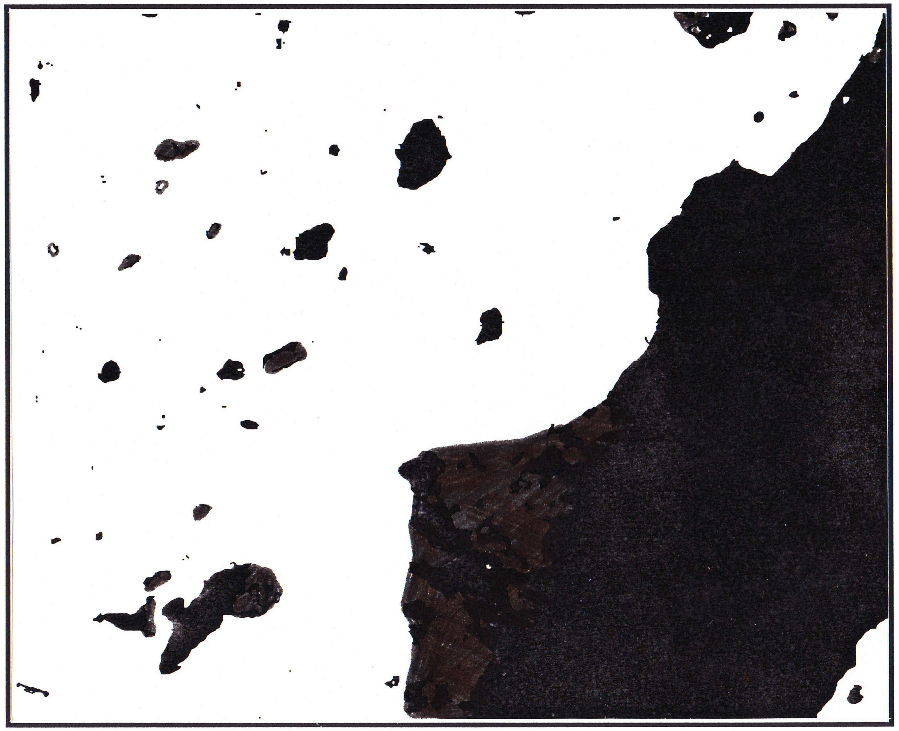

Cienaga Pimiento

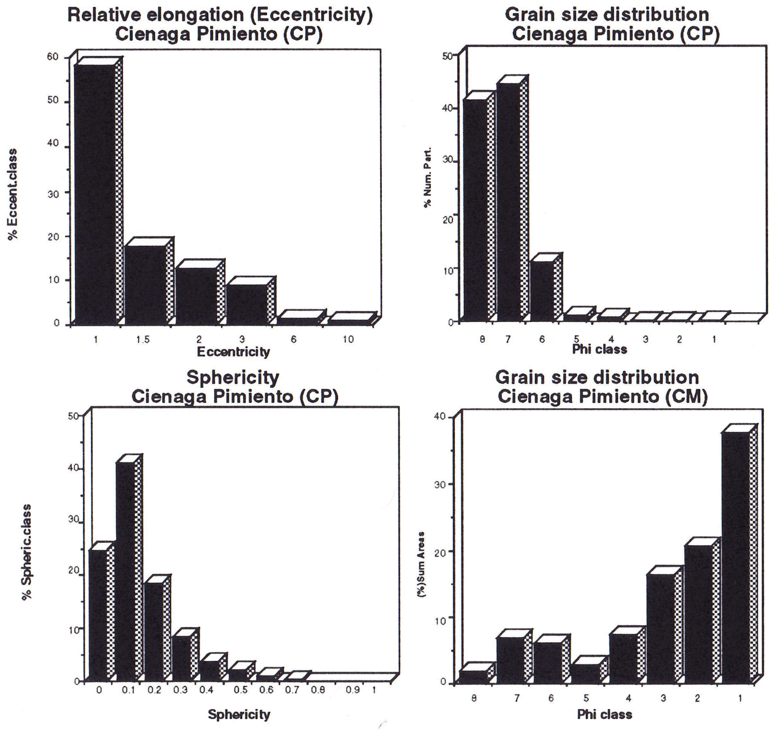


Figure 22: Organic particles morphological characterization. "Cienaga Pimiento" swamp surface sample

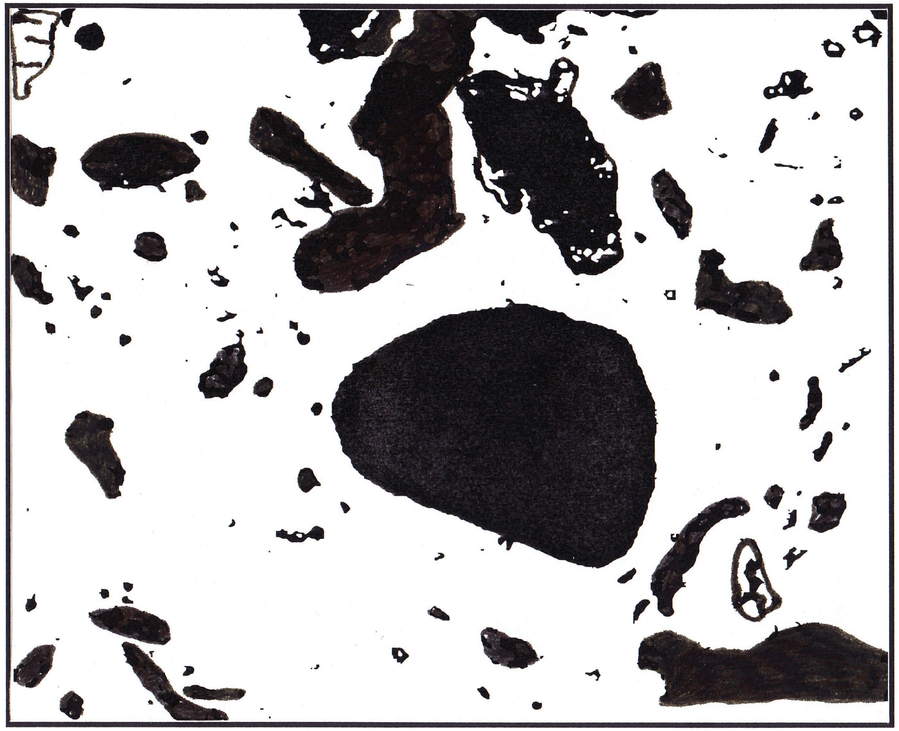

Cienaga Coyongal

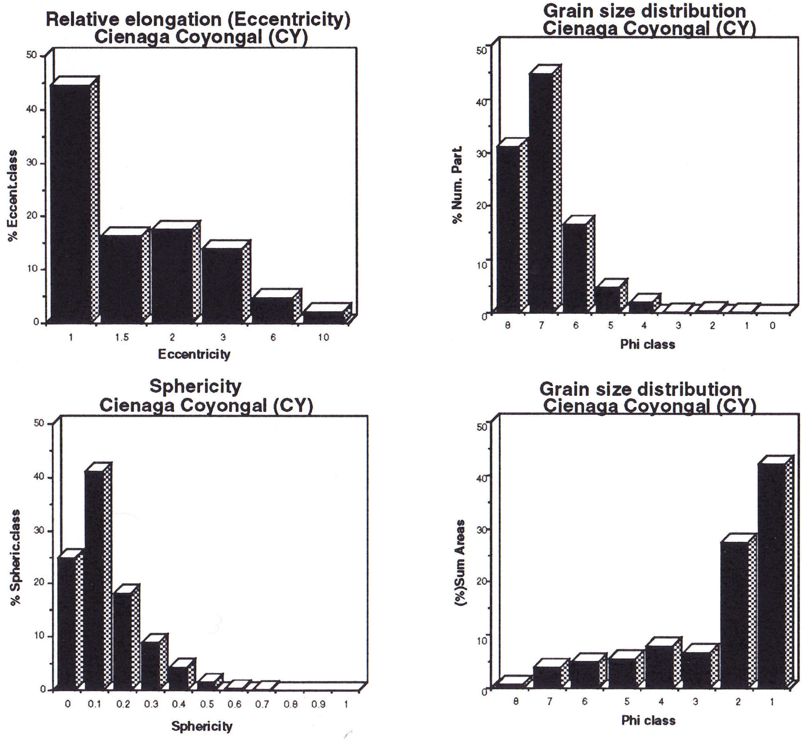


Figure 22: Organic particles morphological characterization. "Cienaga Coyongal" swamp surface sample

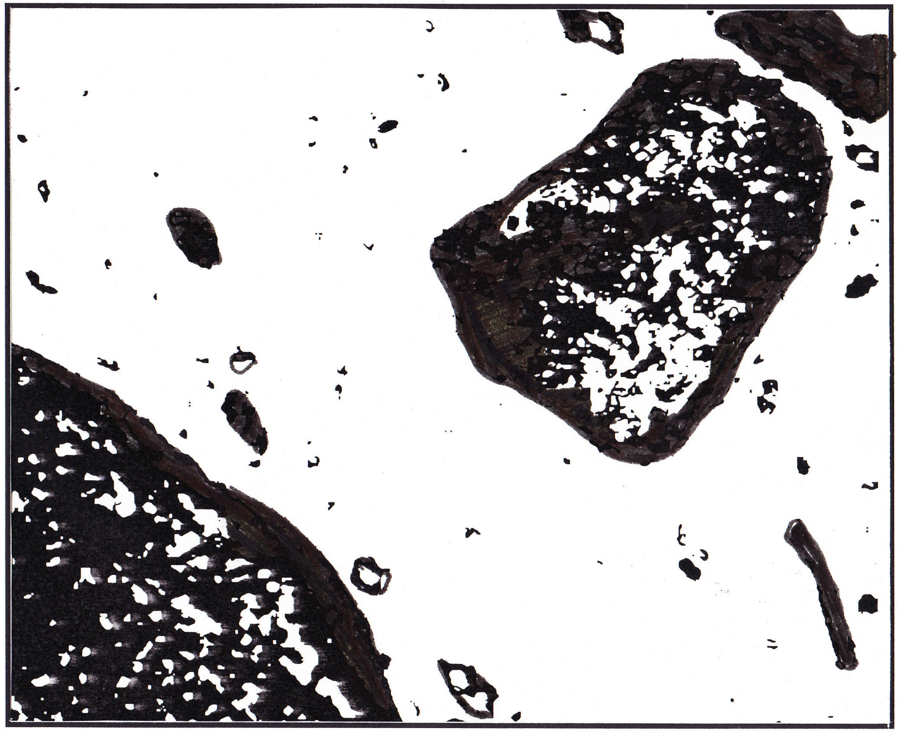

Punta de Blanco

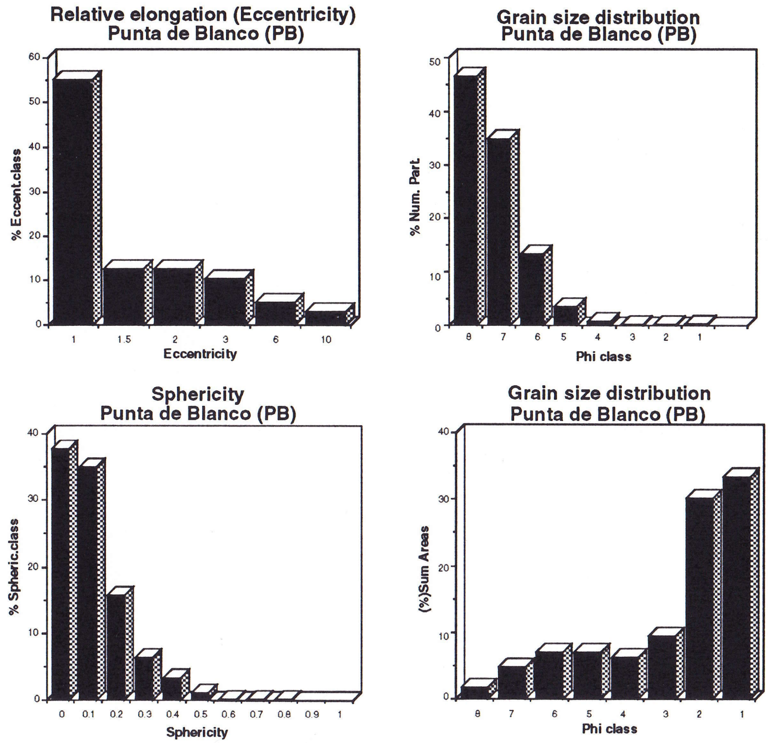


Figure 22: Organic particles morphological characterization. "Cienaga Punta de Blanco" swamp surface sample

### DISCUSSION OF THE ENVIRONMENTS

#### The fresh shallow water lake environment

##### Ayapel section

The fresh water shallow lake lithology is characterized by silty light gray clays, with oxidation spots. Plant remains might be present.

From the point of view of the palynological residue the flora consists mainly of composite, grass pollen and fungal material, being in general sporomorphs more abundant than fungal remains. Fresh water algae (including *Botryococcus* ) are always present in those assemblages, while pollen grains from water plants can be also present but in minor concentrations.

The amount of organic matter is higher close to the swamp bottom surface, at 0.25 m depth: 0.2% value was registered, but decreases sharply below that depth were values does not reach 0.1% (figure 4).

The type of organic matter preserved in this environment varies with burial history.

- In the first 0.25 m the assemblage is characterized by abundant light coloured plant remains as epidermal/cuticular and other plant remains, present but in lower concentrations are opaque amorphous materials, palynomorphs, animal, fungal and woody remains. The finely dispersed organic matter is the most abundant type.
- Downwards a "transition zone" has been observed with a strong decrease on the light coloured tissues (epidermal/cuticular and other plant remains) and a relative increase of amorphous heterogeneous and amorphous finely dispersed. Minor amounts of amorphous opaque and palynomorphs are present.
- Below 0.45 m the assemblages are characterized by amorphous opaque materials. Palynomorphs, other plant tissues, woody, fungal remains and other amorphous materials are present but in very low concentrations.

The morphological characteristics of the particles also varies with burial and follows closely the same patterns recognized for the changes in composition.

- Close to the surface (0.25 m) the assemblage is relatively coarser grained with about 80% of the total area covered by particles in the ø3 to ø1 (sand medium to very fine sand) size range. Numerically more than 55% of the particles fall in the range of ø7 to ø 4 (silt size range). Clay (ø8) size materials are about 45% of the particles but they cover less than 3% of the total particles area.

About 60% of the particles belong to the equidimensional shapes, while about 15% clearly have elongate to strongly elongate shapes.

Sphericity histograms show that more that 60% of the particles are in the 0.0 and 0.1 classes.

- At 0.45 m grain size distribution are similar to the ones described below but elongation and sphericity values are slightly different .

- Below 0.75 m depth the sum of areas grain size distribution histograms are all characterized by a maximum in the ø 5 class (medium silt equivalent size).

About 50% of the particles fall in the equidimensional values, but in general between 15% to 20% fall in the clearly elongate shapes. The sphericity histograms has bimodal distribution with a main maximum at 0.1 and a minor peak at 0.5 or 0.6. (with the only exception of the "transition" sample 0.45m).

Based on the type and amount of organic matter recovered from the samples it is clearly not a favourable environment for good source rocks accumulation. On the other hand due to the fine grained sediments this type of sedimentary bodies can act as seals or barriers to oil migration.

#### The swamp environment
#### (Surface samples and Cienaga de El Medio)

Two different sets of samples swamp surface and a core, were used to characterize the swamp environment.

A group of five "surface" samples taken directly from the bottom of the swamps, were used to characterize the organic assemblages as they are deposited. And a second set of samples from a core recovered from the Cienaga de El Medio swamp were used to study the changes of organic matter within the first meters of burial history. Both sets of samples showed different assemblages. The differences and some similarities will be described below.

From the lithological point of view all samples (surface and core) are very similar: silty clays, light grey or light brown in colour, some with oxidation spots and / or root remains.

The organic assemblages as are deposited on the bottom of the swamps can be characterized as follows:

1.- The main component of the sporomorphs assemblage are the pollen grains of grass and composite, with significative amounts of psilate monolete spores. Occasionally pollen from *Althernatera*, Cyperacea, Palmae, *Croton* and *Ludvigia* (Jussiaea) may be present. In general the most abundant palynomorphs belong to the fungal group, followed by insect remains and sporomorphs (fig. 20b).

2.- The bottom surface assemblages are richer in organic matter than assemblages coming from few decimetres below the surface. Concentration of organic matter in the surface varies between 0.1% and 0.3% in contrast with and average below 0.1% observed in Cienaga El Medio (fig.4).

3.-The composition of the organic matter has to be seen from two different angles. From the absolute view point the most abundant group is the finely dispersed material. When this is removed from the graphs there is abundance of light coloured plant tissues, epidermal/cuticular materials, fungal remains, opaque amorphous materials and animal remains (fig. 21).

4.- From the point of view of the morphology of the particles, assemblages are characterized by a grain size distribution (Number of particles) with materials in the clay size range in concentrations below 47% and coverage of 2 to 3% of the total particle area.

On the other hand in the sum of areas histogram it is possible to see that materials over ø2 (from fine sand upwards) cover more than 60% of the total particle area.

The relative elongation histogram shows that 45 to 60% of the particles fall in the equidimensional category but usually not less than 20% fall in the clearly elongate forms.

Most of the sphericity values belong to the 0.0 to 0.2 classes and they can reach up to class 0.8, but with the exception of the Punta de Blanco sample, the main peak that is at the 0.1 class that exceeds the rest in more than 20%.

The changes of the organic assemblages in the swamp environment are registered in the Cienaga El Medio core (Boquillas area). Some changes are certainly due to the early transformation of organic matter with burial history (oxidation, biodegradation, etc.), other differences are possibly associated with minor changes in climate happened during the sedimentation.

Lithology is in general uniform throughout the studied part of the core and similar to the sediments from the swamps bottom. It is characterized by light gray colour sandy to silty clays and silts, with variable amounts of reddish oxidation spots. The amount of oxidation spots is specially notable in the interval between 0.45 m and 1.95 m. Some intervals have plant remains (below 2.25 m) and some have micaceous *minerals*.

The registered changes in the assemblages are the following:

1.- The palynological assemblage is comparable with the surface assemblages although the flora is dominated by grass pollen and fern spores, with composite pollen as an important component. Significant in the sporomorphs assemblage are also the pollen types from Chenopodiaceae, Cyperaceae and *Mauritia* sp. Fungal remains are at least as abundant as the sporomorphs, while the presence of fresh water algae, including *Botryococcus* is constant throughout the interval (fig. 8c).

2.- The organic matter concentration is lower than at the surface, in general below 0.1%.

3.- There is an apparent change in composition shown as decrease of light coloured tissues in the buried assemblages when compared with surface assemblages. This is probably related with an early decomposition process of the less resistant plant materials.

There are other changes observed and are obviously related with the level of oxidation, as deduced from the relative amount of oxidation spots observed in the lithology, two different types of composition of the organic matter can be recognized:

a.- Normal oxidant conditions: the material is dominated by very fine debris.

b.- High oxidant conditions: the material is dominated by amorphous opaque and "organo-mineral gels" (more than 85% of all the preserved organic matter).

This variation increase/decrease of oxidation is supposed to be related a minor change of climate occurred during medieval times and that has been previously reported from Colombia (Wijmstra, 1967) and is well know in Europe as the "Little Ice Age". Calculations made with the sedimentation rate of the area pointed grossly to the approximate same time interval (between ±1200 and ±1700 A.C.).

4.- From the point of view of the particles morphological characterization two different groups can be recognized and they are closely related with the previously described: Normal oxidant conditions assemblage, probably associated to a climate very similar to todays and high oxidant conditions assemblage related with a relatively more arid climate, due to a change in rainfall distribution over the year.

a.- Normal oxidant conditions assemblage: is characterized by small particles (clay size range, ø 8) numerically these particles are close to the 50% but they cover less than 5% of all the particle area. Organic fragments in the categories of silt grain size (ø7 to ø4) account numerically for almost 50% of the total and each category for about 10% of the total particles areas. Large particles (ø1 and ø2) are present in lower concentrations, considerable below 5%. Between 50% and 65% of particles are equidimensional, with less than 5% in the strongly elongate to filiform shapes. In all samples more than 75% of the particles fall in the sphericity range from 0.0 to 0.20, but varies up to 0.8.

b.- High oxidant conditions assemblage: is characterized in general by the numerical dominance (42-50%) of very fine silt size range (ø7) particles, that cover not less than 15% of the particles area and commonly more than 20% of it. Each category of particles between ø6 and ø3 (fine silt to very fine sand) cover between 10% and 25%. All together a relative increase in particle sizes when compared with the Normal oxidant conditions assemblage. Most of the samples show that 52% to 60% of the particles are in the equidimensional range with sphericity values falling more than 70% on the 0.0 to 0.2 classes although it varies up to 0.7.

The high degradation of the organic matter occurring within few metres of burial in a tropical fluvial sedimentary basin with the conditions of the Magdalena river plain, point to the fact that this environment clearly is not favourable for the accumulation of source rocks. On the other hand, and similar to the case of the shallow lake due to the type of fine grained sediments associated, this type of sedimentary bodies can act as seals or barriers to oil migration.

#### The flood alluvial basin environment (El Limon)

The lithology associated with this environment is the coarsest of the studied sections. It is characterized by fine to medium sands and silts of light colours with oxidation spots or bands.

This is a very difficult environment for organic matter preservation due to the high levels of oxidation, favoured by the short flooded times and the long periods of dry conditions. Energy is also a highly fluctuating in this environment.

The assemblage shows very low organic content, with some samples reaching total absence of organic matter (fig. 4c). Most samples are barren of palynomorphs and when present they consist mainly of grass pollen and fungal remains (fig 13).

Most of the dominant organic matter is either "organo-mineral gel" type or amorphous finely dispersed (fig. 14). In minor amounts might be found amorphous opaque, fungal, woody and other plant materials. The presence of the "organo-mineral gel" type is closely associated with high levels of oxidation. This has been also observed in the interval with high oxidation signs recovered from the Cienaga El Medio.

Grain size distributions range is mainly from ø8 to ø3. The sum of areas grain size distribution have a first peak at ø7 and a second peak in the coarse silt to sand grain size range (ø4 to ø2) usually much more prominent than the first one. These type of histograms reflect the high fluctuation of the particles transport media energy.

Numerically the grain size distribution histogram are dominated by particles in the range of ø8 (clay size range). More than 60% of all particles fall in the equidimensional range, and the sphericity varies from 0.0 to 0.8 but has a main peak at 0.1.

Sediments from this environment, coarse grained, although frequently poorly sorted, may have some potential as reservoirs but of very poor quality.

#### The Lagoon with strong fluvial input environment (Santa Marta)

The assemblage from the lagoon is the richest in species and specimens, and the samples are the richest in organic matter with concentration values from 0.13% to 1%. (excluding the peats).

Peat and peaty clay sediments are dominated by mangrove pollen and fungal remains while clays are dominated by pollen grains of composite, grass and water plants with *Botryococcus* is restricted to clays.

The peaty clays have an assemblage dominated by mangrove and water plants pollen together with psilate monolete spores. The peat assemblage is strongly dominated by mangrove pollen. The clays are dominated by water plants pollen, side by side are the composite, grass, mangrove pollen and psilate monolete spores. Other types associated are Jussiaea, Proteaceae, *Mauritia* sp. and *Acrostichum* sp.

Changes in organic composition do not always are correlated with changes in the lithology:
- Lower peaty clays are characterized by relatively abundant finely dispersed organic matter, amorphous opaque, fungal and palynomorphs.
- Peat is are characterized by finely dispersed organic matter together with fungal remains. Epidermal / cuticular, other plant remains and palynomorphs are also important.
- The peat and clay layer closest to the lithological change have similar organic composition with mainly amorphous material (homogeneous and heterogeneous), and a decrease in the concentration of the finely dispersed material and fungal remains.
- Clays have similar composition with the exception of variations in concentration of finely dispersed organic matter that is relatively low compared with the rest section.Some are characterized by high concentrations of finely dispersed organic matter, with large fragments of different plant remains floating in the finely grained "matrix".

Grain size sum of areas distributions show a general trend to decrease of average grain size from base to top.

The lagoon environment based on the characteristics of the organic assemblages recovered is a potential area for source rock sedimentation. The vicinity of potential reservoirs in the fluvial and coastal sediments (river channels, coastal barriers, etc.) makes this environment very interesting as an ideal model for high potential geological setting for oil exploration.

The better the knowledge of the variations within this complex environment the higher the possibilities of success on its prediction.

This environment should be a first priority target for palynofacies research and modelling.

#### COMPARISON OF THE ENVIRONMENTS

These environments differ in several palynofacies aspects (table 1 and annex 5) and can be summarized as follows:

- The palynomorph association:

The shallow lake assemblage is dominated by pollen grains of composite follow by grass pollen, water plants pollen is present in comparatively lower amounts. The assemblage is in general very similar to those recovered from the swamp environment, but in the swamp environment in general the grass pollen is dominant over the composites.

The flood basin organic assemblage is usually barren of palynomorphs, and when present they are pollen grains of grasses and fungal remains.

The lagoon is rich in species and specimens. Peats are rich in mangrove pollen and fungal remains while clays are dominated by composite, grass and water plants pollen with *Botryococcus*.

- The organic matter:

Concentration of organic matter for surface swamp samples is between 0.1% and 0.3%, while for buried shallow lake and swamps sediments is below 0.1%. The flood basin has very low organic content close to 0.0 % or in the order of 1% of any other environment. The Lagoon, although varies, is the richest in organic matter with concentration values from 0.13% to 1% (excluding the peats).

About the type of preserved organic matter, the shallow lake has mainly very dark (opaque amorphous materials), while the swamp environment has a clear dominance of finely dispersed amorphous and/ or "organo-mineral gels". The scarce material preserved in the flood basin is either "organo-mineral gel" type or amorphous opaque or finely dispersed. The lagoon shows highly variable assemblages but the finely dispersed amorphous is associated with high amounts of cuticular/epidermal, amorphous homogeneous and heterogeneous and fungal remains.

The shape of particles also differs, the shallow lake grain size distribution has always a maximum in class ø5, and a bimodal distribution in the sphericity histogram (max. at 0.1 and at 0.5/0.6) those characteristics are systematically absent in the histograms from the swamps.

In the swamp two types of organic assemblages can be differentiated depending on the amount of oxidation: low oxidation and high oxidation, but in general they are unimodal coarse skewed sum of areas histograms (see table 1).

The alluvial flood basin has sum of areas grain size distribution bimodal with a first pick at ø7 and a second pick in the coarse silt to sand grain size range (ø4 to ø2) usually much more prominent than the first one. Numerically the grain size distribution histogram are dominated by particles in the range of ø8 (clay size range).

The lagoon show a clear trend to decrease in the grain size from very coarse that peaks at ø1 and to ø2 to silt dominated assemblages that peaks at ø5 to ø6. A high proportion of particles (30% to 50%) in this environment has a tendency to elongated shapes (up to elongate tabloids).

### GENERAL REMARKS AND RECOMMENDATIONS

#### General remarks

- To characterize recent environments from a palynofacies quantitative point of view it is necessary to study not only the organic assemblages closest to the surface but also at least the changes that occur to the organic matter within the first meters of burial history.

Those are:
- Changes in composition.
- Changes in particle morphology.
- Changes in palynomorph and organic matter concentration.

- It is necessary to keep in mind that in detailed studies of recent environments the changes observed within the first meters of burial history may have over-imposed origins:
  - Decomposition of organic matter due to **biological activity** (bacterial, microbial, fungal decomposition)
  - Decomposition of organic matter due to **chemical oxidation processes** (generation of inert materials: opaque amorphous and "organo-mineral gel" types).
  - Changes in the average preservation of the organic material as changes in the amount of inert materials may be linked to minor changes of the climate that temporally generate drier or wetter conditions within the same environment.

- The origin, sedimentation and preservation of the organic assemblages is a result of a complex set of geological, climatological and biological conditions. Yet it is possible to identify a group of parameters (organic matter concentration, composition, particle morphology, palynomorphs etc.) to characterize organic assemblages coming from various environments.

#### Recommendations

- This is a preliminary study in which only one example of each environment has been studied. Regardless of all the positive evidences found here (table 1) it is dangerous to generalize results without further studies to grade the among and between variability of the characteristics of the various environments.

  It is recommended to study more examples from each one of the environments.

- From the total set of morphological measurements obtained for characterization of organic particles only a subset could be evaluated, due to the shortness of time and resources allocated to this study.

  It is recommend to complete the evaluation of the whole set in order to identify further significant parameters that can contribute to quantitative characterization of environments.

- Statistical analysis and testing of all the results should be carried out (e.g. cluster and main component analysis), in order to optimize the quantitative computerized models of the environments under study.

- From the oil exploration view point the lagoon and associated environments are very interesting but at the same time very complex. It is recommended here to develop an extended sampling strategy, in order to obtain a 3D static model of the distribution and characteristics of organic assemblages in the lower coastal to shallow marine environments. The static model can be upgraded afterwards into a dynamic model suitable for prediction.

- In order to accomplish these targets it is necessary to re-evaluate the resources allocated to this study.

| ENVIRONMENT | PALYNOMORPH ASSOCIATION | | ORGANIC MATTER | | | ENVIRONMENT |
|---|---|---|---|---|---|---|
| | Concentration | Composition | Concentration | Composition | Particle Morphology | |
| ALLUVIAL FLOOD PLAIN | 50 grains/10 µl<br>70% samples barren | Grass pollen dominated assemblage | Close to 0.0% | Dominant:<br>"organo-mineral gel" type<br>Associated:<br>"other plant remains,<br>woody, amorphous opaque | -Sum of areas bimodal ø2/ø4 and ø7. Number of particles peak at ø7.<br>-60% particles equidimensional<br>-Sphericity peak at 0.1 | ALLUVIAL FLOOD PLAIN |
| SWAMP | 20 to 260 grains / 10 µl | Grass pollen dominated assemblage, very abundant composite pollen.<br>Fungal remains in high concentrations.<br>Fresh water algae, insect and worm remains and bryophyte spores. | Surface: 0.2 to 0.3%<br>Subsurface: < 0.1% | Dominant:<br>finely dispersed amorphous<br>Associated:<br>woody, light coloured plant tissues, animal remains and palynomorphs | -Sum of areas unimodal ø2/ø3. Number of particles peakø7/ø8<br>-50-60% particles equidimensional<br>-Sphericity peak at 0.1 | SWAMP |
| SHALLOW LAKE | 20 to 900 grains / 10 µl | Composite pollen assemblage, grass pollen may be abundant.<br>Fungal remains in high concentrations.<br>Fresh water algae. | Surface: ± 0.2%<br>Subsurface: <0.1% | Dominant: finely dispersed (close surf.) and opaque amorphous (subsurface)<br>Associated: light coloured tissues(surf.)palynomorphs, woody, other plant tissues, fungal and other amorphous. | - Sum of areas peak at ø5<br>- 50% particles equidimensional<br>- Sphericity hist. bimodal peaks at 0.1 and 0.5/0.6 | SHALLOW LAKE |
| LAGOON WITH STRONG FLUVIAL INFLUENCE | 120 to 700 grains / slide | Floating/water plants and mangrove pollen assemblage. Grass and composite pollen in high concentrations.<br>Fungal remains in high concentrations<br>*Botryococcus.* | 0.13 to 1%<br>(excluding peats) | Dominant: finely dispersed amorphous (some), other plant remains, amorphous homogeneous, amorphous heterogeneous, fungal or epidermal/cuticular.<br>Associated: same. | - Sum of areas unimodal, trend to decrease in grain size: base peak at ø1/ø2, med. peak at ø3/ø5, top peak at ø5/ø6.<br>- 30 to 50% particles quasiequidimensional to elongate tabloids<br>- Sphericity peak at 0.0/0.1. | LAGOON WITH STRONG FLUVIAL INFLUENCE |

Table 1: Synthesis of the main characteristics of: alluvial flood plain, swamp, shallow lake and lagoon with strong fluvial influence environment

ANNEX 1

### METHODOLOGY FOR PALYNOLOGICAL ORGANIC MATTER CONCENTRATION MEASUREMENT

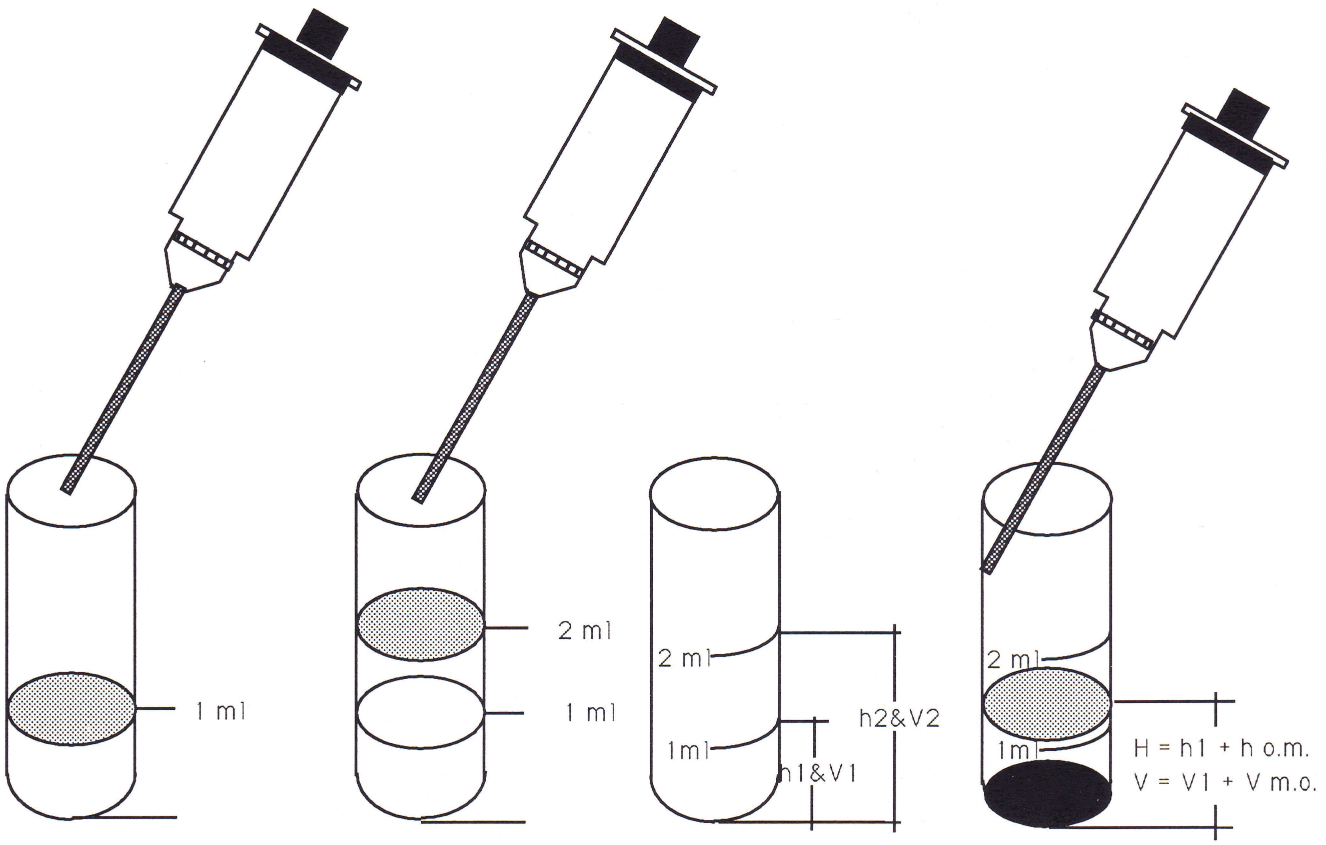


$H = h1 + h\ o.m. \Rightarrow h\ o.m. = H - h1$

$h1 \sim V1 = 1\ ml$

$h2 \sim V2 = 2\ ml$

$h2 - h1 = Hc \sim Vc = 1ml$

$V \sim V1 + V\ m.o.$

$H \sim V \Rightarrow h1 + h\ m.o. \sim V1 + V\ m.o.$

$Vc\ m.o. = (V - V1)/Vc$

$$\text{Organic matter concentration} = \frac{\text{Volume o.m.}}{\text{Volume rock}}$$

ANNEX2

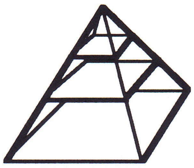

### AMSTERDAM PALYNOLOGICAL ORGANIC MATTER CLASSIFICATION

**1st Division**

**-PALYNOMORPHS**

**-STRUCTURED DEBRIS**

**-AMORPHOUS**

**-INDETERMINATE**

**2nd (and 3rd) Division**

**-PALYNOMORPHS**

- -sporomorphs
  - -pollen
  - -spores
    - -megaspores (> 200 mu)
    - -small spores (≤ 200 mu)
- -'algae'
  - -dinocysts
  - -prasinophytes
  - -chlorococcales
  - -cyanobacteria
- -acritarchs
- -zoomorphs
  - -foram linings
  - -chitinozoa
  - -tintinnids
  - -scolecodonts?
  - -rhizopods
- -fungal spores
  - -spores
  - -sclerotia
- -indeterminate
- -opaque
- -seeds

**-STRUCTURED DEBRIS**

-woody

-plant epidermis/cuticle

-plant tissue (other)

-animal

-fungal

-opaque

-indeterminate

**-AMORPHOUS**

-finely dispersed

-homogeneous

-heterogeneous

-opaque

**-INDETERMINATE**

-opaque

-other

ANNEX 3

#### MEASUREMENTS TAKEN WITH OMAS
#### for the Magdalena Project

**IMAGE ANALYSIS OF SAMPLE**

**Image input ↔ Data output**

↓

**Measurements**

##### Shape features

- AREA: sum of pixels in an object
- PERIMETER: length of the curve following the center of contour pixels
- P2A: square contour lenth divided by 4 PI times the area of the object Minimum value is 1.0 for a circle, any other figure has greater value.
- BENDING ENERGY: represents amount of energy stored in a membrane to form the given shape. Minimum value for circle
- MEAN ABSOLUTE CURVATURE: rnmac measures the total curvature in the object
- SPHERICITY: $= \frac{\text{radius inscribed circle}}{\text{radius circuncrib.circle}}$

  maximum value is 1, for a circle
- ECCENTRICITY $= \frac{\text{mayor axis of insc. ellipse}}{\text{minor axis of insc. ellipse}}$

  for a circle value is 1, any other figure has a value greater than 1.

##### Other features

- Object coordinates

#### ANNEX 4

##### RELATIVE ELONGATION SCALE

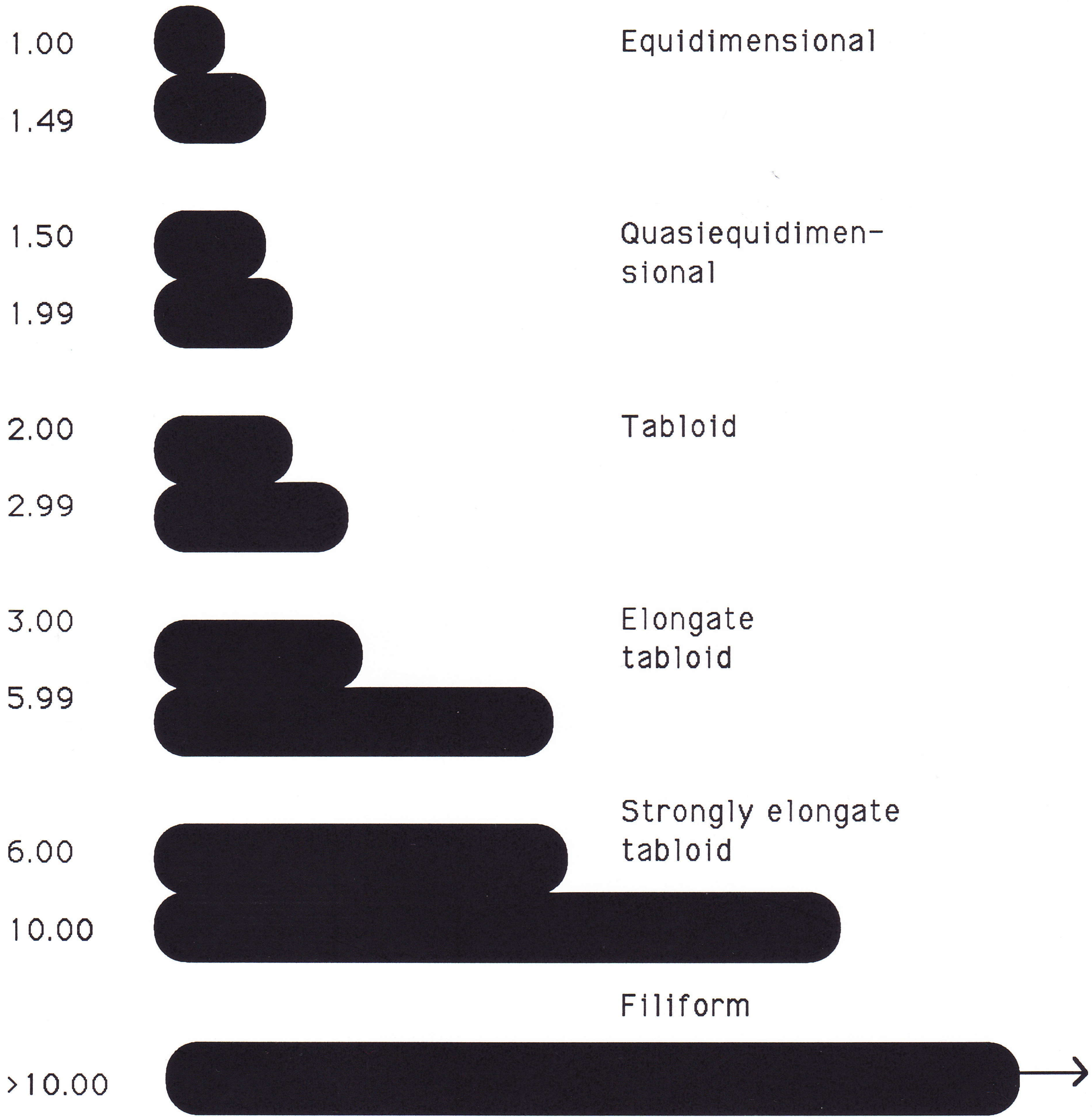

### ANNEX 5
### SUMMARY OF THE ENVIRONMENTS

#### SHALLOW LAKE ENVIRONMENT CHARACTERIZATION

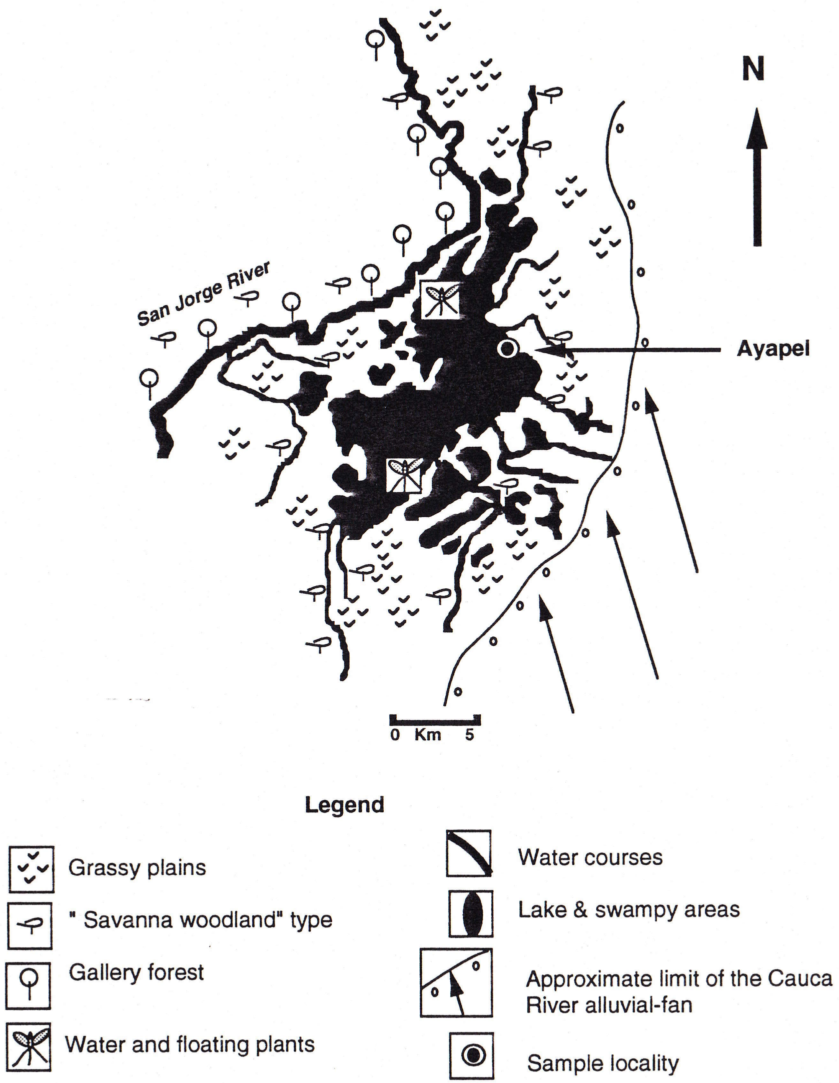


#### SHALLOW LAKE ENVIRONMENT CHARACTERIZATION

##### PALYNOFACIES

- Sporomorph concentration 20 to 900 grains /10 µl
  Average over 400 grains / 10 µl.

- Composite pollen assemblage.

- Grass pollen frequently present in very high concentrations.

- Pollen from Malphigiaceae and Protaceae may be sporadically present in high concentrations.

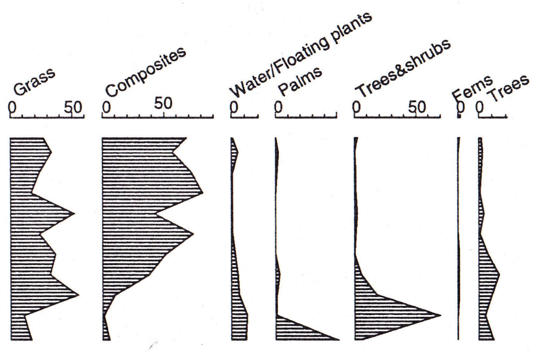


- Sporomorphs are the most abundant palynomorphs.

- Fungal remains follow sporomorphs as dominant elements.

- Fresh water algae are constantly present in the assemblages.

#### SHALLOW LAKE ENVIRONMENT CHARACTERIZATION

##### PALYNOFACIES

###### ORGANIC MATTER

**CONCENTRATION**

- Organic matter concentration is higher close to the surface, value around 0.2%.

- Below 0.45 m concentration drops sharply under 0.1%.

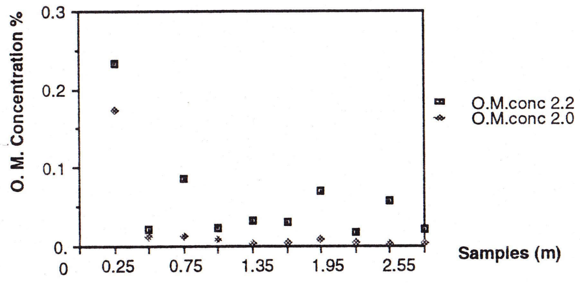



#### SHALLOW LAKE ENVIRONMENT CHARACTERIZATION

##### PALYNOFACIES

###### ORGANIC MATTER


**COMPOSITION**

- Close to the surface finely dispersed matter is the dominant type but light coloured plant tissues are very abundant.

- The transition zone (up to 0.45m) finely dispersed matter remains dominant, but decreases the light coloured material and increases amorphous heterogeneous.

- Subsurface assemblages are opaque amorphous matter dominated. In minor concentrations are: palynomorphs, woody, other plant tissues, fungal remains and other amorphous materials.

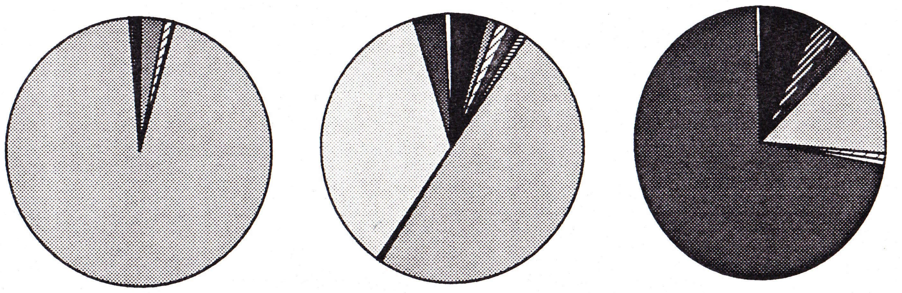

"CLOSE" SURFACE — "TRANSITION " ZONE — SUBSURFACE ASSEMBLAGE

Finely dispersed amorphous — Amorphous heterogeneous — Amorphous opaque

SHALLOW LAKE ENVIRONMENT CHARACTERIZATION

PALYNOFACIES

ORGANIC MATTER

#### MORPHOLOGY OF PARTICLES

Grain Size Distribution

Close to surface sample
- Unimodal sum of areas histogram , peak at ø1 (medium sand) coarse skewed.
- Unimodal number of particles histogram, peak at ø 8 (clay) or ø7 (v.f. silt) fine skewed.

Transition & subsurface assemblages
- All samples have in the sum of areas histogram main peak at ø5 (medium silt) rarely at ø4 (coarse silt).

Relative Elongation

- About 50% of particles belong to the equidimensional type. Between 15 and 20% have clearly elongate shapes.

Sphericity

- All histograms from subsurface assemblages are bimodal, first peak at 0.1 class and second peak at 0.5 or 0.6 class.

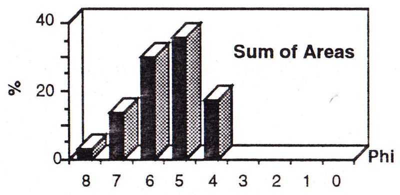


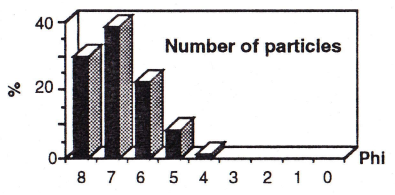


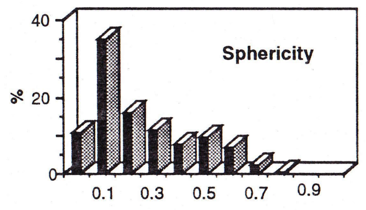


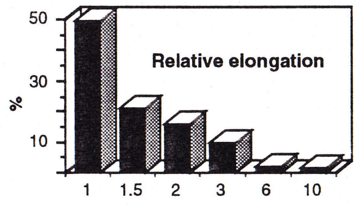

### SWAMP ENVIRONMENT CHARACTERIZATION

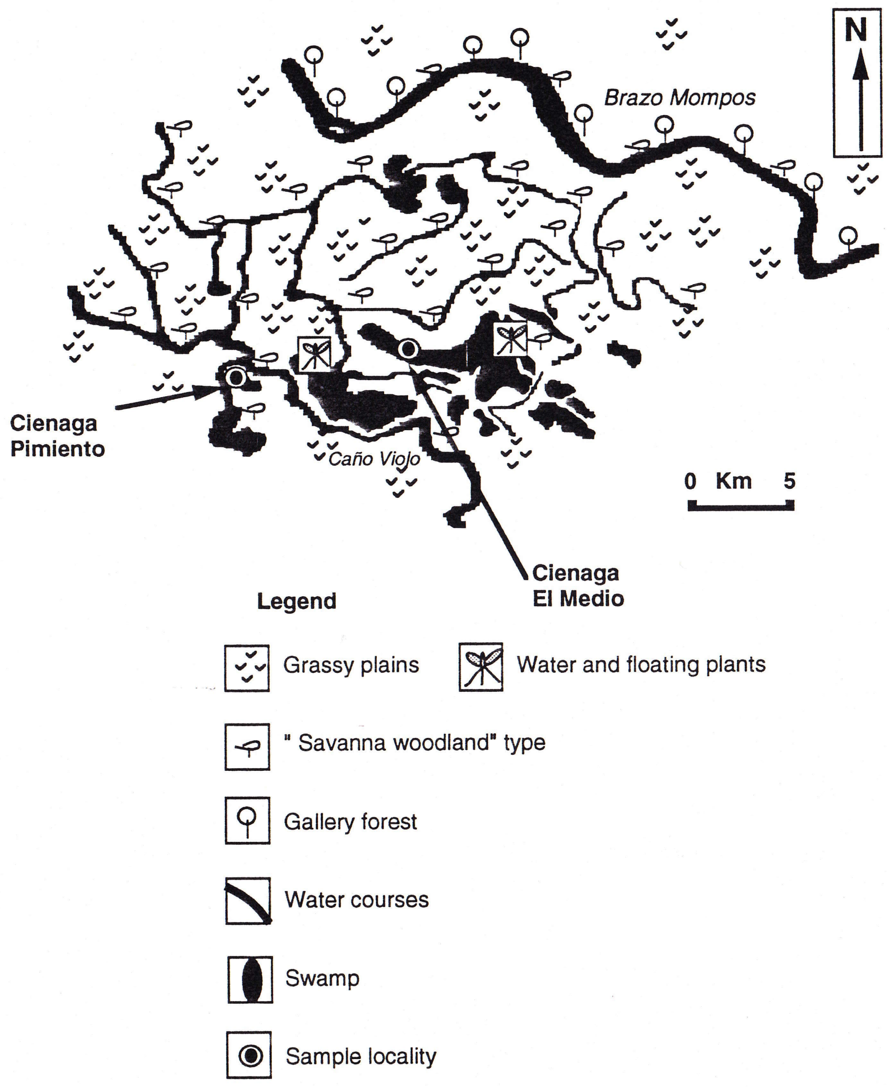


### SWAMP ENVIRONMENT CHARACTERIZATION

#### PALYNOFACIES

- Sporomorph concentration 20 to 260 grains x 10µl (surface and subsurface).

- Grass pollen assemblages.

- Composite pollen may be present in high concentrations.

- Chenopodiaceas, *Alternanthera*, Cyperaceas, Palmae, and various monelete spores important assemblage components.

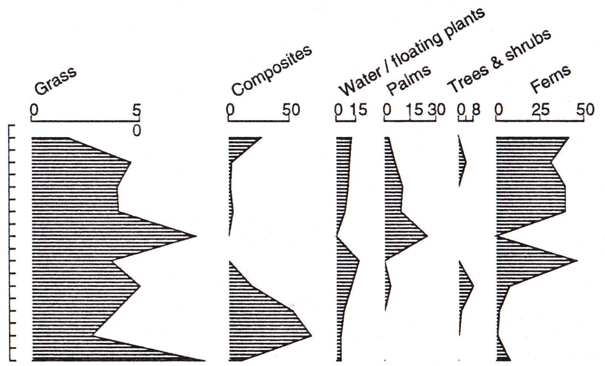


- Fungal remains same concentration as sporomorphs.

- Fresh water algae constantly present in the assemblages.

- Bryophyte spores, insect remains and conodonts may be associated.

### SWAMP ENVIRONMENT CHARACTERIZATION

#### PALYNOFACIES

##### ORGANIC MATTER

**COMPOSITION**

- At the surface light coloured plant tissues, fungal and animal remains are very abundant.

- The main organic matter component is the finely dispersed amorphous.

- Woody materials, other plant remains and palynomorphs, are always present.

- Intervals with higher oxidation show sharp increase of amorphous opaque and "organo-mineral gel" type (subsurface).

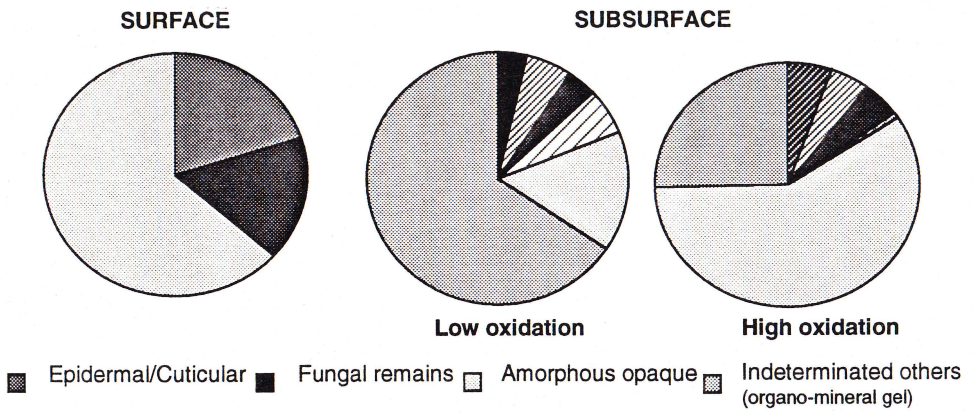


### SWAMP ENVIRONMENT CHARACTERIZATION

#### PALYNOFACIES

##### ORGANIC MATTER

**CONCENTRATION**

- Organic matter concentration at the surface between 0.2% and 0.3%.

- Organic mattter concentration below surface usually lower than 0.1%.

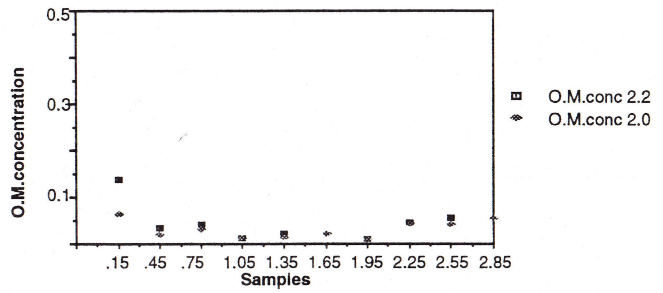

### SWAMP ENVIRONMENT CHARACTERIZATION

#### PALYNOFACIES

##### ORGANIC MATTER

**MORPHOLOGY OF PARTICLES**

Grain Size Distribution

- Sum of areas histograms unimodal, coarse skewed, main peak at ø2 and / or ø3.

- Number of particles histogram unimodal, fine skewed, main peak usually at ø7.

Relative Elongation
- 50% to 60% equidimensional particles. Less than 3% of very elongate particles.

Sphericity
- Unimodal histogram with a peak at 0.1.

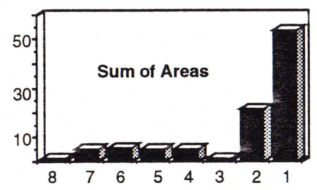


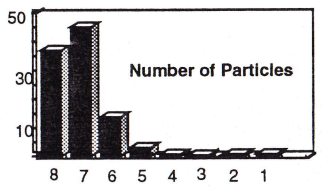


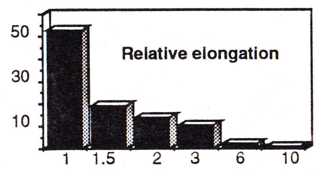


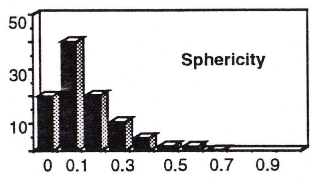

### Alluvial Flood Plain Characterization

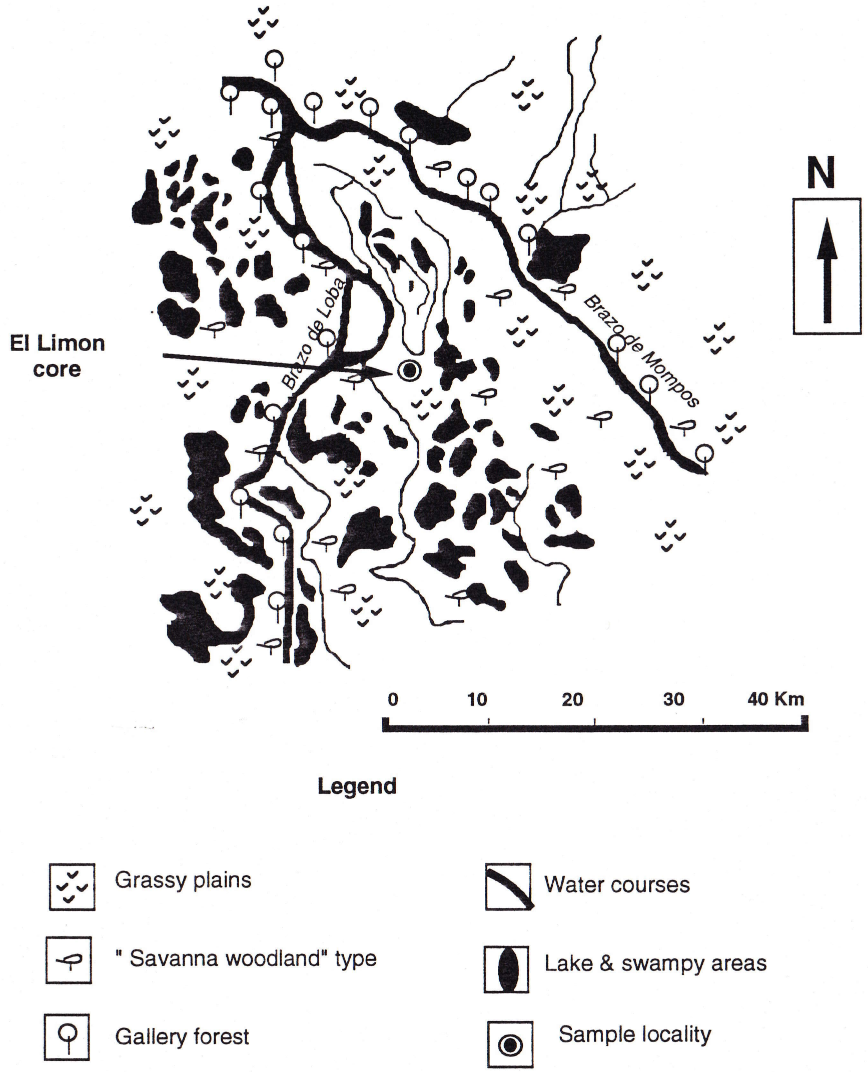


### Alluvial Flood Plain Characterization

#### PALYNOFACIES

- Sporomorphs are rarely present. Concentration under 50 grains / 10 µl

**70% of samples barren.**

- Grass pollen assemblages.

- Few composite grains may be present.

- Fungal remains may be present in same concentrations as sporomorphs.

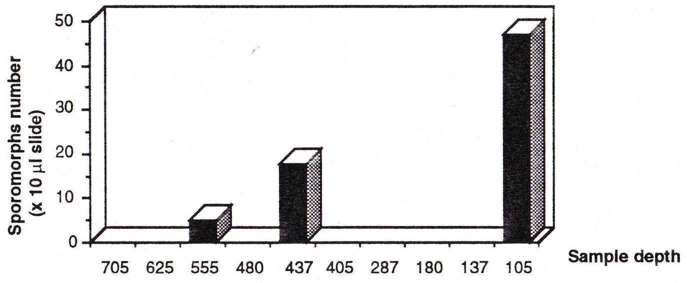

### Alluvial Flood Plain Characterization

#### PALYNOFACIES

##### ORGANIC MATTER

**COMPOSITION**

- In general dominant organic matter is the "Indeterminated others" according with the Amsterdam Palynological Organic Matter Classification".

***In an attemp to give further information, and in the specific case of the aggregates formed by mixture of almost submicroscopical particles of organic matter and minerals are refered in this work as "organo-mineral gel" type.***

- Some samples have subordinate amounts of "other plant remains", woody, and amorphous opaque materials.

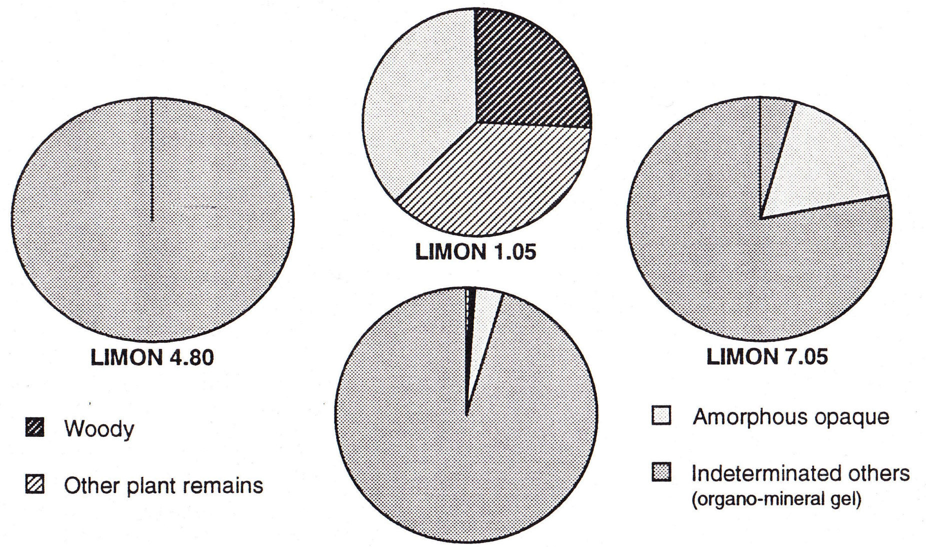


### Alluvial Flood Plain Characterization

#### PALYNOFACIES

##### ORGANIC MATTER

**CONCENTRATION**

- Organic matter in this environment is extremely low.

Concentration < 0.0%

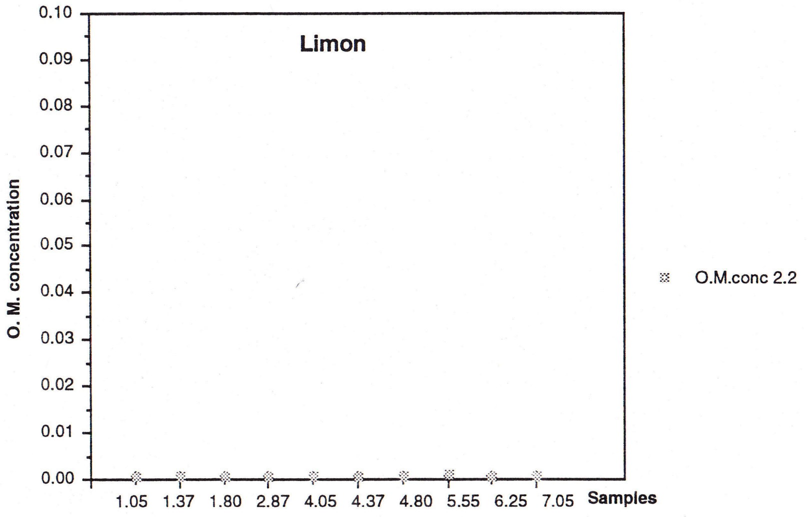

### Alluvial Flood Plain Characterization

#### PALYNOFACIES

#### ORGANIC MATTER

##### MORPHOLOGY OF PARTICLES

Grain size distribution
- Bimodal sum of areas histogram: main peak between ø2 and ø4, minor peak at ø7

  Coarse skewed.

- Unimodal number of particles histogram, main peak at ø 7

  Fine skewed.

Relative elongation
- More than 60% of particles belong to the equidimensional class.

Spericity
- Unimodal histogram with peak at 0.1 class.

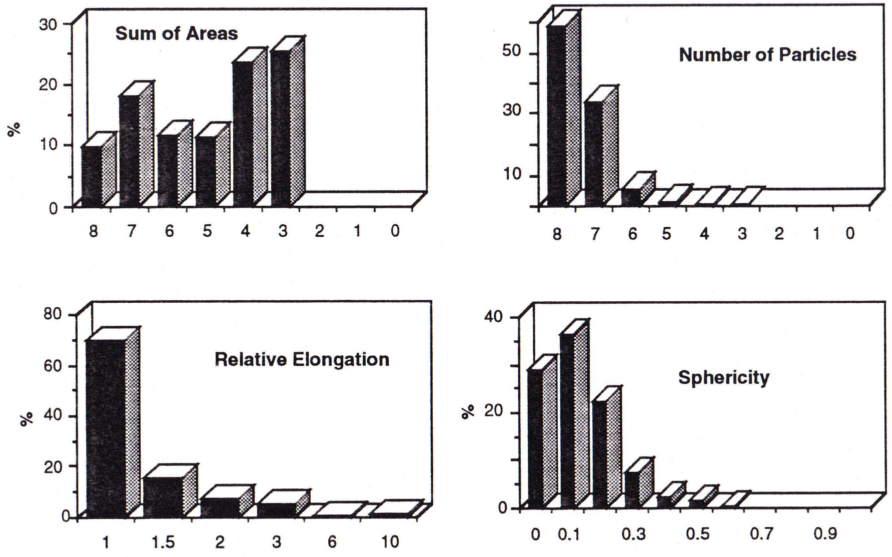

#### LAGOON WITH STRONG FLUVIAL INFLUENCE

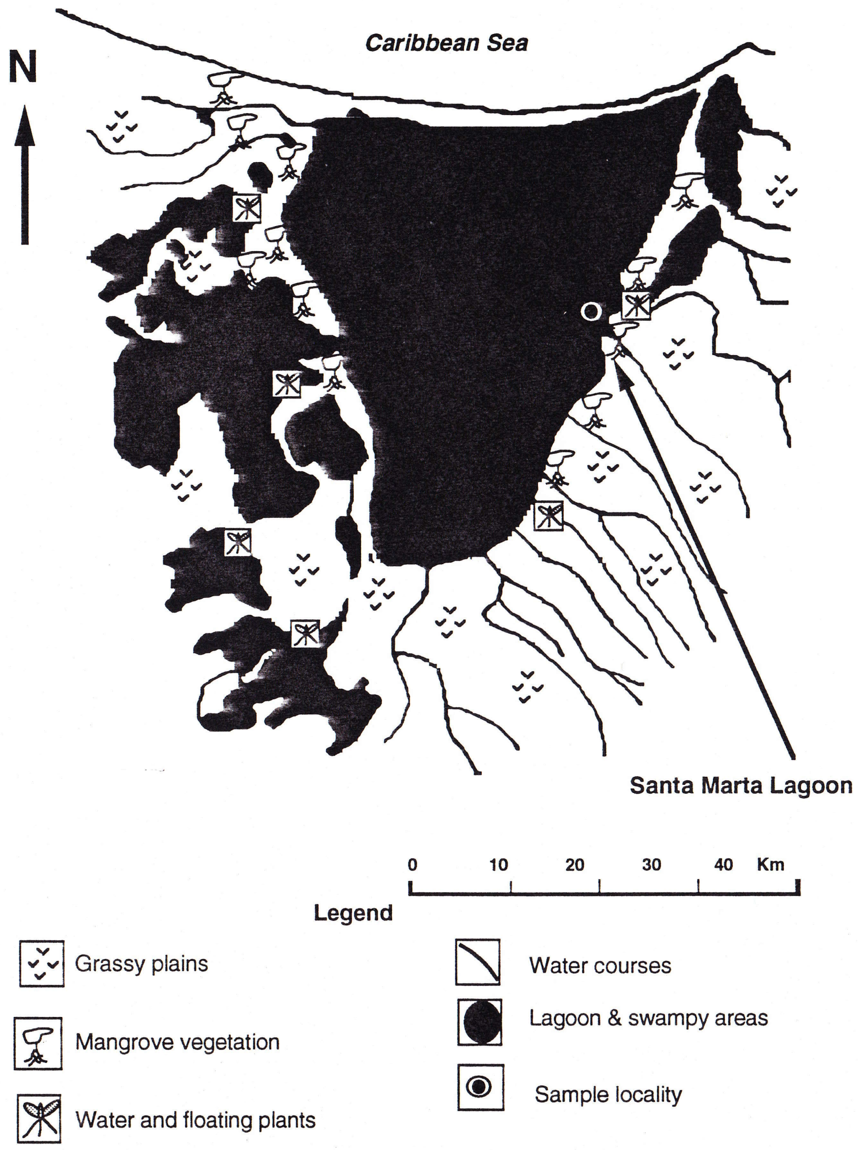


#### LAGOON WITH STRONG FLUVIAL INFLUENCE CHARACTERIZATION

##### PALYNOFACIES

- Sporomorph concentration 120 to 700 grains/slide.
- Floating / water plants and Mangrove pollen assemblages.
- Grass and composite pollen in high concentrations.
- Psilate monolete and trilete spores very abundant.

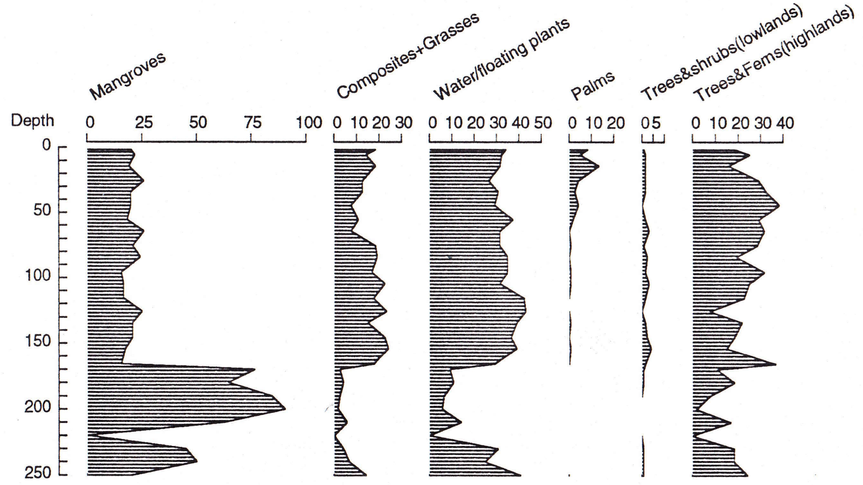


- Sporomorphs are the most abundant palynomorphs.
- Fungal remains very abundant.
- Botryococcus associated to clay lithologies. Absent in peat.

#### LAGOON WITH STRONG FLUVIAL INFLUENCE CHARACTERIZATION

##### PALYNOFACIES

##### ORGANIC MATTER

**CONCENTRATION**

- Organic matter concentration varies from 0.13% to values close to 1%.

Excluding the peats

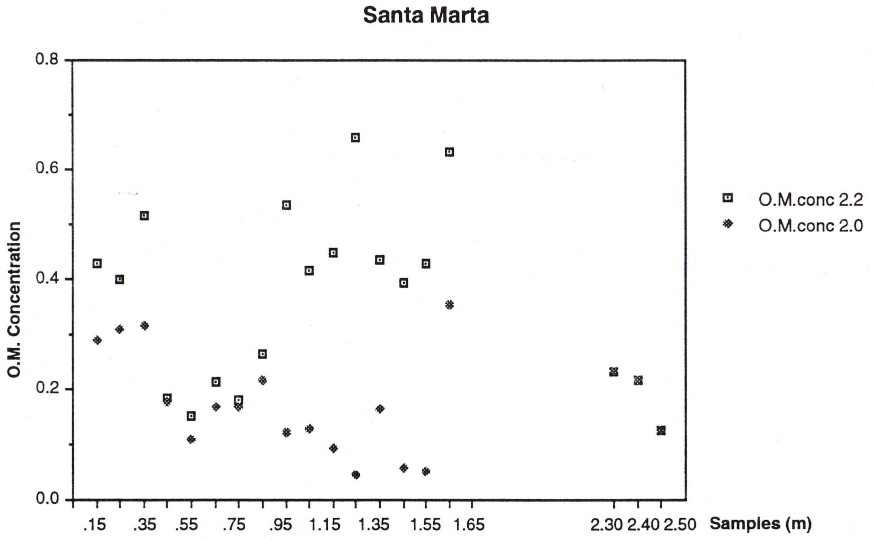


#### LAGOON WITH STRONG FLUVIAL INFLUENCE CHARACTERIZATION

##### PALYNOFACIES

##### ORGANIC MATTER

**COMPOSITION**

- Organic matter assemblages are dominated by finely dispersed organic matter in the intervals between 2.50 and 2.30m, 2.10 to 1.80 m, 1.55 to 1.35 and 0.75 to 0.05 m.

- The rest of the samples are dominated either by other plant remains, or by amorphous homogeneous, amorphous heterogeneous, fungal remains and/or epidermal/cuticular.

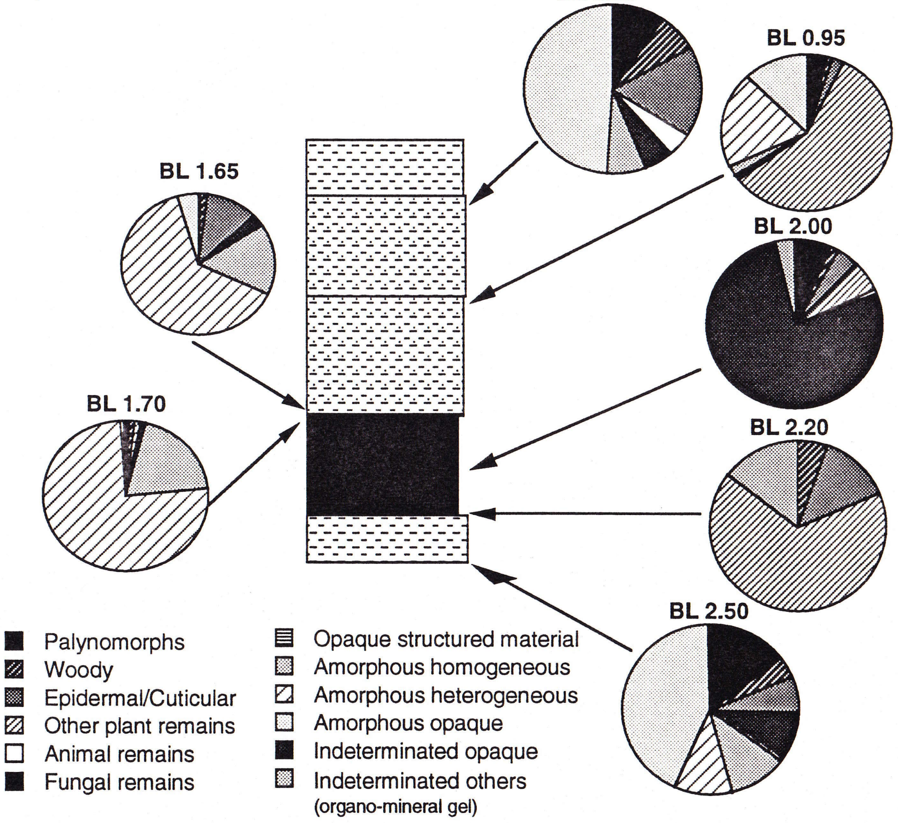

### LAGOON WITH STRONG FLUVIAL INFLUENCE CHARACTERIZATION

#### PALYNOFACIES

#### ORGANIC MATTER

##### MORPHOLOGY OF PARTICLES

Grain size distribution
- Sum of areas histograms show general trend to grain size decrease from base to top:
  2.50 to 1.70 m main peak ø1/ø2 (m. to f. sand)
  1.65 to 0.95 m peak ø3/ø5 (v.f. sand to m. silt)
  0.75 to 0.05 m peak ø5/ø6 (m. to f. silt)

- Number of particles histogram with peak at ø8, fine skewed.

Relative elongation
- High proportion of particles (30% to 50%) in the quasiequidimensional to elongate tabloids. Not less than 10% are comprised on each of these classes.

Sphericity
- Main peak at classes 0.0/0.1.

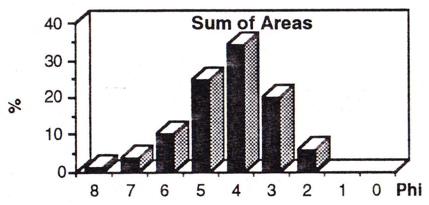


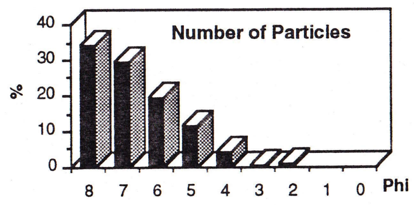


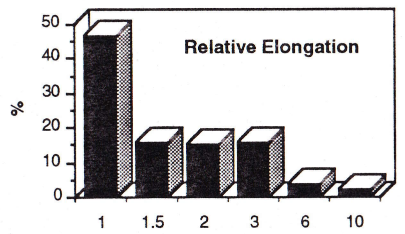


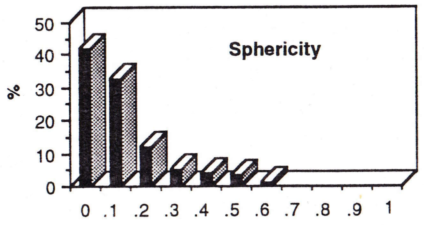
